# Distinct Th17 effector cytokines differentially promote microglial and blood-brain barrier inflammatory responses during post-infectious encephalitis

**DOI:** 10.1101/2023.03.10.532135

**Authors:** Charlotte R. Wayne, Luca Bremner, Travis E. Faust, Violeta Durán-Laforet, Nicole Ampatey, Sarah J. Ho, Philip A. Feinberg, Panos Arvanitis, Bogoljub Ciric, Chunsheng Ruan, Wassim Elyaman, Shannon L. Delaney, Wendy S. Vargas, Susan Swedo, Vilas Menon, Dorothy P. Schafer, Tyler Cutforth, Dritan Agalliu

## Abstract

Group A *Streptococcus* (GAS) infections can cause neuropsychiatric sequelae in children due to post-infectious encephalitis. Multiple GAS infections induce migration of Th17 lymphocytes from the nose into the brain, which are critical for microglial activation, blood-brain barrier (BBB) and neural circuit impairment in a mouse disease model. How endothelial cells (ECs) and microglia respond to GAS infections, and which Th17-derived cytokines are essential for these responses are unknown. Using single-cell RNA sequencing and spatial transcriptomics, we found that ECs downregulate BBB genes and microglia upregulate interferon-response, chemokine and antigen-presentation genes after GAS infections. Several microglial-derived chemokines were elevated in patient sera. Administration of a neutralizing antibody against interleukin-17A (IL-17A), but not ablation of granulocyte-macrophage colony-stimulating factor (GM-CSF) in T cells, partially rescued BBB dysfunction and microglial expression of chemokine genes. Thus, IL-17A is critical for neuropsychiatric sequelae of GAS infections and may be targeted to treat these disorders.

## INTRODUCTION

Bacterial or viral infections may trigger neuropsychiatric and cognitive dysfunction, even in the absence of brain infections^1^. Infections with *Streptococcus pyogenes*, or Group A *Streptococcus* (GAS), give rise to secondary sequelae, including those in the brain characterized by movement abnormalities (Sydenham’s chorea) and psychiatric dysfunction (Pediatric Autoimmune Neuropsychiatric Disorders Associated with *Streptococcus* infections, or PANDAS)^2^. Sydenham’s chorea (SC) is marked by uncoordinated, jerking movements and behavioral abnormalities. PANDAS symptoms have an abrupt onset, and include obsessive-compulsive behavior, verbal or motor tics, anorexia nervosa, separation anxiety, and other behavioral changes following recurrent GAS infections^3^. Although it is postulated that the secondary sequelae are due to an aberrant anti-pathogen immune response that targets the central nervous system (CNS)^2^, the mechanisms that mediate neuroinflammation, blood-brain barrier (BBB) and neuronal circuitry dysfunction are poorly understood^4, 5^. In addition, development of effective diagnosis of PANDAS has been hampered by the lack of reliable disease biomarkers^6, 7^, and there is a need for more effective treatments for severe forms of the disease^8, 9^.

Human brain imaging studies have shown increased basal ganglia volume in both SC and PANDAS patients^10, 11^, and increased microglial/astrocytic activation in the basal ganglia of PANDAS patients compared to controls^12^. Autoantibodies against basal ganglia targets have been implicated in these disorders, including anti-dopamine 2 receptor (D2R) antibodies in SC^13, 14^ and antibodies against striatal cholinergic interneurons and other neuronal targets in PANDAS^15–, 17^; nevertheless, the mechanisms driving this neuroinflammation, collectively termed post-*Streptococcal* basal ganglia encephalitis (BGE), are not well understood. Using a naturalistic GAS infection model in juvenile mice, we have previously demonstrated that multiple intranasal GAS infections cause infiltration of CD4^+^ T lymphocytes (CD4 T cells) from the nose into the anterior brain, predominantly in the olfactory bulb (OB) – the first relay center for olfaction in the brain. This is accompanied by BBB disruption and extravasation of serum immunoglobulin G (IgG) into the brain parenchyma, increased numbers of activated myeloid cells, and degradation of excitatory synapses leading to anosmia and aberrant odor-evoked neural circuitry responses^18, 19^. These studies have suggested a critical role for CD4 T cells in disease pathogenesis^4^.

T helper 17 (Th17) cells producing interleukin-17A (IL-17A) protect against extracellular pathogens, and multiple GAS infections evoke robust Th17 responses in both mice and humans^18, 20, 21^. Th17 cells have also been implicated in the pathogenesis of many autoimmune/inflammatory disorders including targeting the brain, such as multiple sclerosis (MS). Consistent with a pathogenic role for Th17 cells, we have shown that their elimination rescues, in part, BBB dysfunction and serum IgG leakage into the brain parenchyma, activation of microglia in the OB, and olfactory circuitry deficits after multiple GAS infections in the mouse model for the disease^19^. Yet, it is unknown whether specific Th17 effector cytokines play distinct roles in inflammatory responses of endothelial cells (ECs) and microglia CNS after GAS infections. Th17 cells are capable of considerable phenotypic plasticity^22, 23^, and chronic inflammation shifts conventional IL-17A-producing Th17 cells into an atypical population which produce interferon-γ (IFNγ) and granulocyte-macrophage colony-stimulating factor (GM-CSF), and express T-bet, a transcription factor typical of Th1 cells^24^. These IFNγ^+^ and GM-CSF^+^ Th17 cells, termed “pathogenic Th17s”, are required for disease development in experimental autoimmune encephalomyelitis (EAE), an MS mouse model^25, 26^, and may correlate with disease severity in human autoimmunity^27–29^. Despite being a hallmark of pathogenic Th17 cells (Th17^path^) and contributing to damage in several diseases including EAE, it is unknown whether GM-CSF is expressed by CD4 T cells that infiltrate the brain after GAS infections, and how IL-17A and GM-CSF differentially contribute to GAS-mediated neuropathology.

To address how endothelial and myeloid cells from the mouse OB and nasal-associated lymphoid tissue (NALT) respond to multiple GAS infections at the transcriptional level, we used single-cell RNA sequencing (scRNAseq) and validation with immunohistochemistry, multiplex immunoassays, flow cytometry, RNA fluorescence *in situ* hybridization (FISH) and multiplexed error-robust FISH (MERFISH). We find striking transcriptional changes in both cell populations, including downregulation of BBB genes in ECs, and elevated expression of interferon-response, chemokine and antigen-presentation genes in microglia after multiple GAS infections. Several microglial-derived chemokines are elevated in sera from PANDAS patients at the acute phase of the disease, suggesting that they could be used as biomarkers of an inflammatory state. Moreover, administration of a neutralizing antibody against IL-17A reduced microglial expression of interferon-response and chemokine genes in the mouse disease model. IL-17A blockage, but not genetic ablation of GM-CSF in T cells, partially rescued BBB dysfunction. Therefore, IL-17A may be critical for neuropsychiatric sequelae of GAS infections and can be targeted to treat PANDAS/PANS children who are refractory to standard treatments^8, 9^.

## RESULTS

### Microglia and CNS endothelial cells show major transcriptional shifts after multiple GAS infections

We have previously shown that multiple intranasal GAS infections induce infiltration of CD4 T cells from the nose into the anterior brain (predominantly in the OB), BBB disruption, microglial activation, and degradation of excitatory synapses leading to aberrant odor-evoked neural circuitry responses^18, 19^, To understand at the transcriptional level how CNS cell types respond to GAS infections, we isolated and profiled OB cells from P60 mice 18 hours after the fifth GAS infection using scRNAseq (**Figure 1A; Table S1**). Inoculation with PBS was used as a control condition. Following quality filtering and batch correction, data from OB cells were represented using dimensionality reduction with *t*-distributed stochastic neighbor embedding (*t*-SNE) and cluster identities were assigned using established cell-type markers (**Figures 1B and S1A-S1D**). A marked transcriptional shift was evident in microglia, which had the highest number of differentially expressed genes (DEGs) between PBS and GAS conditions, followed by olfactory ensheathing cells (OECs), a subtype of glia specific to the olfactory mucosa and OB (**Figures S1B, S1C, and S1F; Table S2**). Despite playing a key role in response to inflammation in other CNS diseases, astrocyte clusters showed relatively few DEGs (**Figure S1F**; **Table S2**).

**Figure 1:**
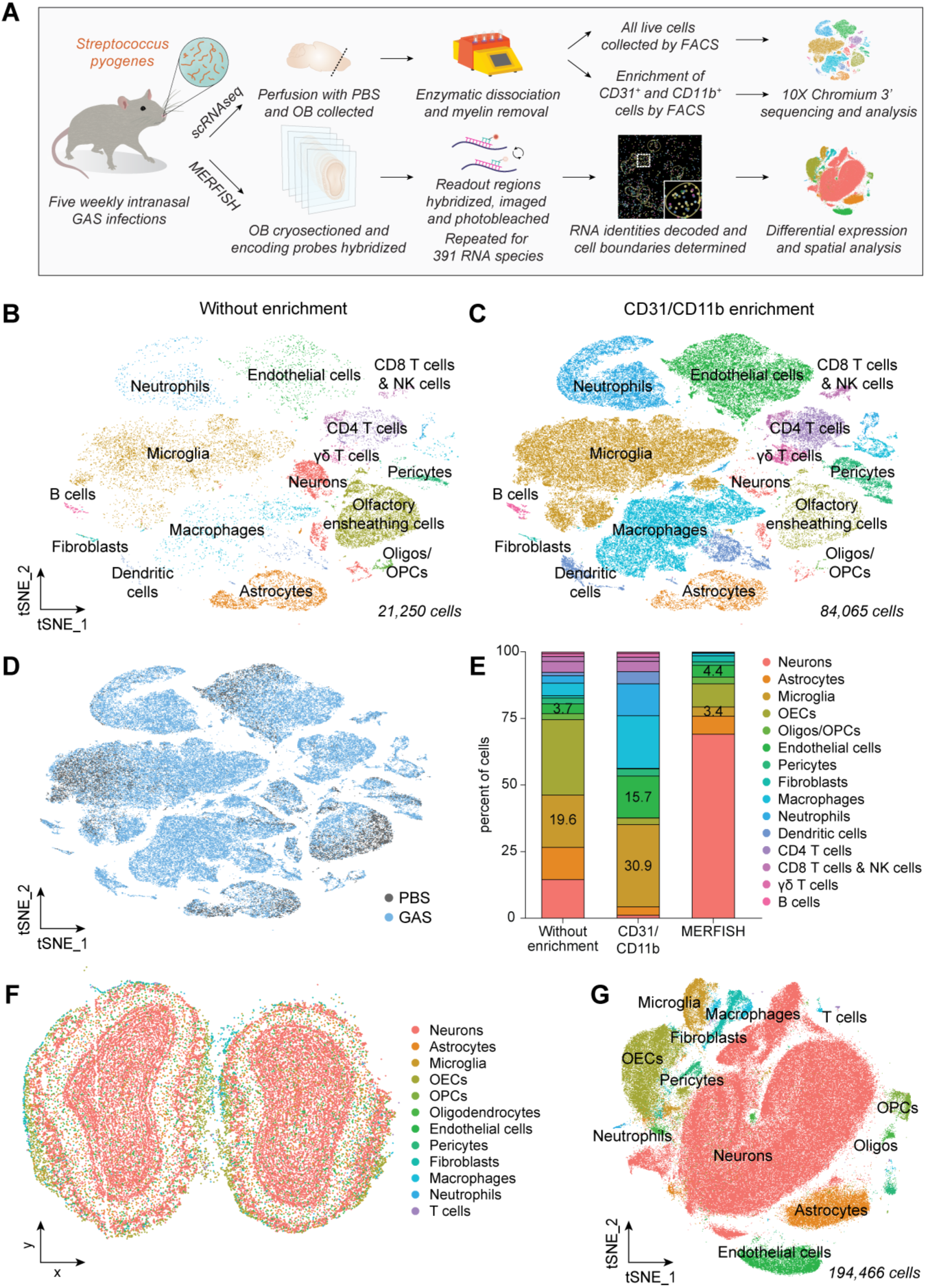
Microglia and CNS endothelial cells display major transcriptional shifts after multiple GAS infections. (**A**) Schematic diagram of the workflow for single-cell RNA sequencing (scRNAseq) and MERFISH experiments. (**B-C**) Dimensionality reduction plot of scRNAseq from the olfactory bulbs (OBs) of GAS-infected and PBS control mice without cell type-specific enrichment (**B**), and with enrichment by sorting for CD31^+^ (endothelial) and CD11b^+^ (myeloid) cells (**C**). (**D**) Dimensionality reduction plot of sample identity for OB cell types isolated from PBS (gray) and GAS (blue) conditions. (**E**) Cell type abundance from OBs from scRNAseq (with and without enrichment) and MERFISH experiments. The labeled bars represent the fraction of endothelial cells (ECs, green) and microglia (gold) among all isolated cells. (**F**) Representative MERFISH coordinate plot and (**G**) *t*-SNE plot for all MERFISH samples. See also **Figure S1**.

Since multiple GAS infections trigger BBB disruption and microglial activation in the mouse disease model^18, 19^, we focused on the transcriptional shifts in ECs and microglia by enriching for CD31^+^ and CD11b^+^ cells, respectively, using fluorescence-activated cell sorting (FACS) (**Figure S1E**). Microglia, macrophages and ECs were present in greater abundance in FACS-enriched runs (**Figures 1E**), and showed striking transcriptional shifts in gene expression between PBS and GAS conditions (**Figures 1C, 1D, and S1G; Table S2**).

In addition to scRNAseq, we performed spatial transcriptomics to gain insight into the regional distribution of cell populations identified by scRNAseq after GAS infections. We used MERFISH^30^ on mouse OB sections and probed for 391 genes predominantly expressed by microglia and ECs given their large transcriptional shifts by scRNAseq (**Figure 1A;** probes listed in methods). Cluster identities were assigned using both cell-type markers and spatial localization (**Figure S1H-L**). Since MERFISH does not enrich for any cell type, it better captured the abundance and heterogeneity of OB neuronal subtypes (**Figures 1E-1G and S1K**).

### CNS endothelial cells upregulate inflammatory signatures and downregulate BBB-specific transcripts after GAS infections

To characterize endothelial-specific transcriptional changes after multiple GAS infections, we subsetted and re-clustered ECs from the scRNAseq data (**Figures 2A**, **S2H**), and performed gene set enrichment analysis (GSEA)^31^ on DEGs using curated gene lists (**Table S4**). There was a significant upregulation in genes related to interferon response, antigen presentation, inflammation and EC response to lipopolysaccharide (LPS)^32^, as well as a downregulation of BBB-associated genes^33^ in ECs isolated from GAS compared to PBS OBs (**Figures 2B-2F; S2A-S2G; Table S3**). CNS-specific EC genes and BBB transporters were the main clusters downregulated in the GAS condition (**Figures S2A-D; Table S3**). We validated decreased expression of two BBB-specific genes, *Itm2a* and *Itih5*^33^, by FISH (**Figure 2D and 2E**) and MERFISH (*Itm2a*; **Figures S2J and S2K**) in OB sections from both conditions. In line with increased transport of serum IgG across the leaky BBB in brains from GAS-infected mice^19^, the BBB transcytosis suppressor gene *Mfsd2a*^34^ was downregulated at both the mRNA and protein level in CNS ECs (**Figure 2F; Table S3**). Surprisingly, several genes that promote EC caveolar transport (e.g. *Cav1*, *Cav2*, *Cavin1*, *Cavin2*, *Cavin3*) were downregulated in CNS ECs from GAS-infected mice (**Figure S2C; Table S3**). Pathway analysis of scRNAseq data also revealed significant downregulation of genes related to cell junctions (e.g. *Cldn5, Ocln, Tjp2, Amot, Magi3, Cgnl1*) and the extracellular matrix (ECM), including *Col4a3*, *Lamc3*, and *Itga1* (**Figures S2A and S2E; Table S3**), critical for BBB formation and maintenance^35^. We validated decreases in EC expression of *Itga1* and *Itgb1,* two integrin receptors that interact with ECM proteins, in GAS-infected OBs using MERFISH (**Figure S2I**).

**Figure 2:**
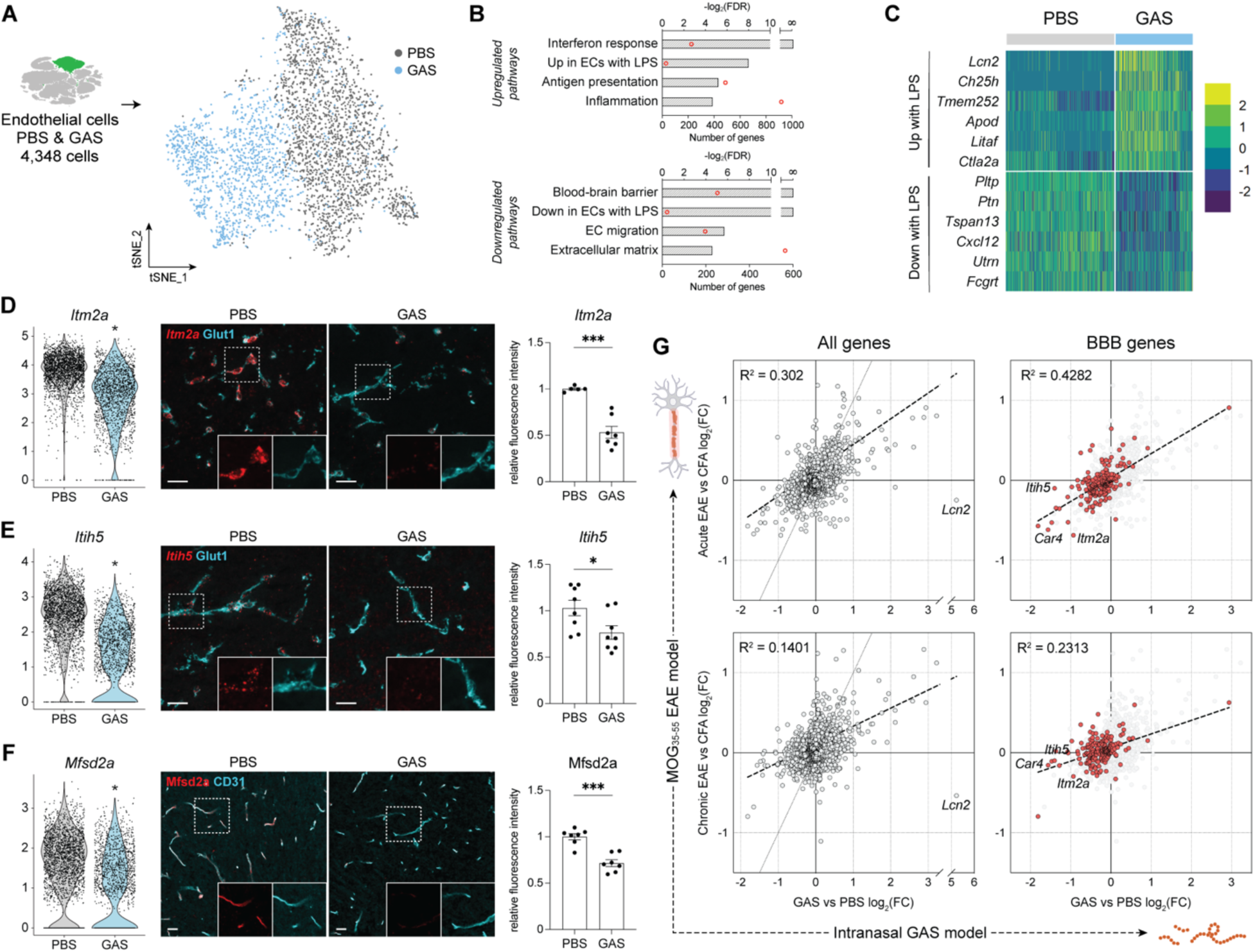
CNS endothelial cells upregulate inflammatory signatures and lose BBB marker expression following GAS infections. (**A**) *t*-SNE plot of the sample identity for ECs isolated from the OBs of PBS (gray) and GAS-infected (blue) mice. (**B**) Gene set enrichment analysis (GSEA) for genes differentially expressed in ECs following GAS infection. Bars indicate significance of enrichment (-log_2_FDR) and red dots indicate the number of genes significantly enriched in each gene set. The inflammatory signatures are upregulated, whereas blood-brain barrier (BBB) gene signatures are downregulated in ECs from GAS-infected mice. (**C**) Heat map of genes related to the endothelial response to systemic LPS in ECs from the OB of PBS and GAS-infected mice. (**D-F**) Expression changes for three BBB transcripts (*Itm2a*, *Itih5* and *Mfsd2a*) in ECs by scRNAseq (left). Statistical comparisons of scRNAseq by Wilcoxon Rank Sum test (* p-adj < 0.05). Representative images (center) and quantification (right) of expression changes in the BBB transcripts by fluorescence *in situ* RNA hybridization (FISH) for the mRNA probes (red) combined with immunofluorescence staining for EC marker Glut1 (**D-E**; cyan) and immunofluorescence staining for Mfsd2a (red) and EC marker CD31 (**F**). Scale bars = 25 μm. Statistical comparisons were performed with unpaired t test with Welch’s correction (* p < 0.05; n = 7-9 per condition). Error bars represent mean with SEM. (**G**) On the left, correlation of log_2_ fold changes in CNS EC genes identified from GAS versus PBS (x-axis) with those identified from either acute experimental autoimmune encephalomyelitis (EAE) versus Complete Freund Adjuvant (CFA) control (top panels, y-axis) and chronic EAE versus CFA (bottom paneled; y-axis). Correlations on the right display only BBB-associated genes (labeled in red). The correlation coefficient is displayed in the top left corner of each plot. Gray dotted lines mark the line of identity; black dashed line indicates the line of best fit. See also **Figure S2**.

Since BBB disruption occurs in other models of neuroinflammation (e.g. EAE), we investigated how the transcriptional shifts identified by scRNAseq in OB ECs from GAS versus PBS conditions compare to those identified by scRNAseq in spinal cord ECs isolated from either acute or chronic EAE versus Complete Freund Adjuvant controls (CFA)^36^. The correlation analysis performed on all EC genes revealed that log_2_ fold changes seen in ECs between GAS and PBS correlated more closely with acute (r^2^ = 0.302), than chronic (r^2^ = 0.1401) EAE (**Figure 2G**). The correlation coefficient was higher when we focused only on BBB-associated genes in both acute (r^2^ = 0.4282) and chronic EAE (r^2^ = 0.2313) (**Figure 2G)**. These findings suggest the presence of a core EC transcriptional response module to neuroinflammation, regardless of the trigger. In contrast, however, the inflammatory mediator lipocalin 2 (*Lcn2*) was the most highly upregulated gene in GAS ECs – as validated by MERFISH (**Figures 2C, S2J, and S2K**) – and was the most upregulated in ECs after LPS exposure^32^, but was downregulated in ECs from EAE (**Figure 2G**). Overall, the scRNAseq data confirm at the molecular level our prior functional data that GAS infections trigger BBB breakdown^18, 19^.

### Microglia increase expression of interferon-response, antigen-presentation and cytokine genes in response to GAS infections

The GSEA on DEGs by microglia in the scRNAseq data set from GAS compared to PBS OB samples (Figures 3A, S3B and S3C) showed enrichment for genes associated with interferon response, antigen-presentation, and cytokine production (Figure 3C; **Tables S3 and S4**). A large number of these genes were also elevated in ECs (**Figure S2F and S2G; Table S3**). Similar to other rodent models of neuroinflammation (e.g. EAE) and neurodegeneration (e.g. Alzheimer’s disease), microglia isolated from GAS samples showed downregulation of homeostatic genes (*Tmem119*, *Cx3cr1*, *P2ry12*, and *Gpr34*) and upregulation of disease-associated microglia (DAM) genes (*Cst7*, *Axl*, *Lpl*, *Spp1,* and *Apoe*) (**Figures S3D and 4C; Table S3**). Surprisingly, *Trem2*, which drives expression of select DAM genes in other disease models^37^, was downregulated in microglia after GAS infections (**Figure S3D; Table S3**). Expression of *Syk*, another DAM regulatory gene^38^, was also significantly decreased in microglia after GAS infections (**Figure S3D; Table S3**). It is unclear whether DAM genes are upregulated in microglia independently of *Trem2* after GAS infections. Alternatively, *Trem2* mRNA may be upregulated earlier in the course of GAS infections, or at the protein level.

**Figure 3:**
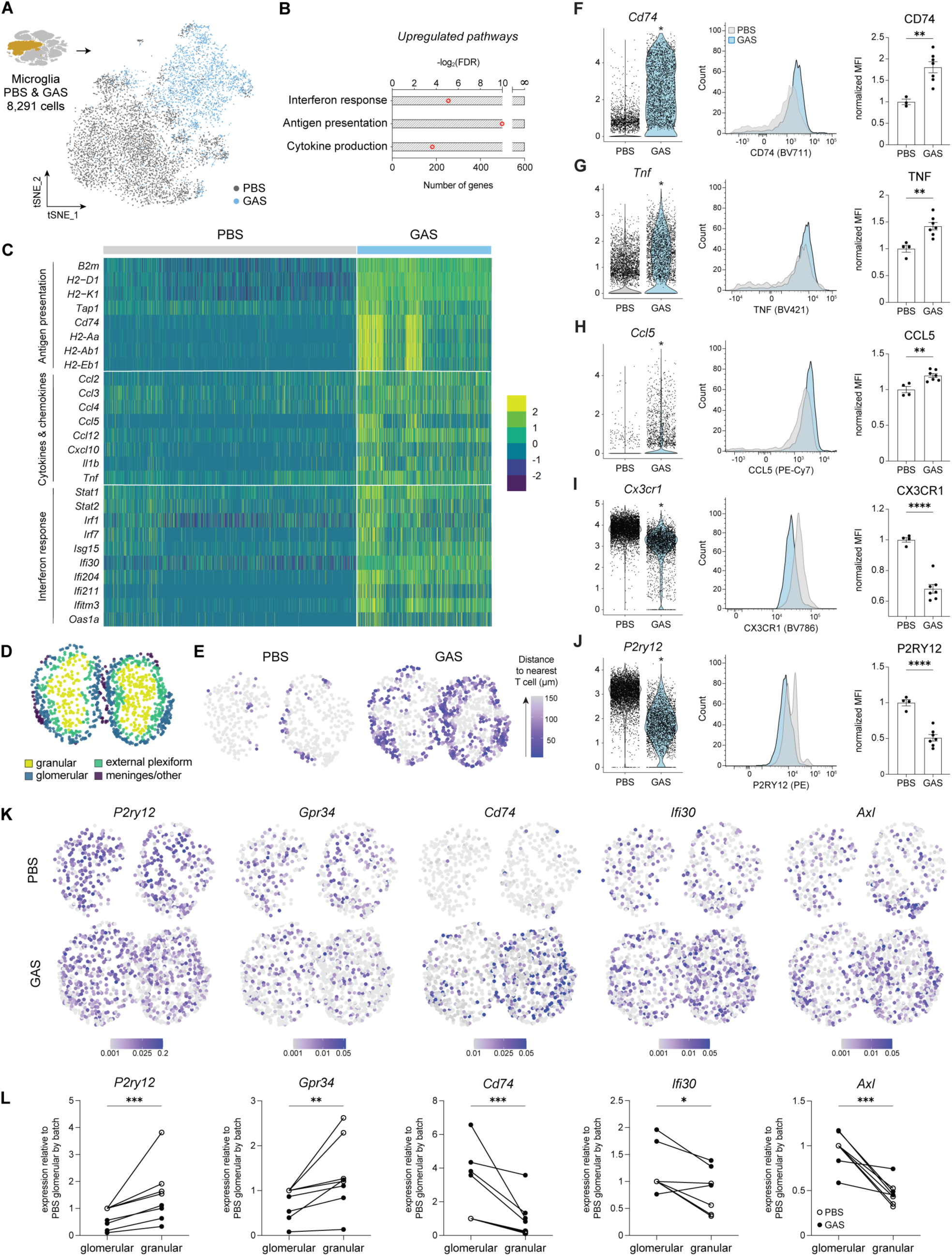
Microglia respond to recurrent GAS infections by upregulating inflammatory gene signatures. (**A**) *t*-SNE plot showing the sample identity of microglia from the OBs of PBS (gray) and GAS-infected (blue) conditions. (**B**) GSEA for genes upregulated by microglia after GAS infection. Bars indicate significance of enrichment (-log_2_FDR) and red dots indicate number of genes significantly enriched in each gene set. (**C**) Heat map of genes related to antigen presentation, cytokines and chemokines, and interferon response in PBS and GAS samples. (**D**) Representative coordinate plot of microglia color-coded for each OB layer in the MERFISH. (**E**) Representative coordinate plot of microglial distance to the nearest T cell in micrometers (µm) in each OB layer in PBS (left) and GAS-infected (right) conditions from the MERFISH analysis, colored by proximity to the nearest T cell. Microglia in the glomerular layer of the OB are closest to T cells. (**F-J**) Expression changes of *Cd74*, *Tnf*, *Ccl5*, *Cx3cr1*, and *P2ry12* in microglia by scRNAseq (left). Statistical comparisons of scRNAseq were performed with Wilcoxon Rank Sum test (* p-adj < 0.05). Representative plots (center) and quantification (right) of the normalized median fluorescence intensity (MFI) for each protein by flow cytometry. All comparisons were performed by unpaired t test (** p < 0.01; **** p < 0.0001). (**K**) Expression of *P2ry12*, *Gpr34*, *Cd74*, *Ifi30*, and *Axl* in representative spatial plots from PBS and GAS OBs in MERFISH samples (n = 4 mice per group). (**L**) Expression of homeostatic and GAS-responsive genes in microglia from the glomerular and granular layers of the OB. Data were normalized to the PBS glomerular value by batch. Comparisons by ratio t test (* p < 0.05; ** p < 0.01; **** p < 0.0001; n = 4 mice per group). See also **Figure S3**.

**Figure 4:**
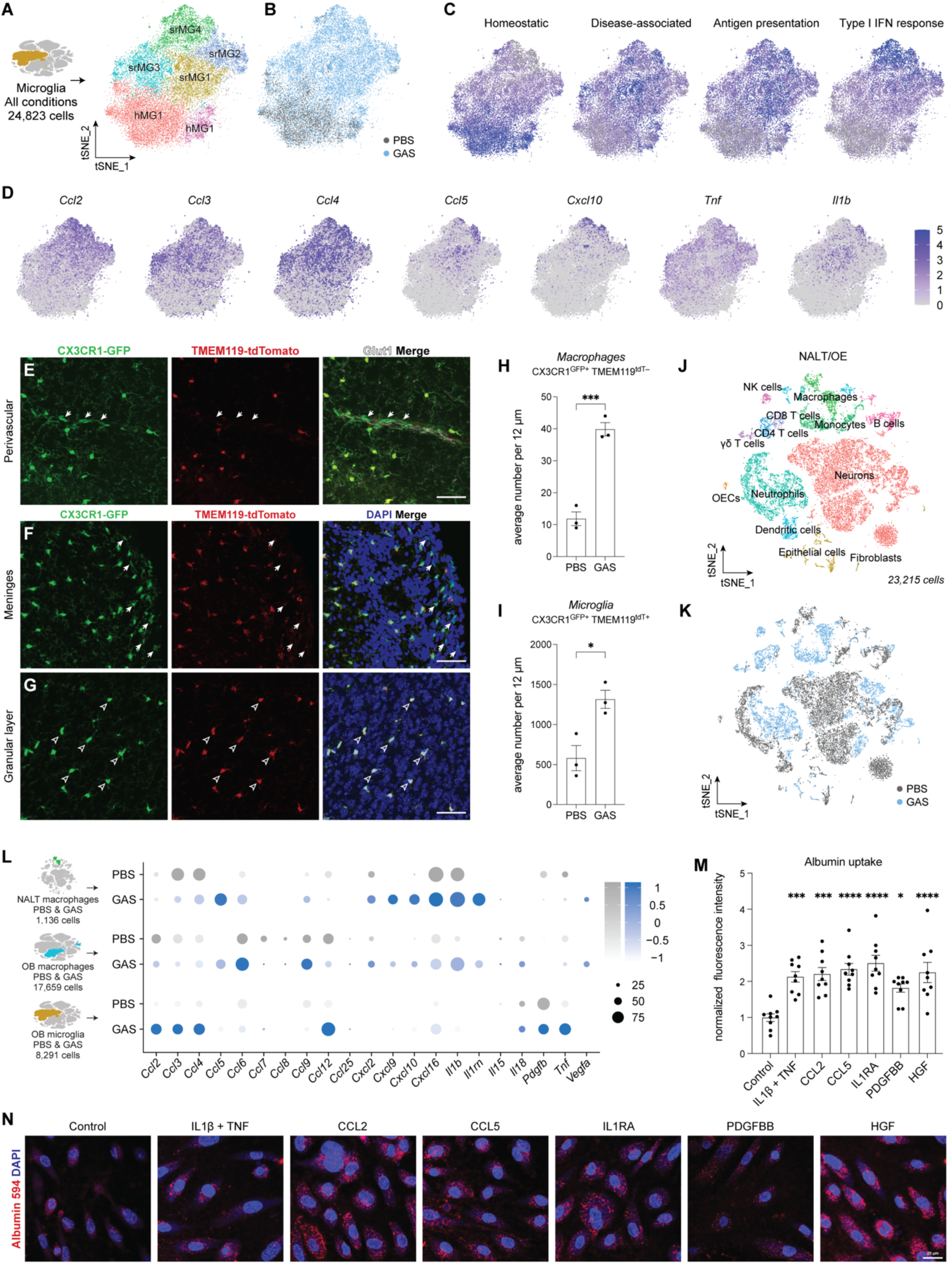
Macrophages in the NALT/OE secrete similar chemokines & cytokines to those infiltrating the OB after multiple GAS infections. (**A**) *t*-SNE plot of microglia isolated from the OBs of all conditions examined in this study reveals six major clusters: two homeostatic microglia (hMG-1 and -2) and four *Streptococcus*-responsive microglia (srMG1-4) subtypes. (**B**) *t*-SNE plot of microglia isolated from the OBs by inoculate (PBS versus GAS) shows that hMG microglia are predominantly from PBS mice (gray) and srMG1-4 from GAS-infected mice (blue). (**C**) Feature plot of the expression of selected homeostatic, disease-associated, antigen-presentation, and interferon-response genes across microglial populations. The color scale indicates average level of expression. (**D**) Feature plot of the expression of key chemokine and cytokine genes across microglial populations. The color scale indicates level of expression. (**E-G**) Representative fluorescence images of perivascular (**E**) and meningeal (**F**) macrophages, and microglia (**G**) in the OB from GAS-infected CX3CR1^GFP^/TMEM119^tdTomato^ reporter mice. White arrows (**E, F**) indicate macrophages (CX3CR1^GFP+^ TMEM119^tdTomato–^) and arrowhead outlines (**G**) indicate microglia (CX3CR1^GFP+^ TMEM119^tdTomato+^). Scale bars = 50 μm. (**H-I**) Dotted bar graph of the number of macrophages (**H**) and microglia (**I**) averaged for OB bregmas 4.5, 4.28 and 3.92 in PBS (n = 3) and GAS-infected (n = 3) CX3CR1^GFP^/TMEM119^tdTomato^ reporter mice. Comparisons were performed with unpaired t test (* p < 0.05; *** p < 0.001). (**J**) *t*-SNE plot of cell types isolated from NALT/OE based on scRNAseq data. (**H**) *t*-SNE plot of sample identity for NALT/OE-isolated cells from PBS (gray) and GAS (blue) mice. (**L**) Expression levels for select cytokines, chemokines and growth factors in NALT/OE-derived macrophages and OB-derived macrophages and microglia in PBS (gray) and GAS-infected (blue) conditions. The size of each dot indicates the percent of the population expressing each marker; the color scale indicates average level of expression. The cell population analyzed is shown in the schematic diagram on the left. (**M**) Dotted bar graph of the normalized mean fluorescence intensity of albumin-Alexa 594 localized inside primary human umbilical vein endothelial cells (HUVECs) exposed to cytokines and growth factors that are present in high concentrations in sera from PANDAS/PANS patients. Incubation with selected cytokines and growth factors for 48 hours upregulates albumin uptake in HUVEC cells. The comparisons were performed by one-way ANOVA (* p < 0.05; *** p < 0.001; **** p < 0.0001). (**N**) Representative fluorescence images of the albumin-Alexa 594 uptake in HUVECs after 48-hour incubation with the following cytokines and concentrations: IL-1β/TNF at 20 ng/mL; CCL2 at 0.5 ng/mL; CCL5 at 4 ng/mL, and IL-1RA, PDGFBB and HGF at 10 ng/mL). Scale bar = 25 μm. See also **Figure S4**.

To validate some of the DEGs obtained from scRNAseq analysis of microglia following GAS infections, we confirmed by flow cytometry microglial upregulation of antigen-presentation marker CD74 and cytokines TNF and CCL5, as well as downregulation of homeostatic receptors CX3CR1 and P2RY12 (**Figures 3F-3J**). In addition, we analyzed levels of CCL2, CCL4, CCL5, CXCL10 and TNF proteins in whole OB lysates using a multiplex immunoassay and found their levels were higher in OBs from GAS-infected mice (**Figure S3E**). At the mRNA level *Ccl2* and *Ccl3* were expressed primarily by microglia and brain macrophages, whereas *Ccl4* and *Ccl5* were also expressed by infiltrating peripheral immune cells (**Figures 3C; S3A, D**). Importantly, microglial expression of *Ccl3* and *Ccl4* was not attributable to enzymatic dissociation^39^, since neither correlate with expression of *ex vivo* activation signature genes (**Figures S3F and S3G**).

To examine the heterogeneity within the microglial response to GAS using a larger cell population, we combined microglia from all scRNAseq experiments including those where either Th17 cells were absent, or specific Th17 effector cytokines were eliminated (**Figure 4A**; see below for more details). We could identify six distinct microglial clusters. Two clusters comprising nearly all cells from the PBS condition (clusters hMG1 and hMG2) expressed homeostatic microglial genes (**Figures 4B and 4C**). Four “*Streptococcus*-responsive” clusters (srMG1-4) showed upregulation of cytokine/chemokine, disease-associated and antigen-presentation genes (**Figure 4C**). The clusters could not be distinguished by expression of single genes, but rather by gradients of up- or downregulated gene expression. For example, *Ccl3* and *Ccl4* were expressed by all srMG clusters, whereas *Ccl2* levels were the highest in srMG2 and srMG4 (**Figure 4D**). In contrast, *Ccl5* and *Il1b* were expressed by a smaller subset of srMG cells. Cluster srMG4 showed the highest expression of interferon-response genes (*Irf7*, *Isg15*, *Ifi30*, *Ifi207*, *Stat1*, *Cxcl10*; **Figure 4C and 4D**), despite no detectable upregulation of IFNα or IFNβ, either transcriptionally or by multiplex cytokine immunoassay, in the OB after the fifth infection (data not shown).

### *Streptococcus*-responsive microglia are enriched in the glomerular OB layer in close proximity to T cells

To gain insight into the spatial distribution of srMG cells, we performed spatial transcriptomics using MERFISH in the OB after intranasal inoculation with PBS or GAS. There was a distinct spatial distribution of genes upregulated in response to GAS infection. “*Streptococcus*-responsive” genes including *Cd74*, *Ifi30* and *Axl,* were expressed at higher levels in microglia located in the glomerular layer of the OB (**Figures 3D, 3K, and 3L**). In contrast, expression of homeostatic genes *P2ry12* and *Gpr34* was higher in the granular layer. A potential driver of this pattern could be the proximity of microglia to T cells, which are most abundant in the glomerular layer of the OB and meninges^18^. Microglia, which are more numerous throughout the OBs of GAS-infected mice (**Figure S3I**), had a significantly shorter distance to the nearest T cell in the glomerular and external plexiform layers of the OB compared to the granular layer as quantified by MERFISH (**Figures 3D, 3E, and S3H-S3J**).

### Brain-derived macrophages are restricted to perivascular and meningeal sites after GAS infections and resemble transcriptionally nasal-derived macrophages

Although scRNAseq data indicated that OB macrophages are enriched in GAS-infected mice (**Figures 1B-1D**), it is unclear whether macrophages infiltrate the brain parenchyma. *CX3CR1^GFP^* transgenic mice^40^ were crossed to *TMEM119^tdTomato^* reporters^41^ to distinguish between microglia (GFP^+^ tdTomato^+^) and macrophages (GFP^+^ tdTomato^−^) in tissue sections. The GFP^+^ tdTomato^−^ cells were restricted to perivascular and meningeal regions in both PBS and GAS OBs, with few to none within the brain parenchyma (**Figures 4E-4G**), although the number of macrophages and microglia was increased in the OBs of GAS compared to PBS mice (**Figures 4H and 4I**). Therefore, unlike other models of neuroinflammation, peripheral macrophages do not penetrate the brain after multiple GAS infections.

Analysis of perivascular macrophages, which express *Cd163*, *Cd207* and *Lyve1* by scRNAseq, revealed upregulation of similar pathways to those found in microglia after GAS infections, including cytokine/chemokine expression, antigen presentation and interferon response (**Figures S3A and S4B; Table S6**). *Saa3*, encoding serum amyloid A (SAA) protein 3 was the most upregulated gene in perivascular macrophages (**Figure S4B**). SAA3 is produced by myeloid cells and has been shown to sustain Th17 responses and inflammation in EAE^42^.

Since the olfactory axons may provide a route for immune cell entry into the CNS after repeated intranasal GAS infections, we performed scRNAseq of cells from the NALT/olfactory epithelium (OE) (**Figures 4J, 4K and S4A**). NALT/OE macrophages showed a similar response to brain macrophages after GAS infections at the molecular level by upregulating several cytokines and chemokines, particularly *Ccl5*, *Ccl6*, *Ccl9*, *Cxcl10* and *Il1b* (**Figure 4L**). The cytokine/chemokine expression patterns of OB macrophages resembled more closely to NALT/OE macrophages than OB microglia (**Figures 3C, 4L, and S4B**) suggesting a shared origin.

### PANDAS/PANS patients have elevated serum levels of inflammatory cytokines and growth factors expressed by microglia/macrophages

Since distinct cytokines and chemokines derived from either T cells, microglia or macrophages are highly upregulated after GAS infections in the rodent model (**Figures. 3C, 4L, S3A, S4B**), we wondered whether similar cytokines are also elevated in sera of PANDAS/PANS patients at the acute phase of the disease. We obtained serum samples from 10 PANDAS cases recruited for an IVIg clinical trial at the National Institute for Mental Health (NIMH)^43^ and 11 age-and sex-matched controls from NIMH, and recruited 13 PANDAS/PANS cases at Columbia University Irving Medical Center (CUIMC) (**Table S10**) and analyzed them for the presence of 45 serum proteins using a multiplex immunoassay (**Table S6**). Serum from PANDAS/PANS patients showed a significant elevation in 13 of 45 tested cytokines, chemokines and growth factors compared to healthy controls (**Table 1;** p-adjusted < 0.05 for 24 significant comparisons). Among the elevated PANDAS/PANS sera proteins, six (CCL2, CCL3, CCL4, CCL5, CXCL10 and TNF) were also highly upregulated by mouse microglia or macrophages after GAS infections (**Table 1**). GM-CSF, a cytokine produced by pathogenic Th17 cells (see below), was also significantly elevated in patient sera (**Table 1**). These data indicate that serum of PANDAS/PANS patients shows an inflammatory signature during the acute phase of the disease.

**Table 1:** Cytokines, chemokines and growth factors upregulated in sera from acute PANDAS/PANS patients. Forty-five proteins encompassing cytokines, chemokines and growth factors were measured by multiplex immunoassay in sera from acute PANDAS/PANS patients (n = 23) and controls (n = 11; see **Table S6**). Twenty-four proteins were significantly elevated in PANDAS/PANS patients compared to controls. Serum concentrations are provided as median (with interquartile range). Statistical comparisons were performed using the two-sample Mann-Whitney test, p-values with Bonferroni correction and adjusted p values (p-adj) for 24 comparisons (ns, p > 0.05; * p < 0.05; ** p < 0.01).

| Serum protein | Healthy controls (pg/mL) | PANDAS/PANS (pg/mL) | p-value | p-adj |
| --- | --- | --- | --- | --- |
| CCL2 | 25.5 (8.475 - 105.5) | 157 (85.5 - 253.5) | 0.0005 | * |
| CCL3 | 4 (0.525 - 10.1) | 29 (14.5 - 47.5) | <0.0001 | ** |
| CCL4 | 81 (55.5 - 90.5) | 131 (93 - 181) | 0.0012 | * |
| CCL5 | 30 (15.25 - 41) | 296.5 (91.75 - 440.75) | <0.0001 | ** |
| CXCL8 | ND | 6.9 (2.1 - 27) | 0.0011 | * |
| CXCL10 | 7.2 (5.975 - 8.525) | 59 (32 - 78) | 0.0004 | ** |
| CXCL12 | 237 (176 - 313) | 664 (385.5 - 809) | 0.0007 | * |
| EGF | 16 (0.85 - 103) | 87 (40 - 152.5) | 0.014 | ns |
| GM-CSF | ND | 16 (4.075 - 24) | <0.0001 | ** |
| HGF | 54 (31.5 - 443) | 443 (358.5 - 589) | 0.0024 | ns |
| IL-1RA | ND | 536 (110.75 - 1115.75) | 0.0156 | ns |
| IL-6 | ND | 7.85 (2.05 - 26.5) | 0.0174 | ns |
| IL-7 | 1 (0.475 - 3.4) | 5.2 (3.55 - 7.65) | 0.0004 | ** |
| IL-12A/B | ND | 2.9 (1.95 - 3.85) | <0.0001 | ** |
| IL-15 | ND | 7.9 (3.525 - 24.5) | 0.0037 | ns |
| IL-22 | ND | 8.9 (4.175 - 30.5) | 0.0183 | ns |
| IL-23 | ND | 16 (5.5 - 44.5) | 0.0244 | ns |
| KITLG | 5.2 (3.45 - 7.4) | 14 (9 - 18) | 0.0009 | * |
| LIF | ND | 28 (18 - 58.5) | 0.0129 | ns |
| PDGFB | 87 (43.5 - 586.5) | 1800 (615.5 - 2620) | 0.0004 | ** |
| PGF | 94 (77 - 116) | 188 (96 - 290) | 0.0187 | ns |
| TNF | 5.2 (4.275 - 12.1) | 18 (9.975 - 22.5) | 0.0458 | ns |
| VEGFA | 88 (31.5 - 152) | 318 (138.5 - 483) | 0.0003 | ** |
| VEGFD | ND | 22 (1.65 - 32) | 0.0253 | ns |

To understand how cytokines and growth factors present in patient sera may act on ECs, we treated human umbilical vein endothelial cells (HUVECs) with some of the cytokines that were upregulated in PANDAS/PANS patient sera (**Table 1**). After 24 hours of incubation with cytokines/growth factors, we quantified the uptake of fluorescently labeled albumin in HUVECs as a measure of caveolar-mediated uptake and transport^44, 45^, since albumin uptake is elevated in the mouse BBB after multiple GAS infections^19^. Several cytokines and growth factors, including CCL2, CCL5, IL-1RA, PDGFBB and HGF, potently upregulated transcytosis in HUVECs (**Figures 4M and 4N)**, suggesting that these elevated serum proteins sera may induce endothelial barrier breakdown in PANDAS/PANS patients during the acute phase of the disease.

### GAS-induced transcriptome shifts in microglial and CNS endothelial cells are rescued in RORγt-deficient mice

We have previously shown that elimination of Th17 cells rescues BBB dysfunction, microglial activation, and olfactory circuitry deficits in the OB after multiple GAS infections^19^. To better understand how infiltrating Th17 cells affect microglial and EC responses after GAS infections, we performed scRNAseq in mice lacking the Th17 fate-specifying transcription factor RORγt^19, 46^. ECs from GAS-infected RORγt^-/-^ mice showed a partial restoration of several BBB transcripts (e.g. *Mfsd2a*, *Itm2a* and *Itih5*) and dampened expression of transcripts related to inflammation and LPS response (e.g. *Lcn2*) compared to wild-type (WT) GAS mice by scRNAseq (**Figure S5A; Table S7**). Similarly, microglia from GAS-infected RORγt^-/-^ mice showed higher expression of homeostatic genes (*P2ry12*, *Tmem119*, *Cx3cr1*, *Gpr34*) and lower expression of DAM genes (*Cst7*, *Lpl*, *Ctsl*, *Spp1*) compared to wild-type GAS-infected microglia by scRNAseq (**Figure S5B and S5C; Table S7**). Expression of chemokines and cytokines (*Ccl2*, *Ccl3*, *Ccl4*, *Tnf*) and interferon-response (*Irf7*, *Ifitm3*, *Ifi207*, *Isg15*) genes was also reduced relative to wild-type GAS microglia (**Figure S5D and S5E; Table S7**). Intriguingly, expression of genes involved in antigen presentation (*H2-Ab1*, *H2-Eb1*, *Cd74*, *H2-D1*) was elevated further in GAS-infected RORγt^-/-^ compared to wild-type mice (**Figure S5F; Table S7**). Flow cytometry analysis confirmed significant elevation in microglial surface expression of CD74 and major histocompatibility complex (MHC) class II (I-A/I-E) proteins in GAS-infected RORγt^-/-^ compared to wild-type mice (**Figures 5A and 5B**). Th17 cell-dependent suppression of antigen-presentation genes was not limited to antigen-presenting cells. MHC class I markers *B2m*, *H2-D1* and *H2-K1* were significantly elevated in astrocytes, OECs, and to a lesser extent in neurons, ECs and microglia from GAS-infected RORγt^-/-^ mice (**Figure S5G**). A potential mechanism for this effect could be increased IFNγ in RORγt^-/-^ mice, since IL-17A negatively regulates IFNγ and Th1 cell identity^47, 48^, and IFNγ upregulates MHC gene expression in myeloid cells^49^. To determine whether the overall concentration of IFNγ is higher in RORγt^-/-^ mice, we measured IFNγ from whole OB lysates of PBS wild-type, GAS wild-type and GAS RORγt^-/-^ mice 18 hours after either the second (2i) or fifth (5i) infection. We found a significant increase in IFNγ concentration in RORγt^-/-^ compared to wild-type GAS mice at 2i, but not 5i (**Figure 5C**), indicating that IFNγ may drive expression of antigen-presentation genes earlier in disease.

**Figure 5:**
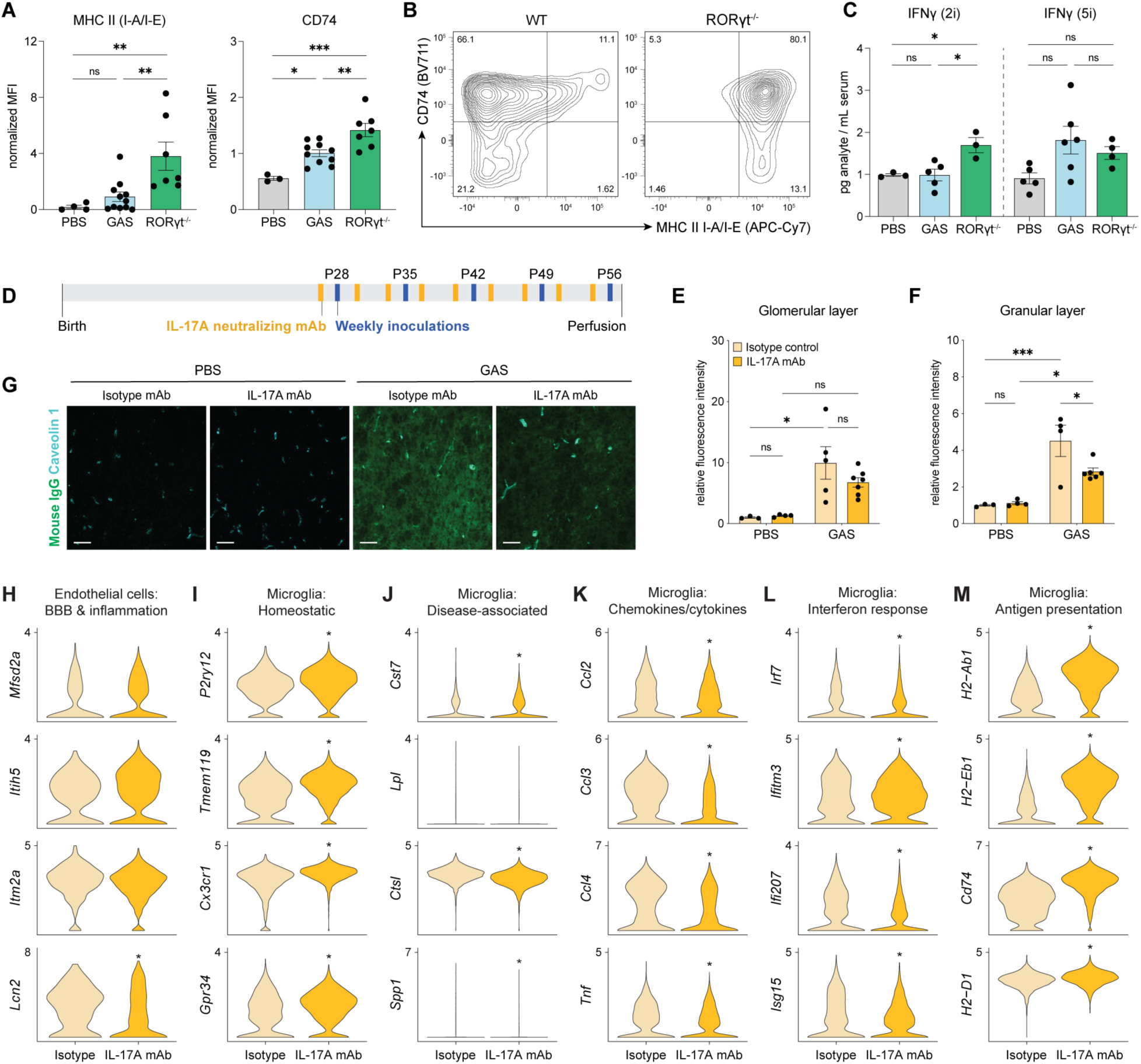
Lack of Th17 cells or inhibition of IL-17A function rescues aberrant transcriptome signatures in CNS endothelial cells and microglia associated with recurrent GAS infections. (**A**) Microglial surface expression of antigen-presentation markers by flow cytometry in wild-type PBS (gray), wild-type GAS-infected (blue) and RORγt^-/-^ GAS-infected (green) mice. Comparisons by one-way ANOVA. (ns, p > 0.05; * p < 0.05; ** p < 0.01). (**B**) Representative plots of CD74 and MHC II (I-A/I-E) expression in wild-type and RORγt^-/-^ mutant microglia after GAS infections. Contour plots represent n = 3,640 and 4,657 cells, respectively. (**C**) Quantification of IFNγ concentrations in whole OBs from wild-type PBS (gray), wild-type GAS (blue) and RORγt^-/-^ GAS (green) mice after two and five GAS infections. Comparisons by one-way ANOVA with Tukey’s multiple comparisons test (ns, p > 0.05; * p < 0.05). (**D**) Timeline of GAS infections and administration of an α-IL-17A-neutralizing antibody or isotype control. (**E-F**) Dotted bar graphs show the quantification of endogenous IgG leakage (relative fluorescence intensity) in the glomerular (**E**) and granular (**F**) layers of the OB in PBS and GAS conditions treated with either an α-IL-17A-neutralizing antibody (dark yellow) or isotype control (pale yellow). IgG intensity is significantly reduced in IL-17A mAb-treated mice relative to isotype mAb in the granular layer. Statistical analyses by two-way ANOVA with Tukey’s multiple comparison test (* p < 0.05; *** p < 0.001; n = 3-7 mice per group). (**G**) Representative images of IgG (green) leakage in the granular layer of the OB. The vasculature is marked with Caveolin 1 (cyan). (**H-M**) Violin plots of expression changes in ECs and microglia from GAS-infected mice treated with either an α-IL-17A-neutralizing antibody (dark yellow) or isotype control (pale yellow). Comparisons by Wilcoxon Rank Sum test (* p-adj < 0.05). See also **Figure S5**.

### IL-17A is required for BBB dysfunction and microglial activation *in vivo* following GAS infections

To better understand how Th17 cells contribute to neurovascular and microglial dysfunction, we focused on two inflammatory cytokines produced by Th17 cells: IL-17A and GM-CSF. We treated mice with either an IL-17A-neutralizing monoclonal antibody (mAb), or an isotype control antibody, administered twice weekly starting 24 hours before the first GAS infection to parse the contribution of IL-17A in the RORγt-dependent pathology (**Figure 5D**). IL-17A blockade did not impact CD4 T cell infiltration into the anterior brain since their number was similar between the two conditions (**Figure S5H**). However, mice treated with the IL-17A-neutralizing antibody had a significantly higher mortality due to sepsis (**Figure S5I**), reflective of a key role for IL-17A in controlling infection^50^. Importantly, IL-17A blockade partially rescued BBB leakage after GAS infections, since there was a two-fold reduction in serum IgG extravasation in the granular layer of the OB in IL-17A blocking condition compared to isotype controls (**Figure 5E-G**), an effect similar to that seen in RORγt^-/-^ mutant mice^19^. We next performed scRNAseq on CD31- and CD11b-enriched OB cells from GAS-infected mice following treatment with the IL-17A-neutralizing antibody or isotype control. Although IL-17A blockade did not rescue expression of *Mfsd2a*, *Itiih5* and *Itm2a* (**Figure 5H**), it restored expression of other BBB genes including cell-junction (e.g. *Cldn5*, *Ocln*, *Cngl1*, *Ctnna1*, *Tjp1*, *Tjp2*), and transporter genes (e.g. *Bsg, Slc7a5, Ap2b1, Abcb1a, Slco1c1*) (**Table S7**). In addition, IL-17A blockade reduced EC expression of some inflammation and LPS response genes including *Lcn2* (**Figure 5H**). Thus, IL-17A contributes to BBB dysfunction after GAS infections, both at a molecular and functional level.

As was the case for RORγt^-/-^ mice (**Figures S5B-S5F**), microglia from the IL-17A mAb-treated condition showed rescue of many homeostatic transcripts and decreased expression of DAM signature, chemokines/cytokines, and interferon-response transcripts (**Figures 5I-5L; Table S7**). However, IL-17A blockade exacerbated upregulation of antigen-presentation genes (**Figure 5M; Table S7**). Genes related to antigen presentation by MHC class I were also upregulated in other OB cell types, including OECs, astrocytes and ECs, in IL-17A mAb-treated mice compared to isotype controls (**Figure S5G**). Therefore, IL-17A promotes several microglial-related neuroinflammatory responses after GAS infections.

### GM-CSF does not drive BBB breakdown but negatively regulates microglial abundance after multiple GAS infections

GM-CSF is another cytokine downstream of RORγt in Th17 cells with the potential to drive neuroinflammation and vascular dysfunction following GAS infections. GM-CSF plays a pathogenic role in some autoimmune paradigms, such as EAE, but is protective in others^51^. In *Streptococcus pneumoniae* infections, GM-CSF is expressed by T cells only in chronic inflammation^52^. In addition, GM-CSF was among the serum proteins significantly elevated in PANDAS/PANS patients (**Table 1**). To determine if GM-CSF^+^ CD4 T cells are present in the brains of GAS-infected mice, we used flow cytometry to track CD4 T cell subsets over the course of GAS infections. The proportions of GM-CSF^+^ CD4 T cells increased with repeated GAS infections, as did the number of IFNγ^+^ Th17 (IFNγ^+^IL-17A^+^) cells in both the OB and NALT/OE (**Figures 6A, 6B and S6A-S6C**). Approximately 90% of GM-CSF^+^ CD4 T cells were either conventional Th17s or IFNγ^+^ Th17 cells, and were significantly decreased in RORγt^-/-^ OBs after repeated GAS infections (**Figures 6C and 6D**).

**Figure 6:**
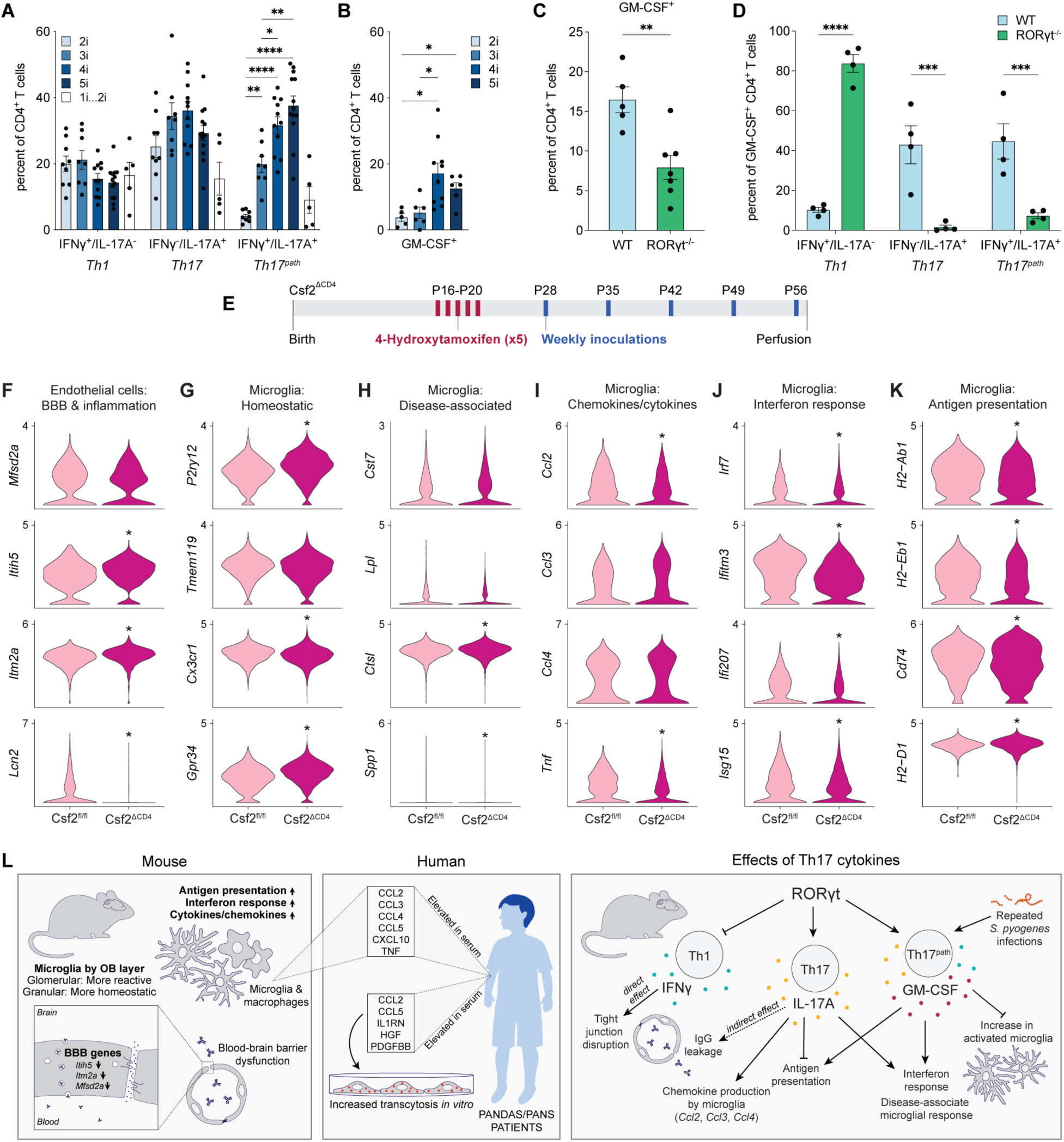
Although IFNγ^+^IL-17A^+^ and GM-CSF^+^ CD4 T cells accumulate in the brain after multiple GAS infections, GM-CSF is primarily critical to induce GAS-related microglial transcription changes. (**A**) Dotted bar graph of the progressive increase in the proportion of IFNγ^+^IL-17A^+^ CD4 T cells in the brain with increasing numbers of GAS infections. The proportion of IFNγ^+^IL-17A^+^ in mice infected twice over a five-week interval (1i…2i) was not significantly different from that seen with two consecutive infections (2i). Comparisons by one-way ANOVA with Dunnett’s T3 multiple comparisons test. (**B**) Dotted bar graph of the progressive increase in the proportion of GM-CSF^+^ CD4 T cells in the brain with increasing numbers of GAS infections. Comparisons by two-way ANOVA with Tukey’s multiple comparisons test (* p < 0.05; ** p < 0.01; *** p < 0.001; **** p < 0.0001; each dot represents one mouse). (**C**) Dotted bar graph of the proportion of GM-CSF^+^ CD4 T cells over the total CD4 T cells from the NALT/OE of wild-type (blue; n = 5) and RORγt^-/-^ (green; n = 7) GAS-infected mice. Statistical analysis by unpaired t-test (** p < 0.01). (**D**) Dotted bar graph of the proportion of GM-CSF^+^ CD4 T cells from the NALT/OE that were Th17 cells or IFNγ^+^ Th17 cells in wild-type (blue; n = 4) and RORγt^-/-^ (green; n = 4) GAS-infected mice. Approximately 90% of GM-CSF^+^ CD4 T cells in the wild-type NALT/OE were Th17 cells or IFNγ^+^ Th17 cells. Comparisons by two-way ANOVA with Tukey’s multiple comparisons test (*** p < 0.001; **** p < 0.0001). (**E**) Timeline of 4-OH-tamoxifen administration and GAS infections in Csf2^fl/fl^ and Csf2^ΔCD4^ mice. (**F-K**) Violin plots of expression changes in ECs and microglia from Csf2^ΔCD4^ mutant (maroon) and Csf2^fl/fl^ control (pink) GAS-infected mice. Comparisons by Wilcoxon Rank Sum test (* p-adj < 0.05). (**L**) A model summarizing three key findings from the study. Left: scRNAseq of mouse OB reveals that microglia upregulate antigen presentation, cytokine and chemokine, and interferon-response genes after multiple GAS infections, whereas CNS ECs lose BBB transcriptome properties. Center: A small number of inflammatory cytokines, chemokines and growth factors, produced primarily by microglia and macrophages after GAS infections, are elevated in sera from acute PANDAS/PANS patients. These cytokines upregulate transcytosis in human EC *in vitro* suggesting a direct impact on EC function. Right: Th17 cell-derived cytokines IFNγ, IL-17A and GM-CSF play distinct roles on BBB dysfunction and microglial activation in post-infectious CNS sequelae after repeated GAS infections. See also **Figure S6**.

To determine the role of GM-CSF in CNS pathology after GAS infections, we generated mice lacking GM-CSF in CD4 T cells (Csf2^ΔCD4^) by crossing *CD4-Cre^ERT2^*^53^ to *Csf2^fl/fl^* ^54^ mice and administering 4-OH-tamoxifen between P16 and P20 prior to beginning GAS infections at P28 (**Figure 6E**). Using flow cytometry, we confirmed GM-CSF knockdown in CD4 T cells isolated from the OB of GAS-infected mice after 4-OH-tamoxifen administration (**Figure S6D**). There was no significant difference in either CD4 T cell infiltration (**Figure S6F**) or serum IgG extravasation across the BBB (**Figures S6G and S6H**) and most BBB-related genes including tight junctions, transcytosis (e.g. *Mfsd2a*) or transporters were unchanged between Csf2^ΔCD4^ and Csf2^fl/fl^ mice after GAS infections (**Figure 6F**; **Table S7**). However, loss of GM-CSF in CD4 T cells rescued EC expression of some BBB-related (e.g. *Itih5*, *Itm2a*) or inflammation genes (e.g. *Lcn2*) (**Figure 6F; Table S7**).

Examination of gene expression profiles in Csf2^ΔCD4^ microglia by scRNAseq indicated a slight rescue in some homeostatic and disease-associated transcripts (**Figures 6G and SH; Table S7**), and a significant reduction in interferon-response and antigen-presentation transcripts (**Figure 6J, K; Table S7**) compared to Csf2^fl/fl^ microglia after GAS infections. Unlike IL-17A blockade, elimination of GM-CSF in CD4 T cells did not reduce microglial expression of *Ccl3* and *Ccl4*, although *Ccl2* and *Tnf* were reduced in Csf2^ΔCD4^ microglia (**Figure 6I; Table S7**). Expression of *Trem2* was even further elevated in Csf2^ΔCD4^ compared to Csf2^fl/fl^ microglia after GAS infections (**Figure S6E; Table S7**). Surprisingly, loss of GM-CSF expression by CD4 T cells caused a significant increase in Iba1^+^ CD68^+^ cell number in the glomerular layer of the OB (**Figure S6I and S6J**). This was not reflected in expression of proliferation genes since Csf2^ΔCD4^ microglia downregulated several cell cycle-related transcripts (e.g. *Top2a*, *Mcm2*, *Mcm5* and *Mki67*) relative to Csf2^fl/fl^ microglia after GAS infections by scRNAseq (**Table S6**). Overall, IL-17A and GM-CSF have distinct roles in BBB dysfunction and neuroinflammatory microglial responses after GAS infections *in vivo*.

To evaluate cell-autonomous effects of IL-17A and GM-CSF on ECs, we tested their ability to disrupt barrier properties of mouse brain capillary endothelial cells (mBECs) *in vitro*, along with IFNγ, which is produced by infiltrating Th1 and IFNγ^+^ Th17 cells after multiple GAS infections (**Figure 6A**). While treatment with positive controls IL-1β/TNF and IFNγ significantly decreased trans-endothelial electrical resistance (TEER), a readout of paracellular EC barrier integrity, neither IL-17A nor GM-CSF had any effect on the TEER (**Figures S6K and S6L**). IFNγ, IL-17A and GM-CSF treatments failed to induce the transport of fluorescently labeled albumin, a readout of transcellular transport, in mBECs in an *in vitro* transwell assay (**Figures S6M and S6N**). Since RORγt deficiency and blockade of IL-17A rescued BBB transcytosis *in vivo,* these findings suggest that IL-17A likely exerts its effects on CNS ECs indirectly.

To evaluate the effect of T cell-derived cytokines on microglia *in vitro*, we performed bulk RNA sequencing on microglia cultures from postnatal brains after 24 hour incubation with IL-17A, GM-CSF or IFNγ. There was a dramatic shift in gene expression of cultured microglia upon treatment with IFNγ, including upregulation of inflammatory and interferon-response genes (e.g. *Irf1* and *Cxcl10*) (**Figure S6O; Tables S8 and S9**) that were also identified in microglia *in vivo* after GAS infections (**Figure 3C; Table S3**). Similarly, primary microglia responded robustly to GM-CSF in culture by upregulating several genes including *Cd300f* (**Figure S6Q; Tables S8 and S9**), a gene downregulated *in vivo* in RORγt^-/-^, IL-17A mAb-treated and Csf2^ΔCD4^ microglia (**Table S7**). In contrast, minimal changes were observed in glial cell with IL-17A treatment (**Figure S6P; Table S8**), although this is difficult to interpret since cultured microglia expressed low levels of the IL-17A receptor transcript *Il17ra*^55^ (**Table S11**). Nevertheless, T cell effector cytokines induce distinct gene expression profiles in microglia responsible for neuroinflammatory changes after repeated GAS infections.

## DISCUSSION

Infections with *S. pyogenes* are a major cause of both short- and long-term morbidity in infants and adults worldwide^56^, not only due to direct primary infections causing pharyngitis, but also due to secondary sequelae including neuropsychiatric disorders^2^. Although an aberrant humoral immune response to GAS infections has been proposed to underlie secondary CNS sequelae, the pathogenesis of CNS complications of GAS infections remains elusive from a molecular standpoint. Using scRNAseq and validation with a variety of approaches, here we provide a molecular atlas of transcriptomic changes that distinct CNS cell populations undergo after multiple peripheral GAS infections^18^. Second, we show that cytokines or chemokines derived from myeloid cells are highly elevated in sera from PANDAS/PANS patients at the acute phase of the disease, and enhance transport across the endothelial barrier. These findings support the hypothesis for a neuroinflammatory origin of CNS sequelae (SC and PANDAS)^4^. Third, we demonstrate that two Th17 effector cytokines, IL-17A and GM-CSF, differentially promote BBB dysfunction and microglial expression of interferon-response and chemokine genes in a mouse model of intranasal GAS infections. Below, we elaborate on these three key findings in relation to other neuroinflammatory/neuroinfectious diseases and their relevance for the diagnosis and treatment of CNS sequelae of GAS infections.

We have employed a naturalistic rodent model of intranasal infections to explore how recurrent GAS exposures trigger BBB breakdown and neuroinflammation at a molecular level, and how these molecular findings relate to the post-*Streptococcal* human CNS disorders PANDAS and SC. Our scRNAseq analysis of more than 100,000 cells from the mouse OB revealed that ECs and microglia are among the CNS cell types most transcriptionally altered after GAS infections. The endothelial response to GAS shares commonalities with the transcriptomes of CNS ECs in other inflammation and disease models^32, 57, 58^, including our scRNAseq findings in EAE^36^, such as upregulating antigen presentation and inflammation genes, and downregulating BBB-associated transcripts^33^. This suggests the presence of a core CNS EC transcriptional response module to neuroinflammation. Several BBB-enriched transcripts such as *Itm2a*, *Itih5*^33^, *Mfsd2a*^34^, cell junction regulators (e.g. *Cldn5, Ocln, Tjp2, Amot, Magi3, Cgnl1*), and ECM proteins and receptors (e.g. *Col4a3, Lamc3, Itga1*), critical for BBB formation and maintenance^35^, were significantly downregulated after GAS infections. These molecular changes are consistent with our findings that BBB structure and function are impaired after GAS infections^18, 19^. Although increased EC transcellular transport is one of the cell biological mechanisms by which the BBB becomes permeable to large proteins (e.g. IgG), our molecular data indicate that genes promoting formation of caveolae (e.g. *Cav1, Cav2*, *Cavin1* and *Cavin2)* were downregulated after GAS infections. Decreased expression of *Mfsd2a* could explain, in part, the upregulation in EC transcellular transport since it inhibits caveolae formation^34^; however, breakdown of cell junctions or increased bulk transcytosis or macropinocytosis can also promote transport across the BBB. The gene most strongly induced in ECs after GAS infections*, Lcn2*, is involved in the innate immune response with pleiotropic roles in the CNS, and has also been shown to depend on IL-17A signaling^59, 60^, and the inflammatory cytokine response to LPS administration is exacerbated in *Lcn2*-deficient brains^61^. Therefore, Lcn2 may regulate this response by ECs in GAS infections, which will be an intriguing future direction.

Microglia also undergo major transcriptional changes after GAS infections reflected in induction of DAMs, cytokine and chemokine, antigen-presentation and interferon-response genes, and suppression of homeostatic genes. Importantly, the MERFISH analysis revealed that expression of *Streptococcus*-responsive genes is higher in microglia in the OB areas with more infiltrating T cells, suggesting that the proximity to T cells is a key factor driving the shift to disease-associated state in microglia. Surprisingly, *Trem2*, which drives expression of DAM genes in other disease models, was downregulated in microglia after GAS infections. Studies of the DAM response in neurodegenerative diseases (e.g. Alzheimer’s disease or amyotrophic lateral sclerosis) indicate that TREM2 may be required for development of “stage 2” DAM signatures, including upregulation of genes such as *Lpl*, *Cst7*, *Axl*, *Itgax* and *Spp1*^37^. It is possible that DAM genes are upregulated in GAS microglia independently of TREM2, mRNA is upregulated earlier in the course of GAS infections, or at the protein level.

Although we could identify four distinct microglial “*Streptococcus*-responsive” clusters (srMG1-4) after GAS infections by scRNAseq, the expression of distinct srMG genes was graded, rather than discrete, across the four clusters. Interferon-response genes were an exception since they were strongly elevated in the srMG4 cluster. Both type I and type II interferon signaling have been implicated in defense against GAS infections^62, 63^. T cells that infiltrate the brain after GAS infections secrete IFNγ (type II interferon); however, microglial transcription appeared skewed toward a type I IFN response, with increased expression of genes like *Irf7*, *Isg15*, *Ifit1*, *Ifit2*, *Ifit3*, *Rsad2* and *Ms4a4c*, which are preferentially induced by IFNα^64^. However, no upregulation of IFNα or -β was detected, either transcriptionally or by multiplex cytokine immunoassay in the OB (data not shown). While interferons are primarily associated with viral infection, microglial expression of interferon-response genes has also been observed in models of inflammation^65, 66^, autoimmunity^67–69^, injury^70, 71^, neurodegeneration^72–74^ and aging^75^. The extent to which interferon signaling plays a conserved role in microglia across diverse perturbations is unclear. Interferon signaling has previously been implicated in driving cell-autonomous microglial morphology, inflammation and phagocytosis, among other functions^75–77^.

The third class of genes elevated in *Streptococcus*-responsive microglia and macrophages were cytokines and chemokines including *Ccl2, Ccl3, Ccl4, Ccl5, Cxcl10, Cxcl12, Il1b,* and *Tnf*. These cytokines are also upregulated in other models of neuroinflammation and neurodegeneration, although with distinct combinatorial patterns. The microglial cytokine expression profile more closely resembles that of OB than NALT/OE macrophages; however, the proportion of microglia expressing cytokines was higher than that of OB macrophages. Upregulation of chemokines by microglia and macrophages could contribute to recruitment of peripheral immune cells, particularly T cells and infiltrating macrophages, into the CNS. For example, CCL2-CCR2 signaling is required for Th17^path^ cell recruitment to the CNS in EAE and *S. pneumoniae* intranasal infection^52^. The receptors for most microglia-derived chemokines are expressed by macrophages and T cells in both the OB and NALT/OE. Finally, myeloid-derived chemokines (e.g. CCL2, CCL5) may also act directly on ECs to increase transcytosis across the vascular barrier. In addition to T cell effector cytokines (see below), cytokines/chemokines derived from myeloid cells may impair BBB function *in vivo*.

Our scRNAseq analysis was focused primarily on gene categories that were significantly changed after GAS infections; categories or genes that change at moderate levels may play an equal or greater role in response to GAS infections and may underlie putative genetic risk factors that predispose children to develop CNS sequelae^5^. Second, our transcriptome analysis in ECs and microglia is focused at a single time point (five intranasal GAS infections) since the CNS pathology (T cell infiltration, BBB damage, microglia activation and neuronal circuit dysfunction) is quite prominent at this stage^18, 19^. However, an analysis of transcriptomics shifts in ECs, microglia or other CNS cell types with scRNAseq or MERFISH at either an earlier stage of GAS infection, or during the resolution of the last infection can provide a more comprehensive temporal atlas of transcriptome shifts in CNS cells during GAS infection that may be critical to understand disease pathogenesis or resilience to treatment. Molecular analysis of novel microglial phenotypes over the course of GAS infections, or the use of fate-mapping microglial reporters^78^, could be useful tools to address these questions.

Although the human studies have mainly focused on “pathological” antibodies, we have previously shown that Th17 cells are critical to induce BBB dysfunction, microglial activation, and olfactory circuitry deficits in the OB after multiple GAS infections^19^. The molecular transcriptome analysis of microglia from *RORγt*^-/-^ mice supports the rescue of the pathological phenotype (i.e, microglia activation) after GAS infections^19^. Our molecular analysis of microglia shows a rescue in most *Streptococcus*-responsive pathways in *RORγt*^-/-^ mice, particularly in chemokine/cytokine and interferon-response genes. Similarly, there is a partial restoration of some BBB transcripts (e.g. *Mfsd2a*, *Itm2a* and *Itih5*) and reduced expression of transcripts related to inflammation and LPS response (e.g. *Lcn2*) in *RORγt*^-/-^ compared to wild-type GAS conditions. Although the decrease in EC transcytosis seen in *RORγt*^-/-^ mice after GAS infections^19^ is not paralleled by a significant rescue of transcytosis genes (e.g. *Mfsd2a* or *Cav1*), the rescue in either cell-junction (e.g. *Cdh5*, *Jam2*, *Tjp1*), or receptor-mediated endocytosis transcripts may explain BBB functional rescue in the absence of Th17 cells. Unexpectedly, expression of genes involved in antigen presentation was elevated further in GAS-infected RORγt^-/-^ CNS cells possibly due to increased expression of IFNγ at two, but not five, GAS infections. IFNγ is known to upregulate MHC expression in myeloid cells^49^ and its production by T cells can be negatively regulated by IL-17A^47, 48^. Remarkably little is understood about the significance of antigen presentation by microglia in different neuropathologic conditions. Microglia are the main MHC II-expressing cell type in the brain^79^, but it is unclear to what extent this is a key function of microglia, rather than a vestige of shared myeloid origins. Future studies will address this important question.

Our analysis of the distinct roles that two Th17 effector cytokines, IL-17A and GM-CSF, play in post-*Streptococcal* neuropathology expands further our understanding of the role of Th17 cells in GAS-mediated CNS sequelae. Antibody blockade of the signature Th17 effector cytokine, IL-17A, in wild-type mice is sufficient to phenocopy the transcriptome rescue in both microglial and endothelial cells after GAS infections, suggesting that IL-17A is a major driver of the CNS pathology after GAS infections. Moreover, IL-17A blockade partially rescues BBB permeability to serum IgG, indicating that it may have therapeutic potential in disorders involving anti-neuronal autoantibodies which have been postulated to underlie the CNS pathology in PANDAS/PANS^15–17^. However, we find no evidence that IL-17A operates directly on ECs since it cannot disrupt tight junctions or promote albumin transcytosis in cultured mBECs. This contrast with a previous study showing that IL-17A-induces TJ dysfunction^80^ in an immortalized EC cell line that differs from those found in the brain^81^. Similar to *RORγt^-/-^* mice, IL-17A mAb-treated wild-type mice dramatically upregulate expression of antigen processing and presentation genes in multiple cell types, suggesting a broad role for IL-17A in suppressing antigen presentation across numerous cell types. To our knowledge, this is the first study to document such a role for IL-17A. It would be interesting to explore whether IL-17A similarly represses antigen presentation in other neurological diseases, and whether this effect depends on interplay with IFNγ signaling. Multiple mAbs against IL-17A (secukinumab and ixekizumab) are already approved by the Food & Drug Administration (FDA) for treatment of psoriasis, psoriatic arthritis and ankylosing spondylitis^82^. Our findings in the mouse disease model suggest that these antibodies can be used to treat PANDAS/PANS children who are refractory to current treatments^8,9^.

We also examined the contributions of GM-CSF, an alternate Th17 effector cytokine, in CNS pathology since it is significantly elevated in the serum of PANDAS/PANS patients. We found that GM-CSF is expressed by CD4 T cells only after multiple GAS infections, which is intriguing since PANDAS is thought to result from repeated GAS exposures. In contrast to IL-17A blockade, genetic ablation of GM-CSF in T cells failed to rescue BBB leakage to serum IgG after GAS infections, although at a molecular level some BBB transcripts (e.g. *Itih5, Itm2a*) were increased and inflammatory genes (e.g. *Lcn2*) were decreased in Csf2^ΔCD4^ mutant ECs after GAS infections. Therefore, GM-CSF does not appear to play a key role for EC function after GAS infections. Similar, the normalization of homeostatic and the reduction in disease-associated microglia transcripts is less significant in Csf2^ΔCD4^ compared to *RORγt* mutant and IL-17A mAb-treated wild-type mice. While Type I IFN response genes were strongly downregulated in Csf2^ΔCD4^ microglia similar to RORγt mutant and IL-17A mAb-treated conditions, the upregulation of some chemokine gene (e.g. *Ccl3* and *Ccl4*) is not rescued in Csf2^ΔCD4^ microglia. Finally, antigen presentation genes were largely rescued in Csf2^ΔCD4^ microglia consistent with published reports that GM-CSF upregulates MHC II and CD74^83^. Therefore, few of the phenotypes rescued in RORγt mutant mice after GAS infections appear attributable to GM-CSF, suggesting that IL-17A may be the predominant driver of the CNS pathology after GAS infections.

Currently it is unclear whether PANDAS/PANS has a neuroinflammatory origin, although higher neuroinflammation has been seen in the basal ganglia of a small number of PANDAS patients^12^ using a PET ligand for TSPO that cannot distinguish between activated microglia and astrocytes^84^. Our mouse model supports microglial, rather than astrocytic reactivity, in response to repeated intranasal GAS exposures. The analysis of serum samples from 23 acute PANDAS/PANS patients and 11 age- and sex-matched healthy controls shows a significant elevation in 12 of the 45 cytokines, chemokines and growth factors. Among the elevated proteins were six that were also highly upregulated by mouse microglia after GAS infections (CCL2, CCL3, CCL4, CCL5, CXCL10 and TNF), further supporting a critical role for microglial activation as a driver of CNS pathology. Some of the elevated cytokines seen in PANDAS/PANS sera can be attributed to GAS infections since patients with invasive GAS diseases have elevated levels of IL- 1β, IL-6, IL-8, IL-10 and IL-18 compared to children with non-invasive GAS infections^85^. Moreover, stimulation of human peripheral blood mononuclear cells (PBMCs) with heat-killed lyophilized GAS for 24 hours evokes production of IL-1β, IL-2, IL-6, IL-8, IFNγ, TNF and GM-CSF^86^. Among serum proteins significantly elevated in PANDAS/PANS, only IL-6 and CXCL10 were shown to be upregulated in acute *S. pyogenes* pharyngitis in adults^87^. In the same study, CCL2, CCL4, CCL5, IL-7 and TNF – all significantly upregulated in PANDAS/PANS cases compared to controls – were unchanged or downregulated in serum following *S. pyogenes* pharyngitis challenge. Overall, these studies suggest some specificity for the cytokine signature seen in PANDAS/PANS children’s sera. Importantly, several cytokines and growth factors including CCL2, CCL5, IL-1RA, PDGFBB and HGF potently upregulated transcytosis in ECs, suggesting that these elevated cytokines in PANDAS/PANS patient sera may induce endothelial barrier breakdown during the acute phase of the disease. These findings support a neuroinflammatory origin of CNS sequelae (SC/PANDAS) and provide some additional mechanisms that can drive the entry of pathological antibodies into the CNS in the acute phase of the disease^4^.

Distinguishing PANDAS/PANS from common childhood neuropsychiatric disorders like Tourette’s syndrome (TS) and obsessive-compulsive disorder (OCD) is a key challenge for clinicians. Although numerous diagnostic approaches ranging from autoantibody detection, to brain imaging, to neuropsychiatric evaluation^7^ have been proposed for PANDAS/PANS, the lack of definitive diagnostic tests is a major obstacle for diagnosis and treatment of patients, as well as the advancement of clinical research. Serum levels of TNF and IL-12 have been temporally correlated with symptom exacerbation in children with tic disorders and OCD^88, 89^. In addition, immune phenotypes linked to TS include elevation of inflammatory cytokines IL-6, IL-12, IL-17A and TNF^88, 90^. Polymorphisms in *IL1RN* – which encodes IL-1RA, one of markers upregulated in PANDAS/PANS serum – have been previously associated with TS in two small association and family-based studies^91, 92^. Further investigation is needed to understand whether the candidate biomarkers identified in the current study can differentiate between PANDAS/PANS, TS or OCD. The identification of reliable biomarkers could resolve some of the controversy around these disorders and, together with advances in diagnosis and treatment protocols, will lessen the disease burden for PANDAS/PANS patients and their families.

## SUPPLEMENTAL INFORMATION

**Document S1**. Figures S1-S6 and Tables S10 and S11

**Table S1**. Sequencing run information, related to **Figures 1** through **6**

**Table S2**. Differentially expressed genes (DEGs) by cell type, with and without FACS enrichment, related to **Figure 1.**

**Table S3**. Genes differentially expressed by CNS endothelial cells and microglia, related to **Figures 2** and 3

**Table S4**. Gene sets used for enrichment analysis, related to **Figures 2** and **3**

**Table S5**. DEGs in perivascular macrophages, related to **Figure 4**

**Table S6**. Cytokine, chemokine and growth factor levels in PANDAS/PANS patient sera and lower detection limit for the assay related to **Table 1**

**Table S7**. DEGs in RORγt^-/-^, IL-17A mAb-treated and Csf2^ΔCD4^ conditions, related to **Figures 5** and **6**

**Table S8**. DEGs in primary microglia after cytokine treatment, related to **Figure S6**

**Table S9**. GO pathway analysis of DEGs in primary microglia after cytokine treatment, related to Figure S6

**Table S10**. Patient demographic data, related to **Table 1**

**Table S11**. *Il17ra* expression in ECs and microglia, related to **Figure S6**

**Document S2**. Scripts for processing of scRNAseq data

**Document S3**. Scripts for processing of MERFISH data

## AUTHOR CONTRIBUTIONS

Conceptualization: CRW, TC, DA; Animal experiments: CRW, DA; RNA sequencing experiments: CRW; MERFISH experiments and analysis: VDL, TEF, CRW, DPS; Bioinformatic analyses: CRW, TEF, VM; Flow cytometry experiments: CRW, LB, SJH; Immunohistochemistry experiments: CRW, LB, SJH, DA; *In situ* hybridization experiments: DA, CRW; Cell culture experiments: CRW, PAF, TC, DA, NA, PA; Patient history & sample collection: SLD, SS, WV; Processing and analysis of patient samples: TC, NA, PA; Statistical analyses: CRW; Resources: DA, BC, DPS, RC, WE; Funding acquisition: DA, TC; Supervision: DA, TC, DPS; Writing: CRW, DA; Revising: CRW, TC, DA.

## ACKNOWLEDGEMENTS

We thank Michael Kissner and Rosemary Gordon-Schneider from the Columbia Stem Cell Initiative Flow Cytometry Core for technical support; Erin Bush and Izabela Krupska from the Single Cell Analysis Core, JP Sulzberger Columbia Genome Center (CUIMC) for library construction and sequencing; S.V. Pollak from the Biomarkers Core Lab, Irving Institute for Clinical and Translational Research (CUIMC) for multiplex cytokine analysis; Drs Ai Yamamoto (CUIMC) and Arnold Han (CUIMC) for equipment use; Dr. Ilir Agalliu (Albert Einstein College of Medicine) for advice on statistical analyses; Dr. Chenghua Gu (Harvard Medical School) and Dr. Wassim Elyaman (CUIMC) for reagents; Dr. Alfred Simkin for the find_nearest_neighbors2 python script; and Drs Jennifer Bain, MD (CUIMC), Wendy Silver, MD (CUIMC), Jay Selman, MD (CUIMC), Rebecca Hommer, MD (NIMH) and Hannah Z. Street (CUIMC) for help with patient recruitment and sample collection.

This work was supported in part by the following funding: NIH/NIMH grant R01MH112849 (DA, TC, CRW); NIH/NIMH grant R56MH109987 (DA, TC); NIH/National Institute of Neurological Disorders and Stroke grant 2T32NS064928 (CRW); International OCD Foundation research grant and Global Lyme Alliance research grant. We are grateful to the following major donors for their generous support of this research: Tom and Patti Walz, Newport Equities LLC, PANDAS Network, Northwest PANDAS/PANS Network, Steve and Wendy Swyter, as well as to many families whose children were affected by this disease for their contributions.

## CONTACT FOR REAGENT AND RESOURCE SHARING

Further information and requests for resources and reagents should be directed to the Lead Contact, Dritan Agalliu.

## DATA AND CODE AVAILABILITY

Raw sequencing data, metadata and count tables for scRNAseq samples have been made available in the Gene Expression Omnibus (GEO). Raw sequencing data and metadata for bulk RNA sequencing of primary microglia are also available in GEO. MERFISH processed data is available in GEO. The MERFISH raw output files cannot be deposited in a public repository due to storage restrictions but can be made available upon request. All GEO accession numbers are listed in the key resources table. The code to reproduce this study’s processed data is available in this paper’s supplementary information.

## EXPERIMENTAL MODEL AND SUBJECT DETAILS

### Mice

Experiments involving mice were approved by Columbia University’s Institutional Animal Care and Use Committees. Mice were bred in the CUIMC vivarium, under 12-hour light/12-hour dark, pathogen-free conditions. Female mice were used for all experiments, except the time course analysis of Th17 cell subtypes by flow cytometry (**Figures 6A-B; S6B-C**) and *Csf2* recombination confirmation flow cytometry (**Figure S6D**), which used even numbers of males and females. The *RORγt^eGFP^* mice^19, 46^ (B6.129P2(Cg)-Rorctm2Litt/J, strain 007572) were obtained from the Jackson Laboratory. The *Csf2^fl/fl^* mouse strain^54^ was provided by Bogoljub Ciric (Thomas Jefferson University, Philadelphia, PA). *CD4-CreER^T2^* transgenic mice (B6(129X1)-Tg(Cd4- cre/ERT2)11Gnri/J, strain 022356)^53^ were obtained from the Jackson Laboratory and crossed to Csf2^fl/fl^ mice for two generations. CD4-CreER^T2+/-^ Csf2^fl/fl^ males were mated to Csf2^fl/fl^ females to generate CD4-CreER^T2+/-^ Csf2^fl/fl^ experimental mice and Csf2^fl/fl^ littermate controls. P16 pups were intraperitoneally injected daily with 100 µg of (Z)-4-Hydroxytamoxifen (Millipore Sigma, H7904), dissolved in 50 µL of corn oil (Millipore Sigma, C8267) for 5 days (P16-P20). *TMEM119- tdTomato*^41^ and *CX3CR1-GFP*^40^ reporter mouse lines were generously provided by Wassim Elyaman (CUIMC).

### Human serum studies

The experiments with human sera were approved by Columbia University’s Institutional Review Board (IRB-AA99). The NIMH sera used in this study were analyzed in a previous publication^93^, and were obtained from the NIMH. Informed consent was obtained from all subjects.

## METHOD DETAILS

### GAS intranasal infections

Mice received weekly intranasal inoculations with either a suspension of *Streptococcus pyogenes* [Group A *Streptococcus* (GAS)], or phosphate-buffered saline (PBS) control, starting at P21-P28. We used a recombinant GAS strain expressing a 2W epitope-tagged M protein as described^18–20^. GAS was streaked out on new blood agar plates each week. Culture media consisted of an autoclaved solution of 3% Todd-Hewitt Broth (Bacto, 90003-430) and 2% Neopeptone (Bacto, 90000-268). Several GAS colonies were used to inoculate 10 mL of culture medium and incubated overnight at 37°C in 5% CO_2_. The following day, the culture was diluted to an OD_600_ of 0.2 and grown to OD_600_ 0.6, centrifuged and washed in 1 mL PBS (without Ca^2+^ and Mg^2+^) and resuspended in 110 µL of PBS. The GAS suspension was kept briefly on ice prior to intranasal infections. All intranasal infections were performed in an ABSL2 vivarium facility. Mice were immobilized with light anesthesia and a P20 pipette was used to drip GAS suspension into nostrils. To reduce lethality due to sepsis, a smaller GAS dose was used during the first two weeks (the first GAS inoculation is 8 x 10^7^ CFU per nostril, the second is 12 x 10^7^ CFU per nostril, and the third, fourth and fifth are 2 x 10^8^ CFU per nostril). During the first two weeks of GAS infections, mice were provided with nutritional supplements (ClearH_2_O, 72-27-5022).

### Neutralizing antibody treatment

Starting 24 hours prior to the first GAS infection, mice were injected intraperitoneally twice weekly with 500 µg of either InVivoMAb anti-mouse IL-17A monoclonal antibody, clone 17F3 (Bio X Cell catalog, BE0173), or mouse IgG1 isotype control monoclonal antibody, clone MOPC-21 (Bio X Cell, catalog BE0083), in 100 µL of dilution buffer (Bio X Cell catalog IP0070 and IP0065, respectively).

### Single-cell RNA sequencing

Mice were anesthetized with isoflurane and perfused intracardially with PBS for 3 minutes. Nasal associated lymphoid tissue (NALT), olfactory epithelium (OE), or olfactory bulb (OB) were dissected and placed in Hanks’ Balanced Salt Solution (HBSS) without Ca^2+^ and Mg^2+^ and cut up with a sterile scalpel blade. Two or three animals were pooled per sample. Tissue was then placed in C Tubes (Miltenyi Biotec, 130-093-237), along with dissociation reagents from the MACS Neural Tissue Dissociation Kit (P) (Miltenyi Biotec, 130-092-628). Samples were loaded onto a gentleMACS Octo Dissociator with Heaters (Miltenyi Biotec, 130-096-427) and the 37C_NTDK_1 program was run. Following dissociation, samples were filtered through a 70 µm cell strainer, washed in HBSS and resuspended in MACS buffer with myelin removal beads (Miltenyi Biotec, 130-096-733), then purified with an LS column (Miltenyi Biotec, 130-042-401), according to manufacturer instructions. Eluent was washed twice, incubated with DR (Bio-Legend, 424101, 1:1000) and CD16/CD32 Fc block (BD Biosciences, 553141, 1:200) at room temperature for 15 minutes. Cells were washed and incubated with antibodies against CD31 (FITC, BD Biosciences, 561813, 1:200) and CD11b (BV421, BioLegend, 101235, 1:100) for 30 minutes on ice. Cells were washed and resuspended in FACS buffer with propidium iodide (1:10,000). Live, nucleated cells (DR^+^ PI^lo^) were sorted on a FACSAria II (BD), equipped with 355 nm, 405 nm, 488 nm, 561 nm and 640 nm lasers and a 130 µm nozzle. In a subset of experiments, CD31^+^ and CD11b^+^ populations were collected to enrich for cell types of interest. Sequencing was performed by the Columbia Single Cell Core using 10X Genomics Chromium Single Cell 3’ technology, with reads aligned to the mm10-2020-A transcriptome.

### Immunofluorescence

Mice were anesthetized with isoflurane and perfused intracardially with PBS for 4 minutes, followed by 4% paraformaldehyde (PFA) for 6 minutes. Brains were extracted and post-fixed in 4% PFA for 4-6 hours, then washed three times in PBS, incubated overnight in 30% sucrose, embedded in Tissue-Plus O.C.T. compound (Fisher, 4585) and stored at -80°C. Coronal sections (12 µm) were cut on a Leica CM3050 S Cryostat and stored at -80°C. For immunofluorescence staining, slides were washed in PBS for 10 minutes, incubated for 1 hour at room temperature in blocking buffer (10% BSA in 1X PBS with 0.1% Triton-X-100), and with primary antibodies diluted in PBST (0.1% Triton-X-100 in 1X PBS) with 1% BSA overnight at 4°C. The table below provides a complete list of immunofluorescence antibodies used for this study.

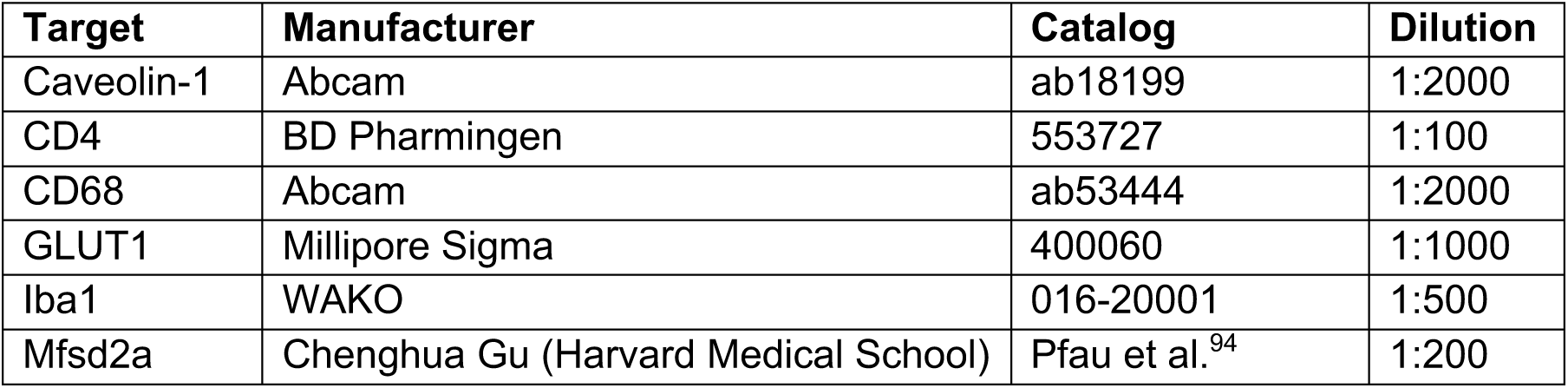

Slides were incubated in primary antibodies at 4°C in a humidified chamber overnight. After three 10-minute washes with PBST, slides were incubated for 2 hours at room temperature in secondary antibodies diluted in PBST with 1% BSA. These were conjugated to AlexaFluor 488 (1:1000), AlexaFluor 594 (1:1000), or AlexaFluor 647 (1:500). Following three washes with PBST and two with PBS, slides were cover slipped with Vectashield (Vector Labs, Burlingame, CA) containing the nuclear stain DAPI, then sealed with clear nail polish and stored at -20°C.

### *In situ* hybridization

Plasmids were obtained from Transomic Technologies and the antisense mRNAs were synthesized using the Digoxigenin RNA Labeling Kit (SP6/T7; Roche, 11175025910). DIG RNA *in situ* hybridization (ISH) and fluorescent *in situ* hybridization (FISH) experiments were performed as previously described^95, 96^. Mice used for ISH or FISH were perfused for 4 minutes with PBS and brains were dissected out and embedded in O.C.T.

### Multiplexed error-robust *in situ* hybridization (MERFISH)

Mice were anesthetized with isoflurane and intracardially perfused with cold RNAse-free PBS for 4 minutes. Brains were dissected out and immediately embedded in Tissue-Plus O.C.T compound and stored at -80°C until samples could be shipped overnight on dry ice to UMass Chan Medical School. Samples were prepared according to the Vizgen Fresh Frozen Tissue Sample Preparation protocol. Tissue was sectioned in 10 µm slices onto a functionalized coverslip covered with fluorescent beads. Each coverslip contained a section from one PBS sample and a section from one GAS sample. Tissue on coverslips was fixed for 15 minutes at room temperature in 4% paraformaldehyde in PBS, followed by three washes with PBS. Tissue was then permeabilized in 70% ethanol for 24 hours, washed with PBS and incubated with blocking solution for 1 hour, followed by 1 hour of incubation with the primary antibody against vessel marker CD31 (BioLegend, 102502), diluted 1:20 in blocking solution (Vizgen). The tissue was then washed three times with PBS and incubated for 1 hour with an oligo-conjugated secondary antibody diluted 1:1000 in blocking solution. The sample was washed three times with PBS and fixed for 15 minutes at room temperature in 4% paraformaldehyde in PBS, followed by three washes with PBS. After a 30-minute wash with Formamide Wash Buffer (30% formamide in 2X saline sodium citrate, or SSC) at 37°C, the MERFISH library mix was added and allowed to hybridize for 48 hours. Sample was then washed and incubated at 47°C with Formamide Wash Buffer twice for 30 minutes each and then the tissue was embedded in a polyacrylamide gel followed by incubation with tissue clearing solution (2X SSC, 2% SDS, 0.5% v/v Triton X-100, and proteinase K 1:100) overnight at 37°C. Then, tissue was washed and hybridized for 15 minutes with the first hybridization buffer containing DAPI, polyT and the readout probes associated with the first round of imaging. After washing, the coverslip was assembled into the imaging chamber and placed into the microscope for imaging. MERFISH imaging was performed as previously described^97^ with parameter files provided by Vizgen. Briefly, the sample was loaded into a flow chamber connected to the Vizgen Alpha Instrument. First, a low-resolution mosaic image was acquired (405 nm channel) with a low magnification objective (10x). Then the objective was switched to a high magnification objective (60x) and seven 1.5-μm z-stack images of each field of view position were generated in 749 nm, 638 nm and 546 nm channels. A single image of the fiducial beads on the surface of the coverslip was acquired and used as a spatial reference (477 nm channel). After each round of imaging, the readout probes were extinguished, and the sample was hybridized with the next set of readout probes. This process was repeated until combinatorial FISH was completed.

Raw data were decoded using the MERlin pipeline (Vizgen, v0.1.12) using the relevant library codebook. Cell boundaries were segmented in each FOV using a seeded watershed algorithm with DAPI signal as the seed and poly-T signal as the watershed channel. The volume, X position, and Y position of these cell boundaries, as well as the probe counts within each cell boundary, were output for further analysis.

Probes for the following transcripts were used: Abca7, Abcc3, Abcg2, Ablim1, Actb, Acvrl1, Adam10, Adam17, Adgrf5, Adgrl4, Adora1, Aff3, Ago4, Agt, Ahr, Akap12, Aldoc, Anxa1, Ap2m1, Aqp4, Arc, Arg1, Arhgap29, Arl15, Arpc2, Atmin, Atp10a, Axl, Baiap2l1, Bard1, Bin1, Birc5, Bmp6, Brca1, Btk, C1qa, C1qb, C1qbp, C1qc, C3, C3ar1, C4a, C5ar1, Cald1, Casp7, Casp8, Cass4, Ccl2, Ccl22, Ccl3, Ccr2, Cd14, Cd163, Cd27, Cd33, Cd3e, Cd4, Cd47, Cd68, Cd72, Cd74, Cd79a, Cd84, Cd86, Cd8a, Cdh5, Cdh9, Cemip, Cenpa, Cgnl1, Chek2, Chit1, Cldn5, Clec7a, Clic4, Clu, Cmtm8, Cobll1, Col6a3, Cotl1, Crim1, Csad, Csf1r, Csf2, Csf2ra, Csf2rb, Cspg4, Cstb, Ctgf, Ctsb, Cx3cr1, Cxcl10, Cyr61, Dach1, Dapk1, Dclre1a, Ddx58, Des, Dlc1, Dna2, Dock2, Dock9, Dusp1, E2f1, Ebf1, Ebi3, Ece1, Edn1, Edn3, Efnb2, Egfl7, Egfr, Egr1, Elovl7, Emcn, Emp1, Eng, Enpp6, Entpd1, Epas1, Epb41l4a, Epha1, Erg, Esam, Esyt2, Fancd2, Fbrs, Fbxw17, Fcer1g, Fcgr1, Fcgrt, Fcrls, Fgd2, Flcn, Flnb, Flt1, Flt3, Flt4, Fn1, Folr2, Fos, Foxj1, Foxp1, Ftsj3, Gad1, Gad2, Galnt18, Gbp2, Gfer, Gna13, Gna15, Gpi1, Gpr183, Gpr34, Gpr84, Grb10, Grn, Gusb, H2-D1, Hcar2, Heg1, Hells, Hexb, Hmgb2, Hmox1, Icam1, Ifi30, Ifih1, Igsf6, Il1a, Il1b, Il1rl2, Il1rn, Il21r, Il2rg, Il3ra, Impact, Inpp5d, Irak4, Irf7, Irf8, Itga1, Itga2, Itga6, Itgae, Itgal, Itgam, Itgax, Itgb1, Itgb3, Itgb5, Itm2a, Jcad, Jun, Kif26a, Kit, Lacc1, Lair1, Lama2, Lcn2, Ldb2, Ldlrad3, Lef1, Lfng, Lgals3, Lgmn, Lig1, Liph, Lmnb1, Lrch3, Lrp1, Lrrc3, Ly6g, Ly9, Lyn, Lyve1, Lyz2, Mb21d1, Mcm2, Mcm5, Mcm6, Mctp1, Mecom, Mef2c, Meg3, Mertk, Mfsd6l, Mgat1, Mki67, Mkl2, Mmp2, Mmp9, Mpnd, Mrc1, Ms4a1, Ms4a6d, Ms4a7, Msn, Mvp, Mybl2, Myrip, Nampt, Napsa, Ncaph, Ncf1, Nckap1l, Nebl, Nfib, Nlrp3, Nos3, Nostrin, Nrm, Ocln, Olfm2, Olfml3, Osm, Osmr, P2ry12, Palmd, Pam, Pard3, Pcna, Pde10a, Pde4d, Pdgfra, Pdgfrb, Pdlim5, Pdpn, Pdzrn3, Pecam1, Picalm, Pik3cg, Pla2g4a, Plac8, Plcb4, Plcg2, Plekha6, Plekhg1, Plp1, Plpp1, Pltp, Plxdc2, Plxna2, Podxl, Pole, Ppfibp1, Prdx5, Prickle2, Prkg1, Pros1, Psmb8, Ptk2b, Ptpn6, Ptprc, Ptprg, Ptprj, Ptprm, Pvalb, Qars, Rad23b, Rad51, Rae1, Rapgef4, Rbms3, Rngtt, Rora, Rorb, Rrm2, Rsad2, Rundc3b, Sall1, Sardh, Sash1, Sdf4, Sell, Serpina3n, Serpine1, Serpinf1, Siglecf, Slamf8, Slamf9, Slc17a6, Slc1a1, Slc39a10, Slc40a1, Slc4a4, Slc7a1, Slco2b1, Slfn8, Smc3, Snx2, Sorbs1, Sorbs2, Sox10, Sox2, Sox9, Spi1, Spp1, Sptbn1, Srgn, Sst, St8sia6, Stat1, Syk, Syne1, Syne2, Tacc1, Tead1, Tek, Tgfa, Tgfbi, Tgfbr1, Tgfbr2, Tgm2, Thsd4, Timeless, Timp2, Timp3, Tlr2, Tlr4, Tmem119, Tmem173, Tmtc1, Tnfrsf11a, Tnfrsf1a, Tnfsf13b, Top2a, Trem1, Trem2, Trim47, Tshz2, Tspan33, Ttll12, Ttr, Txnrd1, Tyrobp, Unc13b, Ung, Utrn, Vac14, Vav1, Vcl, Vegfa, Vim, Vip, Vtn, Vwf, Was, Wwtr1, Xpo1, Zbp1, Zbtb46.

### Flow cytometry

Mice were anesthetized with isoflurane and intracardially perfused with PBS for 4 minutes. Brain, as well as combined nasal associated lymphoid tissue/olfactory epithelium (NALT/OE), were dissected out, placed in cold Dulbecco’s Modified Eagle’s Medium (DMEM) (Genesee, 25-500), and pressed through a cell strainer with the end of a sterile syringe. Samples were collected in 10 mL of a 30% Percoll (Cytiva, GE17-0891-01) suspension in DMEM, and underlaid with 1 mL of 70% Percoll. Spleen samples were suspended in 3 mL of Red Blood Cell Lysis Buffer (155 mM NH_4_Cl, 10 mM KHCO_3_, 0.1 mM EDTA) at room temperature for 10 minutes. Samples were centrifuged at 4°C at 1300 rcf for 30 minutes, then immune cells were collected at the interface. All samples were then filtered through a cell strainer, washed with 2 mL DMEM, and centrifuged for 10 minutes at 800 rcf, and resuspended in T cell media (RPMI with fetal bovine serum (FBS) 1:10, penicillin/streptomycin 1:100, MEMNEAA 1:100, glutamine 1:100, 55 µM β-mercaptoethanol 1:1000) with 1X Cell Stimulation Cocktail (plus protein transport inhibitors) (eBioscience, 00-4975- 93). Samples were incubated at 37°C for 4 hours, washed with FACS buffer, and incubated in anti-CD16/CD32 Fc block (BD Biosciences, 553141, 1:200) for 15 minutes on ice. All subsequent steps were performed at 4°C. After washing with FACS buffer, samples were incubated in cell surface stains (a complete list of antibodies is included in table below) with either Fixable Viability Dye 780 (Invitrogen, 65086518, 1:4000) or Live/Dead Aqua (ThermoFisher, L34965, 1:1000) in the dark for 1 hour, fixed for 30 minutes using the Intracellular Fixation & Permeabilization kit (eBioscience, 88-8824-00) according to manufacturer instructions. Intracellular stains were diluted in 1X permeabilization buffer and incubated for 1 hour, then washed in permeabilization buffer and resuspended in FACS buffer.

Samples were analyzed in the Columbia Stem Cell Initiative Flow Cytometry Core. Time course experiments were analyzed using a ZE-5 analyzer (Bio-Rad, Hercules, CA) equipped with 355 nm, 405 nm, 488 nm, 561 nm, and 640 nm lasers. Compensation controls used splenocytes for most cell surface markers and compensation beads (BD Biosciences, 552844 for rat hosts; ThermoFisher, 01-3333-41 for mouse or Armenian hamster hosts) for cytokines. All other flow cytometry experiments were analyzed using a NovoCyte Penteon (Agilent, Santa Clara, CA) equipped with 349 nm, 405 nm, 488 nm, 561 nm and 637 nm lasers. Compensation controls used splenocytes for live/dead control and compensation beads for all other markers.

All flow cytometry data analysis was performed using FlowJo 10.5 (FlowJo, LLC). Gates for forward and side scatter, singlets, live cells and CD4 T cells were set by eye, while all other gates were set using fluorescence minus one (FMO) controls, with a typical cutoff of <1% of the population.

| **Target and fluorophore** | **Manufacturer** | **Catalog** | **Dilution** |
| --- | --- | --- | --- |
| CCL5 PE-Cy7 | BioLegend | 149105 | 1:200 |
| CD31 FITC | BD Biosciences | 553372 | 1:100 |
| CD4 BV605 | BD Biosciences | 563151 | 1:100 |
| CD11b BV421 | BioLegend | 101235 | 1:100 |
| CD11b PerCP-Cy5.5 | BioLegend | 101227 | 1:100 |
| CD44 BV421 | BioLegend | 103040 | 1:200 |
| CD45 AF700 | BioLegend | 103127 | 1:100 |
| CD45 BUV395 | BD Biosciences | 564279 | 1:100 |
| CD45 BV421 | BD Biosciences | 563890 | 1:100 |
| CD62L PE-Cy7 | BioLegend | 104417 | 1:100 |
| CD74 BV711 | BD Biosciences | 740748 | 1:400 |
| CX3CR1 BV786 | BioLegend | 149029 | 1:400 |
| GM-CSF FITC | BioLegend | 505403 | 1:200 |
| IFNγ APC | BD Biosciences | 554413 | 1:100 |
| IL-17A PE | BD Biosciences | 559502 | 1:100 |
| MHC II I-A/I-E APC-Cy7 | BioLegend | 107627 | 1:400 |
| P2RY12 PE | BioLegend | 848003 | 1:100 |

### Cell culture

Primary mouse brain endothelial cells (mBECs; Cell Biologics, C57-6023) were cultured as monolayers at 37°C with 5% CO_2_, and used to evaluate the effect of cytokines *in vitro*. Cells were grown to confluence on collagen IV-coated (Corning, CB-40233) dishes in endothelial cell media (Cell Biologics, M1168) supplemented with 10% FBS (Cytiva, SH30071.03) and supplements recommended by the supplier. One day prior to cytokine treatment, cells were switched to 2% FBS without growth factor supplements. Media containing either mouse IFNγ (R&D Systems, 485-MI-100), mouse IL-17A (R&D Systems, 7956-ML-025), mouse GM-CSF (R&D Systems, 415- ML-010) or vehicle (PBS with Ca^2+^ and Mg^2+^) was applied to corresponding wells.

Trans-endothelial electrical resistance (TEER) was measured in real time using an electric cell-substrate impedance sensing (ECIS) instrument (Applied BioPhysics, ZTheta 96 Well Array Station) as previously described^96^. mBECs were plated on 96-well plates containing electrode arrays (Applied BioPhysics, 96W20idf). Cytokines or vehicle were added at a concentration of 50 ng/mL, except IL-1β (R&D Systems, 401-ML) and TNF (R&D Systems, 410-MT), which were added at 10ng/mL. Resistance was monitored over 24 hours from the start of cytokine treatment. The area under the curve (AUC) was calculated for each condition and normalized to AUC of vehicle-treated cells, with values for each independent experiment compared by one-way ANOVA.

Albumin transcytosis was measured *in vitro* by culturing mBECs on collagen IV-coated 3.0-µm PET membrane inserts for 24-well plates (Corning, 353096). Cytokines or vehicle were added at a concentration of 100 ng/mL, with 500 ng/mL lipopolysaccharide (LPS) used as a positive control. Cells were switched to endothelial cell media with 2% FBS and without phenol. Media added to wells contained bovine serum albumin at 400 µg/mL, while media added to inside of transwell inserts contained albumin conjugated to AlexaFluor 647 (ThermoFisher, A34785) at 400 µg/mL. Cells were incubated at 37°C with flow-through samples collected from bottom wells at 30 minutes, 1, 2, 4 and 6 hours). Absorbance of flow-through was quantified using the accuSkan FC plate reader (Fisher Scientific, 14-377-576). Background fluorescence was subtracted and AUC normalized to untreated for each experiment.

Mixed glial cultures were generated from forebrains of P0-P2 C57BL/6J pups. After 7-14 days, microglia were obtained using a shake off protocol and cultured in serum-free media containing neurobasal solution (ThermoFisher, 211039049), insulin (Sigma, I6634, 1:100), sodium pyruvate (Invitrogen, 11360-070, 1:100), pen/strep (Life Technologies, 15140-122, 1:100), SATO (containing transferrin, BSA, putrescine, progesterone, and sodium selenite; 1:100), thyroxine, GlutaMAX (Life Technologies, 35050-061, 1:100), B27 (ThermoFisher, 17504-044, 1:50), N-acetyl cysteine (1:1000), and mouse M-CSF (Shenandoah Biotechnology, 200-08- 10, 1:1000). Cultures were treated with 100 ng/mL of either IFNγ, IL-17A or GM-CSF for 24 hours, then washed and collected into TRIzol for RNA extraction and sequencing.

### Multiplex immunoassays

#### Mouse olfactory bulb multiplex immunoassay

Twenty-four hours after the final GAS inoculation, pairs of OBs were dissected and flash frozen in liquid nitrogen, then stored at -80°C. Samples were pulverized on ice in cell lysis buffer (Abcam, ab152163) with protease inhibitor cocktail (ThermoFisher, 78440) and EDTA, using an electric pestle. To remove detergents that may interfere with downstream analysis, the resulting supernatant was dialyzed overnight against PBS with a 2 kDa cassette (ThermoFisher, 66205). After normalization to total protein concentration by Pierce bicinchoninic acid assay (ThermoFisher, 23225), analytes were measured by the Irving Institute for Clinical and Translational Research Biomarkers Core Laboratory using a custom mouse Luminex panel (ThermoFisher, PPX-12-MXEPUF3). Samples were run in duplicate, and standard curves were generated for each analyte. Undetectable values were replaced with half of the lower detection limit for purposes of statistical comparison.

#### Patient serum multiplex immunoassay

Serum protein concentrations were measured by the Irving Institute for Clinical and Translational Research Biomarkers Core Laboratory using a 45- Plex Luminex assay (Invitrogen, EPX260-26088-901). Samples were run in duplicate, and standard curves were run for each analyte. Undetectable values were replaced with half of the lower detection limit (**Table S6**) for statistical comparison.

## QUANTIFICATION AND STATISTICAL ANALYSIS

### Analysis of scRNAseq data

Single-cell RNA sequencing data (**Table S1**) was analyzed using Seurat package v4.0.2^98^ in RStudio. Upon data import, genes detected in fewer than three cells, and cells with fewer than 200 genes were excluded. Cells were removed from the merged data set if they had fewer than 1,000 or more than 50,000 molecules detected, or greater than 20% mitochondrial reads. Data was normalized and highly variable features identified using default parameters, then scaled, followed by linear dimensional reduction using PCA. Dimensionality of the data was selected using the Elbow plot method with 50 dimensions, and cells were clustered with a resolution of 1 for OB, 0.4 for endothelial cells and microglia, and 2 for NALT/OE. Dimensionality reduction for visualization was performed with *t*-distributed stochastic neighbor embedding (*t*-SNE). The Harmony package v1.0^99^ was used for batch correction.

Cluster identity was assigned using the following cell type markers: neurons (*Map2*, *Snap25*), astrocytes (*Gfap*, *Aqp4*), olfactory ensheathing cells (*Frzb*), oligodendrocytes/oligodendrocyte precursor cells (*Pdgfra*), endothelial cells (*Cldn5*, *Pecam1*), pericytes (*Pdgfrb*, *Atp13a5*), fibroblasts (*Col1a1*, *Fbln1*), microglia (*Tmem119*, *P2ry12*), macrophages (*Aif1*, *Plac8*), neutrophils (*Ly6g*, *Camp*), dendritic cells (*Xcr1*, *Ccr9*, *Cd209a*), B cells (*Cd19*), CD4 T cells (*Cd4*), CD8 T cells (*Cd8a*), NK cells (*Klrb1c*), and γδ T cells (*Cd163l1*). To determine the number of differentially expressed genes for each cell type, FindMarkers was used to compare GAS to PBS conditions with minimum percent of 0.2, fold-change threshold of 0.5 and p-value cutoff of less than 0.05. Expression of *ex vivo* activation genes (*Dusp1*, *Fos*, *Hist1h1d*, *Hist1h2ac*, *Jun*, *Nfkbid*, *Nfkbiz*) were added using AddModuleScore and plotted against *Ccl3* and *Ccl4* using FeatureScatter.

Additional analysis was performed with BB Browser 3 (BioTuring) software. Signature scores were plotted in BBrowser3 using the following pathway markers: Antigen presentation (*B2m*, *Cd74*, *H2-Aa*, *H2-Ab1*, *H2-D1*, *H2-Eb1*, *H2-K1*, *H2-Q4*, *H2-Q6*, *H2-Q7*, *Tap1*, *Tap2*), disease-associated microglia (*Apoe*, *Axl*, *Cd9*, *Csf1*, *Cst7*, *Itgax*, *Lpl*, *Spp1*, *Tyrobp*), homeostatic microglia (*Cd33*, *Cst3*, *Cx3cr1*, *Fcrls*, *Gpr34*, *Olfml3*, *P2ry12*, *P2ry13*, *Sall1*, *Tmem119*), and interferon signaling (*Ifi30*, *Ifi204*, *Ifi211*, *Ifit1*, *Ifitm3*, *Irf1*, *Irf7*, *Isg15*, *Oas1a*, *Stat1*, *Stat2*).

Gene set enrichment analysis (GSEA)^31^ was performed using curated and database-derived gene lists for blood-brain barrier, response to LPS, inflammation, extracellular matrix, interferon response, antigen presentation, chemokine and cytokine signaling, endothelial cell proliferation, endothelial cell migration, disease-associated microglia, apoptosis, leukocyte chemotaxis and phagocytosis (**Table S4**). Analysis was run using the GSEA desktop tool (Broad Institute, v4.1.0), using pre-ranked weighted settings.

Stacked violin plots (**Figures 5**, **6 and S5**) were generated using scripts by Dr. Ming Tang (divingintogeneticsandgenomics.rbind.io).

### Analysis of MERFISH data

MERFISH data was analyzed in RStudio using Seurat 4.1.0.9005, R 4.0.0 and custom-made scripts as previously described^100^. Cell segmentations with volume < 50µm^3^ or < 10 unique transcripts were first excluded. Cell gene expression data of each cell was then normalized to that cell’s volume and the total transcript count of that cell, then scaled. To correct for global differences in total transcript counts between coverslips (each containing one GAS sample and one PBS sample), we performed ComBat^101^ batch correction (sva 3.38.0).

To identify individual cell types, we performed principle component analysis was performed using the entire probe library (391 transcripts) as the variable features, followed by linear dimensional reduction. Dimensionality of the data was selected using the jackstraw method with 28 dimensions, and cells were clustered with a resolution of 2.4. Dimensionality reduction was performed with *t*-distributed stochastic neighbor embedding (*t*-SNE). Clusters were manually annotated based on the spatial distribution of the cells in the tissue and the expression cell type-specific marker genes: neurons (*Meg3, Gad1*), astrocytes (*Aqp4, Sox9*), olfactory ensheathing cells (*Plp1, Cldn5*), oligodendrocytes/oligodendrocyte precursor cells (*Pdgfra, Sox10*), endothelial cells (*Cldn5*, *Itm2a*), pericytes (*Pdgfrb*), fibroblasts (*Cemip*), microglia (*Tmem119*, *P2ry12*), macrophages (*Mrc1*), neutrophils (*Itgal*, *Mmp9*), and T cells (*Cd3e*, *Cd4, Cd8a*).

Because of imperfections in cell boundary segmentation, a small fraction of cells expressed cell type markers for multiple cell types. Raw images of a subset of these cells were visually inspected using MERSCOPE Visualizer software (Vizgen, 2.1.2589.1) to confirm that these clusters were due to cell segmentation errors (typified by two distinct clusters of cell-type specific transcripts within the same cell boundary). Clusters composed of these “hybrid” cells were removed from the analysis, and embedding and clustering analysis were iteratively repeated until all “hybrid” clusters were removed.

The glomerular, external plexiform, and granular layers of the OB for each sample were outlined using MobileFish and coordinates recorded for point-in-polygon analysis and regional assignment of microglia. Raw counts were normalized to the PBS condition for each batch and used for gene expression analysis. Endothelial cell gene expression was compared on log_2_ fold change values using a one-sample t test. Microglia gene expression between the glomerular and granular layers used a ratio t test. Nearest neighbor analysis of microglial distance to T cells was calculated based the x,y coordinates of the centers of the cell segmentations using a custom python script.

### Analysis of bulk RNAseq data

All sequencing and analysis of microglia cultures were performed by the Single Cell Analysis Core at the JP Sulzberger Columbia Genome Center. Following TRIzol RNA extraction, polyA libraries were prepared with the TruSeq Stranded mRNA kit (Illumina, San Diego, CA). Data processing used RTA (Illumina) for base calling, bcl2fastq2 (v2.19) for converting FASTQ and adaptor trimming, pseudoalignment from mouse transcriptomes GRCm38, kallisto (v0.44.0) for abundance quantification, and DESeq2 for differential expression analysis.

### Statistical analysis

Most statistical analyses were performed by GraphPad Prism 9. All tests were two-sided using a significance level α = 0.05. Outliers were identified and excluded using the ROUT method (Q = 1%). Significance was notated as ns, p > 0.05; *, p < 0.05; **, p < 0.01; ***, p < 0.001. Error bars represent mean with SEM throughout. Differential expression analysis of scRNAseq data was performed using the FindMarkers function in Seurat. Comparisons of more than two conditions underwent Bonferroni correction for multiple comparisons.

### Immunohistochemistry quantification

#### Quantification of microglial number

Three OB sections, corresponding to bregmas 4.5, 4.28 and 3.92, were imaged using a Zeiss AxioImager microscope at magnification 20x for each animal. The number of Iba1^+^CD68^+^ cells (corresponding to activated microglia) in the glomerular layer was manually counted, and averaged across the three sections. Iba1^+^ cells (microglia) were considered CD68^+^ if the CD68 fluorescence occupied more than 50% of the cell surface area, as previously described^18, 19^.

#### Quantification of BBB leakage

Bregma 4.28 sections were imaged using a Zeiss AxioImager microscope at magnification 10x. Using ImageJ^102^, 20 small, rectangular regions of interest (ROIs) were placed around the glomerular layer of the OB, avoiding the vasculature, and average fluorescence quantified for each animal. The same process was repeated for the granular layer.

#### Quantification of EC marker expression

Tiled images of bregma 4.28 sections were taken at 40x using a Zeiss LSM700 confocal microscope, and maximum intensity projections were created. Using FIJI, ROIs across the whole OB were selected using Otsu thresholding on a vessel marker. Then average fluorescence intensity was measured within the ROIs. Due to neuronal expression of *Itih5* in the glomerular layer, ROIs were restricted to the granular layer of the OB for quantification.

