## Supplementary material for "Distinct Th17 effector cytokines differentially promote microglial and blood-brain barrier inflammatory responses during post-infectious encephalitis": Figure S

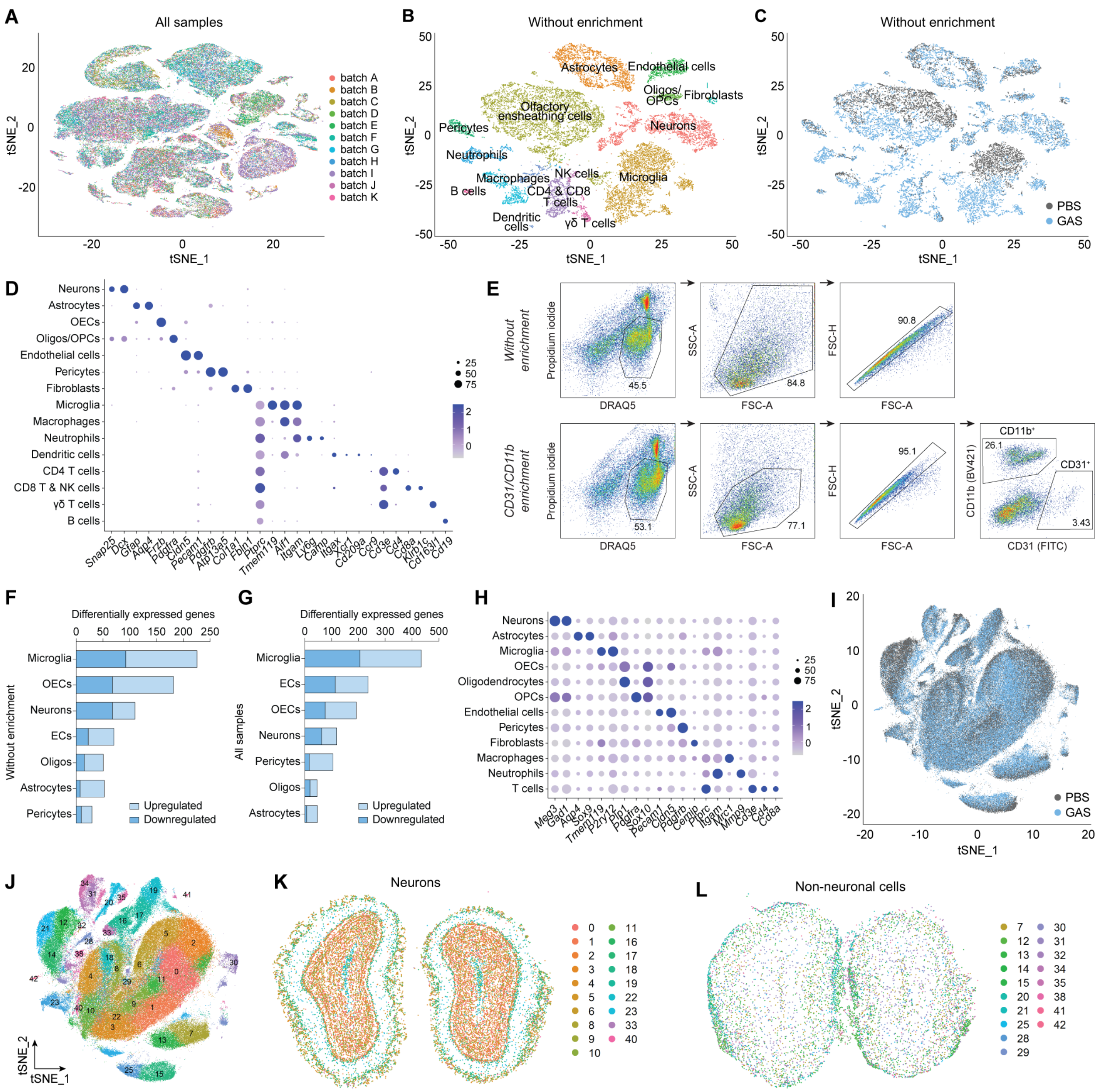


**Figure S1: Identification of olfactory bulb cell types isolated from PBS and GAS-infected mice for scRNAseq and MERFISH analysis, related to Figure 1**

(**A-C**) *t*-SNE plots depicting clusters for all cell types isolated from all samples labeled by sequencing batch (**A**), clusters without enrichment labeled by either cell type (**B**), or sample origin (PBS in gray and GAS in blue; **C**). (**D**) Dot plot of the molecular markers used to assign cell identity in the olfactory bulb (OB) from PBS and Group A *Streptococcus*- (GAS) infected mice for single-cell RNA sequencing (scRNAseq) experiments. The color scale indicates the average level of expression; the dot size indicates the percentage of cell type expressing the gene mRNA. (**E**) Gating strategy for fluorescence-activated cell sorting (FACS) of the myeloid cells and endothelial cells (ECs) isolated from the OB. The top row shows gating of samples without cell type-specific enrichment; the bottom row shows gating of samples enriched for $CD31^+$ (ECs) and $CD11b^+$ (myeloid) cells. (**F-G**) Plots of the numbers of differentially expressed genes (DEGs) by cluster, displaying upregulated (light blue) and downregulated (dark blue) DEGs. (**H**) Dot plot of the molecular markers used to assign OB cell identity in the MERFISH experiments. (**I, J**) *t*-SNE plot illustrating sample clustering from MERFISH experiments labeled by sample origin (PBS in gray and GAS in blue**;** **I**), and cluster number (**J**). (**K, L**) Representative coordinate plots showing labeled clusters of neuronal (**K**), and non-neuronal (**L**), populations in the MERFISH experiment.

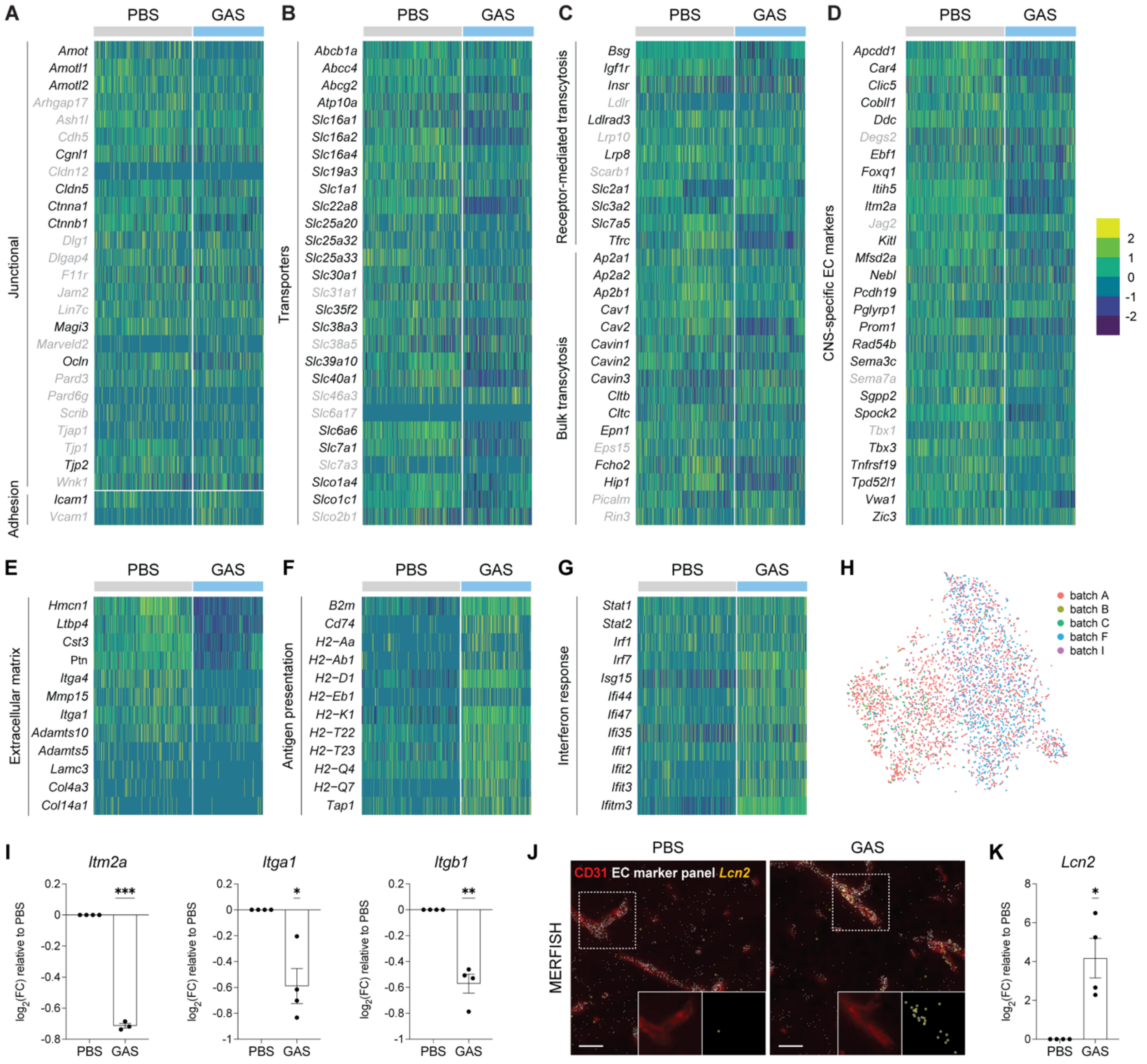


**Figure S2: CNS endothelial cells undergo major transcriptional shifts after multiple intranasal GAS infections, related to Figure 2**

(**A-G**) Heat maps of CNS endothelial cell (EC) gene expression related to blood-brain barrier (BBB) function (**A-D**), extracellular matrix (ECM, **E**), antigen presentation (**F**) and the interferon response (**G**). Genes with black labels are significant (p-adj < 0.05), and those with gray labels are non-significant (p-adj > 0.05). (**H**) *t*-SNE plot of all ECs isolated from the OBs labeled by sequencing batch. (**I**) Downregulation of BBB-related genes *Itm2a*, *Itga1* and *Itgb1* by MERFISH (* $p < 0.05$; ** $p < 0.01$; *** $p < 0.001$; one sample t-test; n = 4 mice per group). Error bars represent mean with SEM. (**J, K**) Representative images of *Lcn2* mRNA upregulation in the OB of GAS-infected mice by MERFISH (**J**, yellow dots; insets) and quantification of the data (**K**). CD31 (EC marker, red) and the EC marker panel used for MERFISH (white dots) label blood vessels in **J**.

*Lcn2* mRNA is highly upregulated in the OB ECs after GAS infections by MERFISH (* $p < 0.05$; one sample t-test; $n = 4$ mice per group). Scale bars = 25 μm.

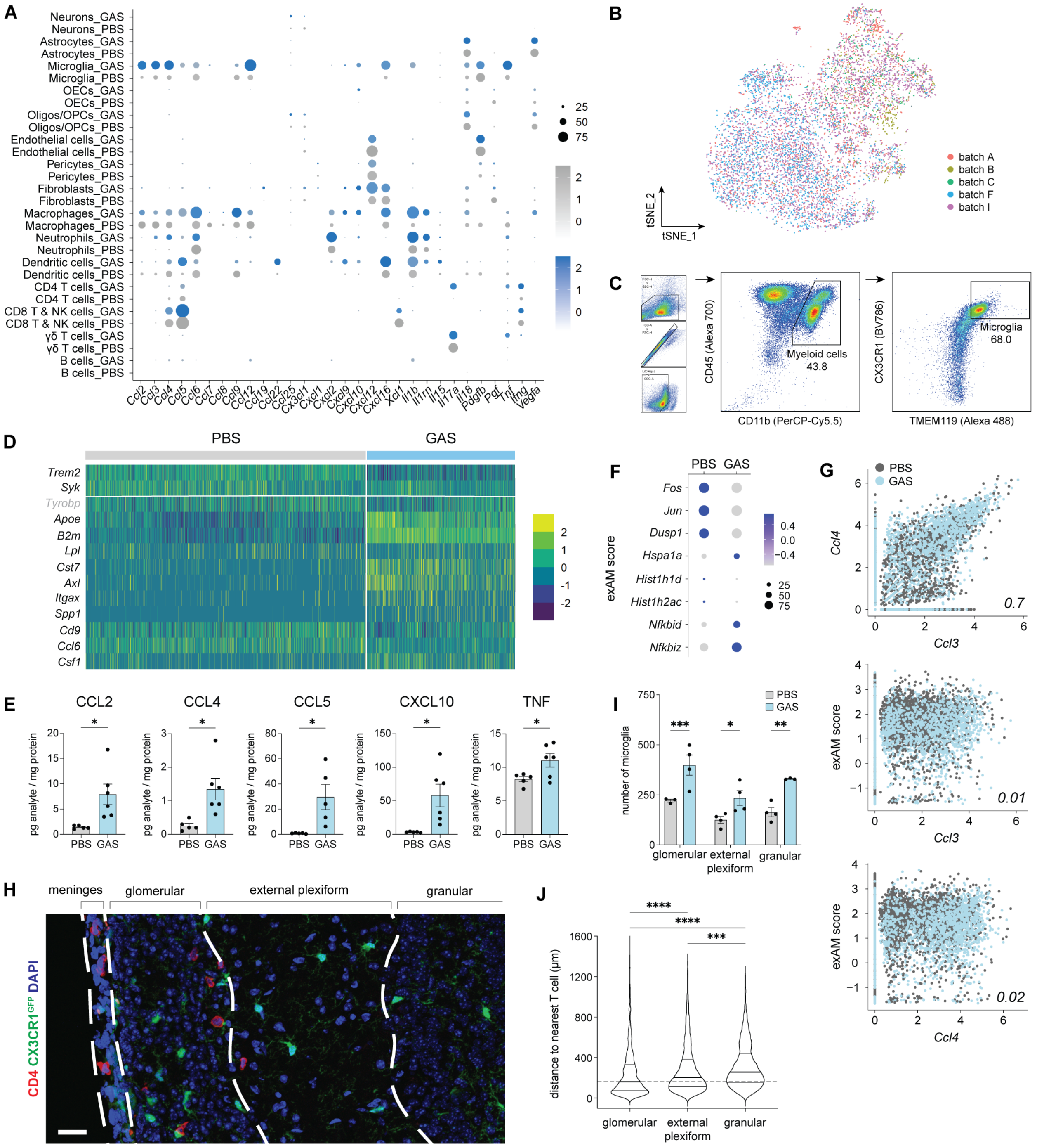


**Figure S3: Microglia upregulate disease-associated and chemokine genes after multiple GAS infections in particular where T cells are enriched in the outer OB layer, related to Figure 3**

(**A**) Dot plot of the expression levels for several cytokines, chemokines and growth factors by cell type in the OB bulb from PBS (gray) and GAS-infected (blue) mice. The dot size indicates the percent of the population expressing each marker; the color scale indicates the average level of gene expression. (**B**) *t*-SNE plot of the batch identity of microglia from PBS- and GAS-infected

OBs. (**C**) Representative gating strategy for microglia analysis (live singlets, $CD45^+$ $CD11b^+$ $CX3CR1^+$ $TMEM119^+$) by flow cytometry. (**D**) Heat map of disease-associated microglial (DAM) genes, as well as DAM regulators *Trem2* and *Syk*. Genes with gray labels are non-significant (p-adj > 0.05). (**E**) Dotted bar graphs for select cytokine levels (pg/mg) in whole OB by multiplex immunoassay. Comparisons between PBS and GAS-infected mice were done by student t-test [* $p < 0.05$; n = 5 (PBS) and n = 6 (GAS)]. (**F**) Expression of *ex vivo* activation ("exAM") genes in microglia isolated from PBS and GAS-infected OBs. The dot size indicates the percent of population expressing each marker; the color scale indicates the average level of gene expression. (**G**) Scatter plots of *Ccl3* and *Ccl4* mRNA expression by microglia (X-axis) with module score for *ex vivo* activation (exAM score; Y-axis). The Pearson correlation for each comparison is displayed in the lower right corner of the plot. (**H**) Immunofluorescence image of $CD4^+$ T cells (red) and microglia (green) localization in each layer of the OB in $CX3CR1^{eGFP}$ mice after multiple intranasal GAS infections; DAPI (blue) marks the nuclei. T cells are predominantly located in the meninges and the glomerular layer of the OB. (**I, J**) The number of microglia (**I**) and the distance from the nearest T cell (**J**) in the OB after GAS infections as assessed by MERFISH. Comparisons in **I** between PBS and GAS-infected mice were done by one-way ANOVA (* $p < 0.05$; ** $p < 0.01$; *** $p < 0.001$; n = 4 mice per group).

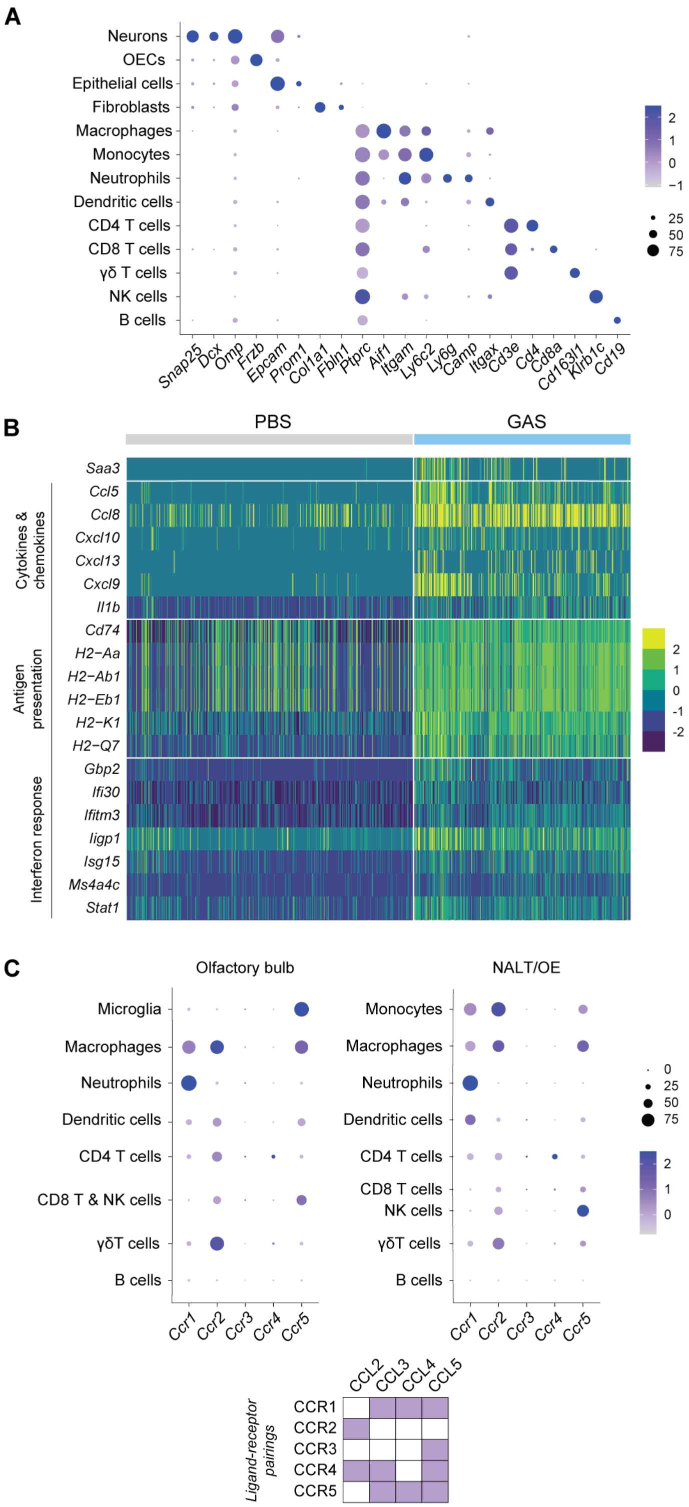


**Figure S4: Identification of NALT and OE cell types and measurement of transcriptional shifts in perivascular macrophages after intranasal GAS infections, related to Figure 4**

(**A**) Molecular markers used to assign cell identity in the nose associated lymphoid tissue (NALT) and olfactory epithelium (OE) in scRNAseq experiments. The dot size indicates the percentage of the population expressing each marker; the color scale indicates the average level of expression. (**B**) Heat map of genes related to antigen presentation, cytokines and chemokines, and interferon response, as well as the most upregulated gene (*Saa3*) in perivascular macrophages from the OB. (**C**) Expression of chemokine receptors in immune cell clusters isolated from the OB (left) and NALT/OE (right). The dot size indicates the percent of population expressing each marker; the color scale indicates the average level of expression. The purple color in the grid indicates receptor-ligand binding pairs for the relevant GAS-induced chemokines in mouse microglia / macrophages.

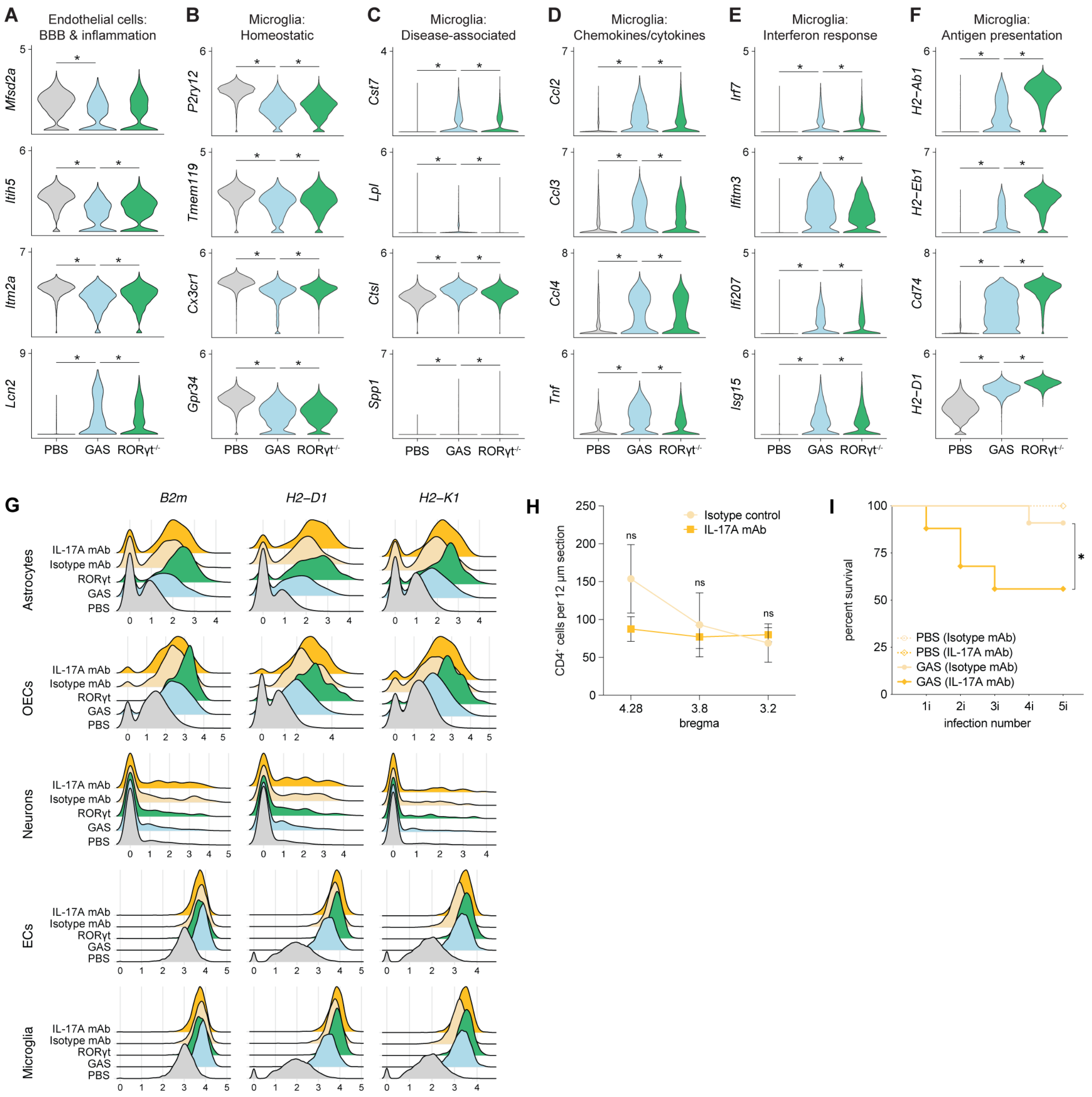


**Figure S5: Elimination of Th17 cells or blockade of IL-17A rescues BBB transcriptome changes in CNS ECs, and reduces expression of chemokine and interferon response genes by microglia in GAS-infected mice, related to Figure 5**

(**A-F**) Gene expression changes in endothelial cells (**A**) and microglia (**B-F**) isolated from the OBs of wild-type PBS (gray), wild-type GAS-infected (light blue) and RORγt$^{-/-}$ GAS-infected (green) mice. The displayed comparisons between PBS and GAS and wild-type GAS and RORγt$^{-/-}$ GAS were significant by Wilcoxon Rank Sum test with Bonferroni correction (* p-adj < 0.05). The comparisons between PBS and RORγt$^{-/-}$ GAS are not shown. (**G**) Ridge plots showing expression of key major histocompatibility complex (MHC) class I genes for antigen presentation by various

OB cell types in wild-type PBS (gray), wild-type GAS-infected mice (blue), RORγt$^{-/-}$ GAS-infected mice (green), isotype control antibody treated GAS-infected mice (yellow), and α-IL-17A mAb-treated GAS infected mice (orange). (**H**) Quantification of the number of CD4$^{+}$ T cells in the OBs of isotype-control (yellow) and α-IL-17A mAb-treated (orange) GAS infected mice (bregmas 4.28, 3.8 and 3.2). The comparison was done by mixed-effects analysis with Sidák's multiple comparisons test (ns, $p > 0.05$; n = 4 isotype-control, n = 6 α-IL-17A mAb-treated mice). (**I**) α-IL-17A mAb-treated mice showed a significant increase in mortality after GAS infection relative to isotype controls by Mantel-Cox test (* $p < 0.05$; n = 4 PBS isotype-control; n = 3 PBS α-IL-17A mAb; n = 11 GAS isotype-control and n = 19 GAS α-IL-17A mAb mice).

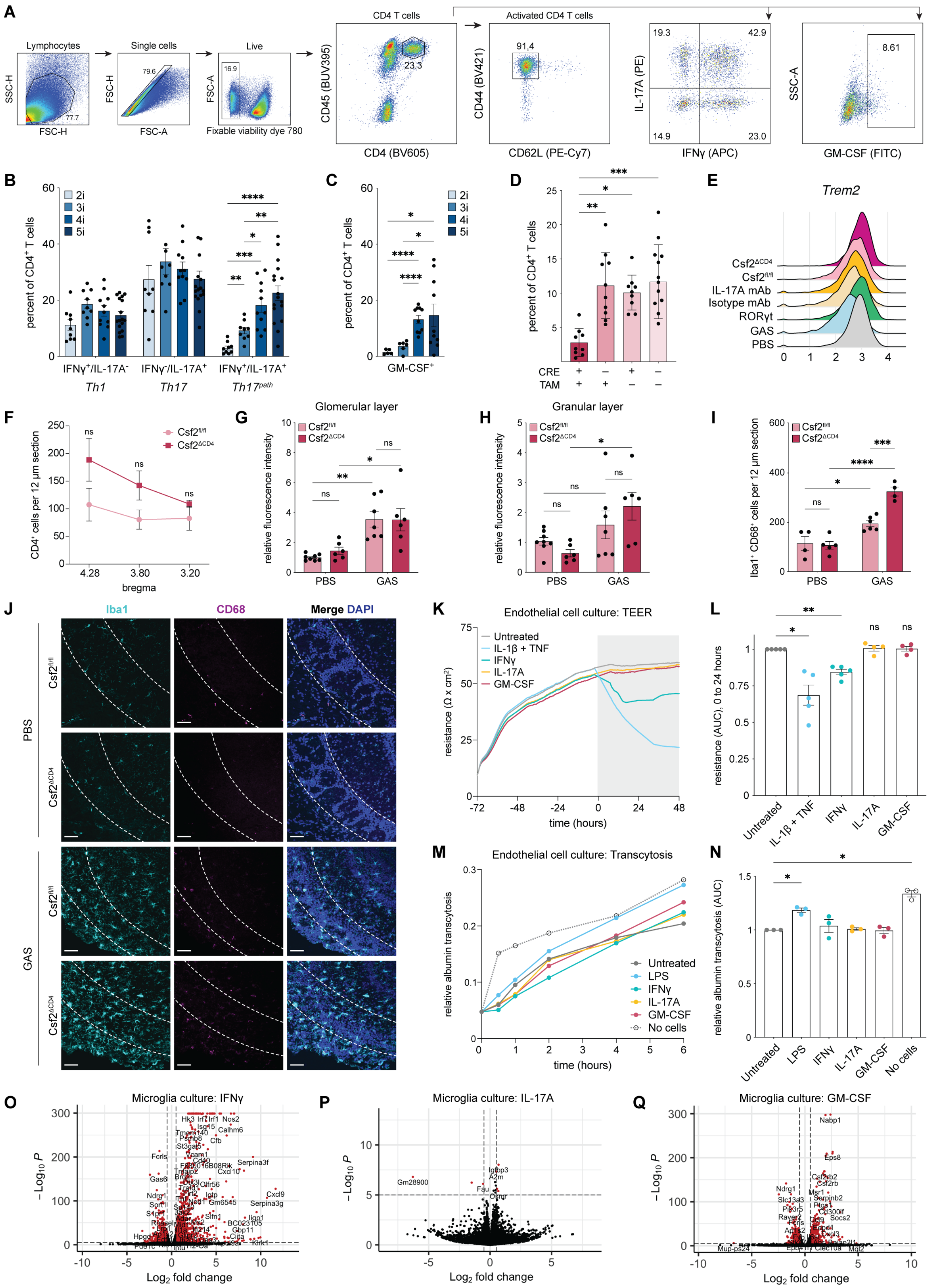

A
Lymphocytes
SSC-H
FSC-H
77.7
Single cells
FSC-H
FSC-A
79.6
Live
FSC-A
16.9
Fixable viability dye 780
CD4 T cells
CD45 (BUV395)
CD4 (BV605)
23.3
Activated CD4 T cells
CD44 (BV421)
CD62L (PE-Cy7)
91,4
IL-17A (PE)
IFNγ (APC)
19.3
42.9
14.9
23.0
SSC-A
GM-CSF (FITC)
8.61
B
percent of CD4+ T cells
2i
3i
4i
5i
IFNγ+/IL-17A-
Th1
IFNγ-/IL-17A+
Th17
IFNγ+/IL-17A+
Th17path
C
percent of CD4+ T cells
GM-CSF+
D
percent of CD4+ T cells
CRE
TAM
E
Trem2
Csf2ΔCD4
Csf2fl/fl
IL-17A mAb
Isotype mAb
RORγt
GAS
PBS
F
CD4+ cells per 12 μm section
Csf2fl/fl
Csf2ΔCD4
bregma
4.28
3.80
3.20
G
Glomerular layer
relative fluorescence intensity
PBS
GAS
H
Granular layer
relative fluorescence intensity
I
Iba1+ CD68+ cells per 12 μm section
J
Iba1
CD68
Merge DAPI
PBS
GAS
Csf2fl/fl
Csf2ΔCD4
K
Endothelial cell culture: TEER
resistance (Ω x cm2)
time (hours)
Untreated
IL-1β + TNF
IFNγ
IL-17A
GM-CSF
L
resistance (AUC), 0 to 24 hours
Untreated
IL-1β + TNF
IFNγ
IL-17A
GM-CSF
M
Endothelial cell culture: Transcytosis
relative albumin transcytosis
time (hours)
Untreated
LPS
IFNγ
IL-17A
GM-CSF
No cells
N
relative albumin transcytosis (AUC)
Untreated
LPS
IFNγ
IL-17A
GM-CSF
No cells
O
Microglia culture: IFNγ
−Log10 P
Log2 fold change
P
Microglia culture: IL-17A
Gm28900
Igfbp3
A2m
Fau
Q
Microglia culture: GM-CSF
Nabp1
Eps8

**Figure S6: Activated microglia are upregulated in the Csf2$^{\Delta CD4}$ mutant OBs following GAS infections, and effects of T cell-derived cytokines on microglia and endothelial cells *in vitro*, related to Figure 6**

(**A**) Representative gating for flow cytometry on CD4$^{+}$ T cells. (**B, C**) Dotted bar graph of the proportion of IFNγ$^{+}$IL-17A$^{+}$ CD4$^{+}$ T cells (**B**) and GM-CSF$^{+}$ T cells (**C**) in the nose associated lymphoid tissue (NALT) after 2-5 GAS infections. Cells were gated on live singlet CD45$^{+}$CD4$^{+}$ cells. The number of these T cell subtypes increases with the number of intranasal GAS infections (The comparison was done by one-way ANOVA with Dunnett's T3 multiple comparisons test; ns $p > 0.05$; * $p < 0.05$, *** $p < 0.001$; **** $p < 0.0001$; n = 9-12 mice / group). (**D**) Confirmation of CD4-specific GM-CSF knockout by flow cytometry (cells were gated on live singlet CD45$^{+}$CD4$^{+}$ cells). NALT from Csf2$^{\Delta CD4}$ mice (CRE+) that received 4-OH-tamoxifen (TAM+) prior to intranasal GAS infections (Figure 5E) had significantly fewer GM-CSF$^{+}$ CD4$^{+}$ T cells. Comparisons were done by two-way ANOVA with Sidak's multiple comparisons test (ns, $p > 0.05$; * $p < 0.05$, *** $p < 0.001$; **** $p < 0.0001$; n = 9-12 mice / group). (**E**) Ridge plot showing expression of *Trem2* in PBS (gray), wild-type GAS (blue), RORγt$^{-/-}$ GAS (green), isotype control antibody GAS (pale yellow), α-IL-17A mAb-treated GAS conditions (orange), Csf2$^{fl/fl}$ GAS (pink) and Csf2$^{\Delta CD4}$ GAS (maroon) mice. (**F**) Quantification of CD4$^{+}$ T cell numbers in the OBs (bregmas 4.28, 3.8 and 3.2) of PBS and GAS-infected Csf2$^{fl/fl}$ and Csf2$^{\Delta CD4}$ mice. The comparison was done by mixed-effects analysis with Sidák's multiple comparisons test (ns, $p > 0.05$; * $p < 0.05$; n = 6 mice / group). (**G-H**) Quantification of serum IgG leakage in the glomerular (**G**) and granular (**H**) layers of the OB in PBS and GAS-infected Csf2$^{fl/fl}$ and Csf2$^{\Delta CD4}$ mice. Statistical analyses were done by two-way ANOVA with Tukey's multiple comparison test (ns, $p > 0.05$; * $p < 0.05$; ** $p < 0.01$; n = 4-7 mice / group). (**I**) Quantification and (**J**) representative images of Iba1$^{+}$CD68$^{+}$ myeloid cells in the glomerular layer of the OB (dashed outline) in PBS and GAS-infected Csf2$^{fl/fl}$ and Csf2$^{\Delta CD4}$ mice. Statistical analyses were done with two-way ANOVA with Tukey's multiple comparison test (ns, $p > 0.05$; * $p < 0.05$, *** $p < 0.001$; **** $p < 0.0001$; n = 4-6 mice / group). (**K-N**) Effects of T cell-derived cytokines on cultured primary mouse brain endothelial cells (mBECs). Representative plot of the transendothelial electrical resistance (TEER) measurement (**K**) and quantification of independent experiments (**L**) in mBECs as a readout of changes in paracellular permeability. Representative plot (**M**) and quantification of independent experiments (**N**) of transcytosis of fluorescently tagged albumin after 24 hours of cytokine treatment in mBECs as a readout of transcellular permeability. T cell-derived cytokines (IL-17A and GM-CSF) have no direct effect on mBEC permeability *in vitro*. All comparisons were done by one-way ANOVA with Tukey's multiple comparisons test (* $p < 0.05$; ** $p < 0.01$, *** $p < 0.001$, **** $p < 0.0001$; n = 3-5 experiments per condition). (**O-Q**) Volcano plots showing significant upregulated (right) and downregulated (left) genes by bulk RNA sequencing in cultured primary brain microglia after 24 hours of treatment with IFNγ (**O**), IL-17A (**P**), or GM-CSF (**Q**) cytokines.

##### Table S10: Patient demographic data, related to Table 1

Summary of patient demographic data used for serum cytokine analysis.

| | Mean age (± SD) | Sex | | Source[a] | |
|---|---|---|---|---|---|
| | | Female | Male | CUIMC | NIMH |
| **Total** | 8.7 (± 2.6) | 19 / 34 | 15 / 34 | 13 / 34 | 21 / 34 |
| **Healthy controls** | 8.7 (± 2.2) | 4 / 11 | 7 / 11 | 0 / 11 | 11 / 11 |
| **PANDAS/PANS** | 8.7 (± 2.8) | 11 / 23 | 12 / 23 | 13 / 23 | 10 / 23 |

Sample IDs and demographic information for patient cases and controls.

| Type | Sample ID[b] | Age | Sex | Source[a] |
|---|---|---|---|---|
| Healthy control | CF3 | 7.2 | Female | NIMH |
| Healthy control | CF4 | 7.6 | Female | NIMH |
| Healthy control | CF5 | 9.9 | Female | NIMH |
| Healthy control | CF7 | 12.8 | Female | NIMH |
| Healthy control | CM1 | 5.6 | Male | NIMH |
| Healthy control | CM2 | 6.8 | Male | NIMH |
| Healthy control | CM3 | 7 | Male | NIMH |
| Healthy control | CM4 | 8.3 | Male | NIMH |
| Healthy control | CM5 | 8.4 | Male | NIMH |
| Healthy control | CM6 | 10.3 | Male | NIMH |
| Healthy control | CM7 | 11.5 | Male | NIMH |
| PANDAS/PANS | PF1 | 6 | Female | CUIMC |
| PANDAS/PANS | PF4 | 8 | Female | CUIMC |
| PANDAS/PANS | PF8 | 9 | Female | CUIMC |
| PANDAS/PANS | PF9 | 9 | Female | CUIMC |
| PANDAS/PANS | PF10 | 9 | Female | CUIMC |
| PANDAS/PANS | PF2 (a,b) | 6.4 | Female | NIMH |
| PANDAS/PANS | PF3 (a,b) | 6.8 | Female | NIMH |
| PANDAS/PANS | PF5 (a,b) | 8 | Female | NIMH |
| PANDAS/PANS | PF7 (a,b) | 8.8 | Female | NIMH |
| PANDAS/PANS | PF11 (a,b) | 10.3 | Female | NIMH |
| PANDAS/PANS | PF12 | 10.6 | Female | NIMH |
| PANDAS/PANS | PM1 | 4 | Male | CUIMC |
| PANDAS/PANS | PM2 | 5 | Male | CUIMC |
| PANDAS/PANS | PM4 | 7 | Male | CUIMC |
| PANDAS/PANS | PM6 | 8 | Male | CUIMC |
| PANDAS/PANS | PM7 | 8 | Male | CUIMC |
| PANDAS/PANS | PM13 | 13 | Male | CUIMC |
| PANDAS/PANS | PM14 | 14 | Male | CUIMC |

[a] Abbreviations: Columbia University Irving Medical Center (CUIMC) and National Institute of Mental Health (NIMH)
[b] We obtained two separate serum samples (designated "a" and "b") for 9 of the 10 NIMH cases, which had been collected at a four- to six-week interval during their published IVIg trial (Williams et al., 2016). Overall, little difference was observed for the two timepoints in both the cytokine expression and effects of sera on HUVEC cells described in this study (Table 1 and Figure 4).

| | | | | |
|---|---|---|---|---|
| PANDAS/PANS | PM15 | 15 | Male | CUIMC |
| PANDAS/PANS | PM3 (a,b) | 5.7 | Male | NIMH |
| PANDAS/PANS | PM5 (a,b) | 7.4 | Male | NIMH |
| PANDAS/PANS | PM8 (a,b) | 8.9 | Male | NIMH |
| PANDAS/PANS | PM11 (a,b) | 12 | Male | NIMH |

#### Table S11: *Il17ra* expression in mouse BECs and microglia, related to Figure S6

| **Sequencing type** | **Source** | **Cell type** | ***Il17ra* expression**[c] |
|---|---|---|---|
| scRNAseq | Olfactory bulb | Microglia | 88.3 CPM |
| scRNAseq | Olfactory bulb | Endothelial cells | 17.6 CPM |
| bulk RNAseq | Primary culture | Microglia | O TPM |
| bulk RNAseq[d] | Primary culture | Endothelial cells | 1.6 TPM |

[c] Abbreviations: Reads per kilobase million (RPKM); transcripts per million (TPM)
[d] Endothelial cell bulk RNAseq values obtained by personal correspondence

#### Document S2: Scripts for processing of scRNAseq data

```
# analysis run with R v 4.0.2 and Seurat 4.0.2

# load programs
library(Seurat)
library(dplyr)
library(ggplot2)

## code for scRNAseq analysis of mouse olfactory bulb after GAS infection ----

# import data files & merge
ds004.data <- Read10X(data.dir = "Directory/CW004/raw_feature_bc_matrix")
ds005.data <- Read10X(data.dir = "Directory/CW005/raw_feature_bc_matrix")
ds008.data <- Read10X(data.dir = "Directory/CW008/raw_feature_bc_matrix")
ds009.data <- Read10X(data.dir = "Directory/CW009/raw_feature_bc_matrix")
ds010.data <- Read10X(data.dir = "Directory/CW010/raw_feature_bc_matrix")
ds011.data <- Read10X(data.dir = "Directory/CW011/raw_feature_bc_matrix")
ds012.data <- Read10X(data.dir = "Directory/CW012/raw_feature_bc_matrix")
ds013.data <- Read10X(data.dir = "Directory/CW013/raw_feature_bc_matrix")
ds014.data <- Read10X(data.dir = "Directory/CW014/raw_feature_bc_matrix")
ds015.data <- Read10X(data.dir = "Directory/CW015/raw_feature_bc_matrix")
ds016.data <- Read10X(data.dir = "Directory/CW016/raw_feature_bc_matrix")
ds017.data <- Read10X(data.dir = "Directory/CW017/raw_feature_bc_matrix")
ds018.data <- Read10X(data.dir = "Directory/CW018/raw_feature_bc_matrix")
ds021.data <- Read10X(data.dir = "Directory/CW021/raw_feature_bc_matrix")
ds022.data <- Read10X(data.dir = "Directory/CW022/raw_feature_bc_matrix")
ds023.data <- Read10X(data.dir = "Directory/CW023/raw_feature_bc_matrix")
ds024.data <- Read10X(data.dir = "Directory/CW024/raw_feature_bc_matrix")
ds025.data <- Read10X(data.dir = "Directory/CW025/raw_feature_bc_matrix")
ds026.data <- Read10X(data.dir = "Directory/CW026/raw_feature_bc_matrix")

ds004 <- CreateSeuratObject(counts = ds004.data, project = "PBS", min.cells = 3,
       min.features = 200)
ds005 <- CreateSeuratObject(counts = ds005.data, project = "GAS", min.cells = 3,
       min.features = 200)
ds008 <- CreateSeuratObject(counts = ds008.data, project = "GAS", min.cells = 3,
       min.features = 200)
ds009 <- CreateSeuratObject(counts = ds009.data, project = "GAS", min.cells = 3,
       min.features = 200)
ds010 <- CreateSeuratObject(counts = ds010.data, project = "RORg", min.cells = 3,
       min.features = 200)
ds011 <- CreateSeuratObject(counts = ds011.data, project = "GAS", min.cells = 3,
       min.features = 200) # Setting orig.ident on CW011 as "GAS" but condition as
       "Isotype_mAb"
ds012 <- CreateSeuratObject(counts = ds012.data, project = "IL-17A_mAb", min.cells =
       3, min.features = 200)
ds013 <- CreateSeuratObject(counts = ds013.data, project = "RORg", min.cells = 3,
       min.features = 200)
ds014 <- CreateSeuratObject(counts = ds014.data, project = "PBS", min.cells = 3,
       min.features = 200)
ds015 <- CreateSeuratObject(counts = ds015.data, project = "GAS", min.cells = 3,
       min.features = 200) # Setting orig.ident on CW015 as "GAS" but condition as
       "Isotype_mAb"
ds016 <- CreateSeuratObject(counts = ds016.data, project = "IL-17A_mAb", min.cells =
       3, min.features = 200)
ds017 <- CreateSeuratObject(counts = ds017.data, project = "IL-17A_mAb", min.cells =
       3, min.features = 200)
ds018 <- CreateSeuratObject(counts = ds018.data, project = "RORg", min.cells = 3,
       min.features = 200)
ds021 <- CreateSeuratObject(counts = ds021.data, project = "GAS", min.cells = 3,
       min.features = 200) # Setting orig.ident on CW021 as "GAS" but condition as
       "Csf2flox"
ds022 <- CreateSeuratObject(counts = ds022.data, project = "Csf2_CD4ko", min.cells =
       3, min.features = 200)
```

```
ds023 <- CreateSeuratObject(counts = ds023.data, project = "PBS", min.cells = 3,
       min.features = 200)
ds024 <- CreateSeuratObject(counts = ds024.data, project = "GAS", min.cells = 3,
       min.features = 200)
ds025 <- CreateSeuratObject(counts = ds025.data, project = "GAS", min.cells = 3,
       min.features = 200) # Setting orig.ident on CW025 as "GAS" but condition as
       "Csf2flox"
ds026 <- CreateSeuratObject(counts = ds026.data, project = "Csf2_CD4ko", min.cells =
       3, min.features = 200)

dsmerged <- merge(ds004, y = c(ds005, ds008, ds009, ds010, ds011, ds012, ds013, ds014,
       ds015, ds016, ds017, ds018, ds021, ds022, ds023, ds024, ds025, ds026),
       add.cell.ids = c("CW_004", "CW_005", "CW_008", "CW_009", "CW_010", "CW_011",
       "CW_012", "CW_013", "CW_014", "CW_015", "CW_016", "CW_017", "CW_018", "CW_021",
       "CW_022", "CW_023", "CW_024", "CW_025", "CW_026"))

rm(ds004, ds004.data, ds005, ds005.data, ds008, ds008.data, ds009, ds009.data, ds010,
       ds010.data, ds011, ds011.data, ds012, ds012.data, ds013, ds013.data, ds014,
       ds014.data, ds015, ds015.data, ds016, ds016.data, ds017, ds017.data, ds018,
       ds018.data, ds021, ds021.data, ds022, ds022.data, ds023, ds023.data, ds024,
       ds024.data, ds025, ds025.data, ds026, ds026.data)

# add meta data

# add sample to meta data
samplevec <- rep("CW004",nrow(dsmerged@meta.data))
samplevec[grep("CW_005",rownames(dsmerged@meta.data))] <- "CW005"
samplevec[grep("CW_008",rownames(dsmerged@meta.data))] <- "CW008"
samplevec[grep("CW_009",rownames(dsmerged@meta.data))] <- "CW009"
samplevec[grep("CW_010",rownames(dsmerged@meta.data))] <- "CW010"
samplevec[grep("CW_011",rownames(dsmerged@meta.data))] <- "CW011"
samplevec[grep("CW_012",rownames(dsmerged@meta.data))] <- "CW012"
samplevec[grep("CW_013",rownames(dsmerged@meta.data))] <- "CW013"
samplevec[grep("CW_014",rownames(dsmerged@meta.data))] <- "CW014"
samplevec[grep("CW_015",rownames(dsmerged@meta.data))] <- "CW015"
samplevec[grep("CW_016",rownames(dsmerged@meta.data))] <- "CW016"
samplevec[grep("CW_017",rownames(dsmerged@meta.data))] <- "CW017"
samplevec[grep("CW_018",rownames(dsmerged@meta.data))] <- "CW018"
samplevec[grep("CW_021",rownames(dsmerged@meta.data))] <- "CW021"
samplevec[grep("CW_022",rownames(dsmerged@meta.data))] <- "CW022"
samplevec[grep("CW_023",rownames(dsmerged@meta.data))] <- "CW023"
samplevec[grep("CW_024",rownames(dsmerged@meta.data))] <- "CW024"
samplevec[grep("CW_025",rownames(dsmerged@meta.data))] <- "CW025"
samplevec[grep("CW_026",rownames(dsmerged@meta.data))] <- "CW026"
table(samplevec)
dsmerged@meta.data$sample=samplevec

# add batch to meta data
batchvec <- rep("batch A",nrow(dsmerged@meta.data))
batchvec[grep("CW_008",rownames(dsmerged@meta.data))] <- "batch B"
batchvec[grep("CW_009",rownames(dsmerged@meta.data))] <- "batch C"
batchvec[grep("CW_010",rownames(dsmerged@meta.data))] <- "batch C"
batchvec[grep("CW_011",rownames(dsmerged@meta.data))] <- "batch D"
batchvec[grep("CW_012",rownames(dsmerged@meta.data))] <- "batch D"
batchvec[grep("CW_013",rownames(dsmerged@meta.data))] <- "batch E"
batchvec[grep("CW_014",rownames(dsmerged@meta.data))] <- "batch F"
batchvec[grep("CW_015",rownames(dsmerged@meta.data))] <- "batch F"
batchvec[grep("CW_016",rownames(dsmerged@meta.data))] <- "batch F"
batchvec[grep("CW_017",rownames(dsmerged@meta.data))] <- "batch G"
batchvec[grep("CW_021",rownames(dsmerged@meta.data))] <- "batch H"
batchvec[grep("CW_022",rownames(dsmerged@meta.data))] <- "batch H"
batchvec[grep("CW_023",rownames(dsmerged@meta.data))] <- "batch I"
batchvec[grep("CW_024",rownames(dsmerged@meta.data))] <- "batch I"
batchvec[grep("CW_018",rownames(dsmerged@meta.data))] <- "batch J"
batchvec[grep("CW_025",rownames(dsmerged@meta.data))] <- "batch K"
batchvec[grep("CW_026",rownames(dsmerged@meta.data))] <- "batch K"
```

```
table(batchvec)
dsmerged@meta.data$batch=batchvec

# add enrichment to meta data
enrichmentvec <- rep("cd31_cd11b_FACS",nrow(dsmerged@meta.data))
enrichmentvec[grep("CW_008",rownames(dsmerged@meta.data))] <- "live_FACS"
enrichmentvec[grep("CW_023",rownames(dsmerged@meta.data))] <- "live_FACS"
enrichmentvec[grep("CW_024",rownames(dsmerged@meta.data))] <- "live_FACS"
table(enrichmentvec)
dsmerged@meta.data$enrichment=enrichmentvec

# add inoculate to meta data
inoculatevec <- rep("GAS",nrow(dsmerged@meta.data))
inoculatevec[grep("CW_004",rownames(dsmerged@meta.data))] <- "PBS"
inoculatevec[grep("CW_014",rownames(dsmerged@meta.data))] <- "PBS"
inoculatevec[grep("CW_023",rownames(dsmerged@meta.data))] <- "PBS"
table(inoculatevec)
dsmerged@meta.data$inoculate=inoculatevec

# add tissue to meta data
tissuevec <- rep("OB",nrow(dsmerged@meta.data))
table(tissuevec)
dsmerged@meta.data$tissue=tissuevec

# add genotype to meta data
genotypevec <- rep("WT",nrow(dsmerged@meta.data))
genotypevec[grep("CW_010",rownames(dsmerged@meta.data))] <- "RORg"
genotypevec[grep("CW_013",rownames(dsmerged@meta.data))] <- "RORg"
genotypevec[grep("CW_018",rownames(dsmerged@meta.data))] <- "RORg"
genotypevec[grep("CW_021",rownames(dsmerged@meta.data))] <- "Csf2flox"
genotypevec[grep("CW_022",rownames(dsmerged@meta.data))] <- "Csf2floxCD4Cre"
genotypevec[grep("CW_025",rownames(dsmerged@meta.data))] <- "Csf2flox"
genotypevec[grep("CW_026",rownames(dsmerged@meta.data))] <- "Csf2floxCD4Cre"
table(genotypevec)
dsmerged@meta.data$genotype=genotypevec

# add drug to meta data
drugvec <- rep("none",nrow(dsmerged@meta.data))
drugvec[grep("CW_011",rownames(dsmerged@meta.data))] <- "Isotype_mAb"
drugvec[grep("CW_012",rownames(dsmerged@meta.data))] <- "IL-17A_mAb"
drugvec[grep("CW_015",rownames(dsmerged@meta.data))] <- "Isotype_mAb"
drugvec[grep("CW_016",rownames(dsmerged@meta.data))] <- "IL-17A_mAb"
drugvec[grep("CW_017",rownames(dsmerged@meta.data))] <- "IL-17A_mAb"
drugvec[grep("CW_021",rownames(dsmerged@meta.data))] <- "4-OH-tamoxifen"
drugvec[grep("CW_022",rownames(dsmerged@meta.data))] <- "4-OH-tamoxifen"
drugvec[grep("CW_025",rownames(dsmerged@meta.data))] <- "4-OH-tamoxifen"
drugvec[grep("CW_026",rownames(dsmerged@meta.data))] <- "4-OH-tamoxifen"
table(drugvec)
dsmerged@meta.data$drug=drugvec

# add condition to meta data
conditionvec <- rep("GAS",nrow(dsmerged@meta.data))
conditionvec[grep("CW_004",rownames(dsmerged@meta.data))] <- "PBS"
conditionvec[grep("CW_010",rownames(dsmerged@meta.data))] <- "RORg"
conditionvec[grep("CW_011",rownames(dsmerged@meta.data))] <- "Isotype_mAb"
conditionvec[grep("CW_012",rownames(dsmerged@meta.data))] <- "IL-17A_mAb"
conditionvec[grep("CW_013",rownames(dsmerged@meta.data))] <- "RORg"
conditionvec[grep("CW_014",rownames(dsmerged@meta.data))] <- "PBS"
conditionvec[grep("CW_015",rownames(dsmerged@meta.data))] <- "Isotype_mAb"
conditionvec[grep("CW_016",rownames(dsmerged@meta.data))] <- "IL-17A_mAb"
conditionvec[grep("CW_017",rownames(dsmerged@meta.data))] <- "IL-17A_mAb"
conditionvec[grep("CW_018",rownames(dsmerged@meta.data))] <- "RORg"
conditionvec[grep("CW_021",rownames(dsmerged@meta.data))] <- "Csf2flox"
conditionvec[grep("CW_022",rownames(dsmerged@meta.data))] <- "Csf2_CD4ko"
conditionvec[grep("CW_023",rownames(dsmerged@meta.data))] <- "PBS"
```

```
conditionvec[grep("CW_025",rownames(dsmerged@meta.data))] <- "Csf2flox"
conditionvec[grep("CW_026",rownames(dsmerged@meta.data))] <- "Csf2_CD4ko"
table(conditionvec)
dsmerged@meta.data$condition=conditionvec


# remove dead cells

dsmerged[["percent.mt"]] <- PercentageFeatureSet(dsmerged, pattern = "^mt-")
VlnPlot(dsmerged, features = c("nFeature_RNA", "nCount_RNA", "percent.mt"), ncol = 3)
dsmerged <- subset(dsmerged, subset = nCount_RNA > 1000 & nCount_RNA < 50000 &
       percent.mt < 20)
VlnPlot(dsmerged, features = c("nFeature_RNA", "nCount_RNA", "percent.mt"), ncol = 3)

# normalize & generate PCA
dsmerged <- NormalizeData(dsmerged)
dsmerged <- FindVariableFeatures(dsmerged, selection.method = "vst", nfeatures = 3000)
top10 <- head(VariableFeatures(dsmerged), 10)
plot1 <- VariableFeaturePlot(dsmerged)
plot2 <- LabelPoints(plot = plot1, points = top10, repel = TRUE, xnudge = 0, ynudge =
       0)
plot2
all.genes <- rownames(dsmerged)
dsmerged <- ScaleData(dsmerged, features = all.genes)
dsmerged <- RunPCA(dsmerged, features = VariableFeatures(object = dsmerged))
print(dsmerged[["pca"]], dims = 1:5, nfeatures = 5)
VizDimLoadings(dsmerged, dims = 1:5, reduction = "pca")
ElbowPlot(dsmerged,ndims=50)
dsmerged <- FindNeighbors(dsmerged, dims = 1:39)
dsmerged <- FindClusters(dsmerged, resolution = 1)
dsmerged <- RunTSNE(dsmerged, dims = 1:39)
DimPlot(dsmerged, reduction = "tsne",label=T)
DimPlot(dsmerged, reduction = "tsne",group.by="orig.ident")
rm(plot1, plot2, all.genes, top10)

# run harmony to reduce batch effects

library(harmony)
library(cowplot)

DimPlot(dsmerged, reduction = "tsne",group.by="batch")

dsmerged <- dsmerged %>%
RunHarmony("batch", plot_convergence = TRUE)

harmony_embeddings <- Embeddings(dsmerged, 'harmony')
harmony_embeddings[1:5, 1:5]

dsmerged <- dsmerged %>%
RunTSNE(reduction = "harmony", dims = 1:39) %>%
FindNeighbors(reduction = "harmony", dims = 1:39) %>%
FindClusters(resolution = 1) %>%
identity()

DimPlot(dsmerged, reduction = "tsne",label=T)
DimPlot(dsmerged, reduction = "tsne",group.by="batch")


# identify cell types

neuroglial.markers <- c("Snap25", "Dcx", "Gfap","Aqp4", "Frzb","Cldn5", "Pecam1",
       "Pdgfra","Pecam1","Pdgfrb", "Atp13a5", "Col1a1", "Fbln1")
lymphoid.markers <- c("Ptprc", "Cd3e", "Cd4", "Cd8a", "Cd163l1", "Klrb1c", "Cd19",
       "Ms4a1")
myeloid.markers <- c("Ptprc", "Tmem119", "Aif1", "Itgam", "Ly6g", "Camp", "Ly6c2",
       "Itgax", "Xcr1", "Cd209a", "Ccr9", "Nudt17")
```

```r
macrophage.vs.microglia.markers <- c("Ptprc", "Tmem119", "Cx3cr1", "P2ry12", "Itgam",
       "Aif1", "Hexb", "Plac8", "Mrc1", "Cd163")

# assign cluster identities
Idents(object=dsmerged) <- dsmerged@meta.data$RNA_snn_res.1
DimPlot(dsmerged, reduction = "tsne",label=T)
new.cluster.ids <- c("Macrophages", "Microglia", "Microglia", "Endothelial cells",
       "Neutrophils", "Endothelial cells", "CD4 T cells", "Microglia", "Astrocytes",
       "Olfactory ensheathing cells", "Neutrophils", "Olfactory ensheathing cells",
       "Macrophages", "Neurons", "Dendritic cells", "Pericytes", "Microglia",
       "Microglia", "CD8 T cells & NK cells", "Astrocytes", "gamma delta T cells",
       "Dendritic cells", "Macrophages", "Mixed signature: Microglia/Neutrophils",
       "Microglia", "Olfactory ensheathing cells", "Pericytes", "Neurons", "Neurons",
       "Neutrophils", "Neutrophils", "Oligos/OPCs", "B cells", "Macrophages", "Mixed
       signature: Microglia/Endothelial cells", "Endothelial cells", "Fibroblasts",
       "Mixed signature: Microglia/T cells", "Dendritic cells", "Mixed signature:
       Olfactory ensheathing cells/Endothelial cells", "Mixed signature:
       Microglia/Astrocytes", "Mixed signature: Endothelial cells/T cells", "Mixed
       signature: Macrophages/T cells")
names(new.cluster.ids) <- levels(dsmerged)
dsmerged <- RenameIdents(dsmerged, new.cluster.ids)
DimPlot(dsmerged, reduction = "tsne",label=T)+ NoLegend()
dsmerged[["cluster.names_all.cells"]] <- Idents(object = dsmerged) # stash identity


# remove mixed signature clusters
dsmerged <- subset(x = dsmerged, idents = c("Neurons", "Astrocytes", "Olfactory
       ensheathing cells", "Oligos/OPCs", "Microglia", "Endothelial cells",
       "Pericytes", "Fibroblasts", "Macrophages", "Dendritic cells", "Neutrophils",
       "CD4 T cells", "CD8 T cells & NK cells", "gamma delta T cells", "B
       cells"),invert = FALSE)
DimPlot(dsmerged, reduction = "tsne",label=T)

# subset on unenriched samples & recluster

DimPlot(dsmerged, reduction = "tsne",label=T)
DimPlot(dsmerged, reduction = "tsne",group.by="enrichment")

Idents(object= dsmerged) <- dsmerged@meta.data$sample
dsmerged_008_023_024 <- subset(x = dsmerged, idents = c("CW008", "CW023",
       "CW024"),invert = FALSE)
DimPlot(dsmerged_008_023_024, reduction = "tsne",label=T)
dsmerged_008_023_024 <- NormalizeData(dsmerged_008_023_024)
dsmerged_008_023_024 <- FindVariableFeatures(dsmerged_008_023_024, selection.method =
       "vst", nfeatures = 3000)
top10 <- head(VariableFeatures(dsmerged_008_023_024), 10)
plot1 <- VariableFeaturePlot(dsmerged_008_023_024)
plot2 <- LabelPoints(plot = plot1, points = top10, repel = TRUE, xnudge = 0, ynudge =
       0)
plot2
all.genes <- rownames(dsmerged_008_023_024)
dsmerged_008_023_024 <- ScaleData(dsmerged_008_023_024, features = all.genes)
dsmerged_008_023_024 <- RunPCA(dsmerged_008_023_024, features =
       VariableFeatures(object = dsmerged_008_023_024))
print(dsmerged_008_023_024[["pca"]], dims = 1:5, nfeatures = 5)
VizDimLoadings(dsmerged_008_023_024, dims = 1:5, reduction = "pca")
ElbowPlot(dsmerged_008_023_024,ndims=50)
dsmerged_008_023_024 <- FindNeighbors(dsmerged_008_023_024, dims = 1:42)
dsmerged_008_023_024 <- FindClusters(dsmerged_008_023_024, resolution = 1)
dsmerged_008_023_024 <- RunTSNE(dsmerged_008_023_024, dims = 1:42)
DimPlot(dsmerged_008_023_024, reduction = "tsne",label=T)

DimPlot(dsmerged_008_023_024, reduction = "tsne",group.by="orig.ident")
DimPlot(dsmerged_008_023_024, reduction = "tsne",label=F, group.by="inoculate",
       split.by = "condition")
DimPlot(dsmerged_008_023_024, reduction = "tsne",label=F, group.by="batch")
```

```
# run harmony

dsmerged_008_023_024 <- dsmerged_008_023_024 %>%
RunHarmony("batch", plot_convergence = TRUE)

harmony_embeddings <- Embeddings(dsmerged_008_023_024, 'harmony')
harmony_embeddings[1:5, 1:5]

dsmerged_008_023_024 <- dsmerged_008_023_024 %>%
RunTSNE(reduction = "harmony", dims = 1:42) %>%
FindNeighbors(reduction = "harmony", dims = 1:42) %>%
FindClusters(resolution = 1) %>%
identity()

DimPlot(dsmerged_008_023_024, reduction = "tsne",label=T)
DimPlot(dsmerged_008_023_024, reduction = "tsne",label=F, group.by="inoculate")

rm(plot1, plot2, harmony_embeddings, top10)


# assign cluster identities

DimPlot(dsmerged_008_023_024, reduction = "tsne",label=T)
new.cluster.ids <- c("Olfactory ensheathing cells", "Microglia", "Microglia",
       "Neurons", "Olfactory ensheathing cells", "Olfactory ensheathing cells",
       "Astrocytes", "CD4 & CD8 T cells", "Astrocytes", "Endothelial cells",
       "Olfactory ensheathing cells", "Olfactory ensheathing cells", "Neurons",
       "Neutrophils", "Neurons", "Macrophages", "Pericytes", "Astrocytes",
       "Oligos/OPCs", "Macrophages", "Dendritic cells", "gamma delta T cells",
       "Microglia", "Neurons", "Microglia", "Mixed signature: Multiple cell types",
       "Neutrophils", "Mixed signature: Multiple cell types", "NK cells",
       "Fibroblasts", "B cells", "Mixed signature: Neurons/Microglia", "Mixed
       signature: Astrocytes/Neurons", "Fibroblasts", "Dendritic cells", "Olfactory
       ensheathing cells", "Dendritic cells", "Mixed signature: Endothelial
       cells/Pericytes")
names(new.cluster.ids) <- levels(dsmerged_008_023_024)
dsmerged_008_023_024 <- RenameIdents(dsmerged_008_023_024, new.cluster.ids)
DimPlot(dsmerged_008_023_024, reduction = "tsne",label=T)+ NoLegend()
dsmerged_008_023_024[["cluster.names_unenriched"]] <- Idents(object =
       dsmerged_008_023_024) # stash identity

# remove mixed-signature clusters

dsmerged_008_023_024 <- subset(x = dsmerged_008_023_024, idents = c("Mixed signature:
       Multiple cell types", "Mixed signature: Neurons/Microglia", "Mixed signature:
       Astrocytes/Neurons", "Mixed signature: Endothelial cells/Pericytes"),invert =
       TRUE)
DimPlot(dsmerged_008_023_024, reduction = "tsne",label=T)+ NoLegend()
```

```
## code for scRNAseq analysis of endothelial cells after GAS infection ----

# subsetting & reclustering ECs

Idents(object=dsmerged) <- dsmerged@meta.data$cluster.names_all.cells
dsmerged_ECs <- subset(x = dsmerged, idents = c("Endothelial cells"),invert = FALSE)
DimPlot(dsmerged_ECs, reduction = "tsne",label=T)
dsmerged_ECs <- NormalizeData(dsmerged_ECs)
dsmerged_ECs <- FindVariableFeatures(dsmerged_ECs, selection.method = "vst", nfeatures
       = 3000)
top10 <- head(VariableFeatures(dsmerged_ECs), 10)
plot1 <- VariableFeaturePlot(dsmerged_ECs)
plot2 <- LabelPoints(plot = plot1, points = top10, repel = TRUE, xnudge = 0, ynudge =
       0)
plot2
all.genes <- rownames(dsmerged_ECs)
dsmerged_ECs <- ScaleData(dsmerged_ECs, features = all.genes)
dsmerged_ECs <- RunPCA(dsmerged_ECs, features = VariableFeatures(object =
       dsmerged_ECs))
print(dsmerged_ECs[["pca"]], dims = 1:5, nfeatures = 5)
VizDimLoadings(dsmerged_ECs, dims = 1:5, reduction = "pca")
ElbowPlot(dsmerged_ECs,ndims=50)
dsmerged_ECs <- FindNeighbors(dsmerged_ECs, dims = 1:38)
dsmerged_ECs <- FindClusters(dsmerged_ECs, resolution = 0.4)
dsmerged_ECs <- RunTSNE(dsmerged_ECs, dims = 1:38)
DimPlot(dsmerged_ECs, reduction = "tsne",label=T)

DimPlot(dsmerged_ECs, reduction = "tsne",label=F, group.by="batch")

# batch correction

dsmerged_ECs <- dsmerged_ECs %>%
RunHarmony("batch", plot_convergence = TRUE)

harmony_embeddings <- Embeddings(dsmerged_ECs, 'harmony')
harmony_embeddings[1:5, 1:5]

dsmerged_ECs <- dsmerged_ECs %>%
RunTSNE(reduction = "harmony", dims = 1:38) %>%
FindNeighbors(reduction = "harmony", dims = 1:38) %>%
FindClusters(resolution = 0.4) %>%
identity()

DimPlot(dsmerged_ECs, reduction = "tsne",label=F, group.by = "batch")

FeaturePlot(dsmerged_ECs, c("nCount_RNA", "nFeature_RNA", "percent.mt")) # cluster 4
       is low viability
FeaturePlot(dsmerged_ECs, c("Aqp4", "Gfap"))# cluster 5 has mixed signature:
       astrocytes/endothelial cells
FeaturePlot(dsmerged_ECs, c("Pdgfrb", "Atp13a5"))# cluster 6 has mixed signature:
       pericytes/endothelial cells
FeaturePlot(dsmerged_ECs, c("Cd3e", "Ptprc"))# cluster 7 has mixed signature:
       microglia/endothelial cells
FeaturePlot(dsmerged_ECs, c("Frzb", "Plp1"))# cluster 9 has mixed signature: olfactory
       ensheathing cells/endothelial cells

# remove low viability, mixed signature clusters

dsmerged_ECs <- subset(x = dsmerged_ECs, idents = c(4, 5, 6, 7, 9, 10),invert = TRUE)
DimPlot(dsmerged_ECs, reduction = "tsne",label=T)+ NoLegend()

rm(plot1, plot2, harmony_embeddings, all.genes, top10)


# subset and recluster PBS & GAS ECs
```

```r
Idents(object=dsmerged_ECs) <- dsmerged_ECs@meta.data$condition
DimPlot(dsmerged_ECs, reduction = "tsne",label=T)
dsmerged_ECs_PBS_GAS <- subset(x = dsmerged_ECs, idents = c("PBS", "GAS"),invert =
       FALSE)
DimPlot(dsmerged_ECs_PBS_GAS, reduction = "tsne",label=T)
dsmerged_ECs_PBS_GAS <- NormalizeData(dsmerged_ECs_PBS_GAS)
dsmerged_ECs_PBS_GAS <- FindVariableFeatures(dsmerged_ECs_PBS_GAS, selection.method =
       "vst", nfeatures = 3000)
top10 <- head(VariableFeatures(dsmerged_ECs_PBS_GAS), 10)
plot1 <- VariableFeaturePlot(dsmerged_ECs_PBS_GAS)
plot2 <- LabelPoints(plot = plot1, points = top10, repel = TRUE, xnudge = 0, ynudge =
       0)
plot2
all.genes <- rownames(dsmerged_ECs_PBS_GAS)
dsmerged_ECs_PBS_GAS <- ScaleData(dsmerged_ECs_PBS_GAS, features = all.genes)
dsmerged_ECs_PBS_GAS <- RunPCA(dsmerged_ECs_PBS_GAS, features =
       VariableFeatures(object = dsmerged_ECs_PBS_GAS))
print(dsmerged_ECs_PBS_GAS[["pca"]], dims = 1:5, nfeatures = 5)
VizDimLoadings(dsmerged_ECs_PBS_GAS, dims = 1:5, reduction = "pca")
ElbowPlot(dsmerged_ECs_PBS_GAS,ndims=50)
dsmerged_ECs_PBS_GAS <- FindNeighbors(dsmerged_ECs_PBS_GAS, dims = 1:36)
dsmerged_ECs_PBS_GAS <- FindClusters(dsmerged_ECs_PBS_GAS, resolution = 0.4)
dsmerged_ECs_PBS_GAS <- RunTSNE(dsmerged_ECs_PBS_GAS, dims = 1:36)
DimPlot(dsmerged_ECs_PBS_GAS, reduction = "tsne",label=T)

DimPlot(dsmerged_ECs_PBS_GAS, reduction = "tsne",group.by="orig.ident")
DimPlot(dsmerged_ECs_PBS_GAS, reduction = "tsne",label=F, group.by="inoculate",
       split.by = "condition")
DimPlot(dsmerged_ECs_PBS_GAS, reduction = "tsne",label=F, group.by="batch")

# run harmony

dsmerged_ECs_PBS_GAS <- dsmerged_ECs_PBS_GAS %>%
RunHarmony("batch", plot_convergence = TRUE)

harmony_embeddings <- Embeddings(dsmerged_ECs_PBS_GAS, 'harmony')
harmony_embeddings[1:5, 1:5]

dsmerged_ECs_PBS_GAS <- dsmerged_ECs_PBS_GAS %>%
RunTSNE(reduction = "harmony", dims = 1:36) %>%
FindNeighbors(reduction = "harmony", dims = 1:36) %>%
FindClusters(resolution = 0.4) %>%
identity()

DimPlot(dsmerged_ECs_PBS_GAS, reduction = "tsne",label=F, group.by="batch")

rm(plot1, plot2, harmony_embeddings, all.genes, top10)
```

```r
## code for scRNAseq analysis of microglia after GAS infection ----

# subsetting & reclustering microglia

Idents(object=dsmerged) <- dsmerged@meta.data$cluster.names_all.cells
dsmerged_microglia <- subset(x = dsmerged, idents = c("Microglia"),invert = FALSE)
DimPlot(dsmerged_microglia, reduction = "tsne",label=T)
dsmerged_microglia <- NormalizeData(dsmerged_microglia)
dsmerged_microglia <- FindVariableFeatures(dsmerged_microglia, selection.method =
       "vst", nfeatures = 3000)
top10 <- head(VariableFeatures(dsmerged_microglia), 10)
plot1 <- VariableFeaturePlot(dsmerged_microglia)
plot2 <- LabelPoints(plot = plot1, points = top10, repel = TRUE, xnudge = 0, ynudge =
       0)
plot2
all.genes <- rownames(dsmerged_microglia)
dsmerged_microglia <- ScaleData(dsmerged_microglia, features = all.genes)
dsmerged_microglia <- RunPCA(dsmerged_microglia, features = VariableFeatures(object =
       dsmerged_microglia))
print(dsmerged_microglia[["pca"]], dims = 1:5, nfeatures = 5)
VizDimLoadings(dsmerged_microglia, dims = 1:5, reduction = "pca")
ElbowPlot(dsmerged_microglia,ndims=50)
dsmerged_microglia <- FindNeighbors(dsmerged_microglia, dims = 1:42)
dsmerged_microglia <- FindClusters(dsmerged_microglia, resolution = 0.4)
dsmerged_microglia <- RunTSNE(dsmerged_microglia, dims = 1:42)
DimPlot(dsmerged_microglia, reduction = "tsne",label=T)

DimPlot(dsmerged_microglia, reduction = "tsne",label=F, group.by="batch")


# run harmony

dsmerged_microglia <- dsmerged_microglia %>%
RunHarmony("batch", plot_convergence = TRUE)

harmony_embeddings <- Embeddings(dsmerged_microglia, 'harmony')
harmony_embeddings[1:5, 1:5]

dsmerged_microglia <- dsmerged_microglia %>%
RunTSNE(reduction = "harmony", dims = 1:42) %>%
FindNeighbors(reduction = "harmony", dims = 1:42) %>%
FindClusters(resolution = 0.4) %>%
identity()

DimPlot(dsmerged_microglia, reduction = "tsne",label=F, group.by="batch")

FeaturePlot(dsmerged_microglia, features = c("nCount_RNA", "nFeature_RNA",
       "percent.mt")) # clusters 5 and 6 are low viability
FeaturePlot(dsmerged_microglia, features = c("Ly6c2", "P2ry12")) # cluster 8 has mixed
       signature: microglia/monocytes
FeaturePlot(dsmerged_microglia, features = c("Cldn5", "Itm2a", "Flt1", "Podxl")) #
       cluster 9 has mixed signature: microglia/endothelial cells

# remove low viability, mixed signature clusters

dsmerged_microglia <- subset(x = dsmerged_microglia, idents = c(5, 6, 8, 9),invert =
       TRUE)
DimPlot(dsmerged_microglia, reduction = "tsne",label=T)+ NoLegend()

rm(plot1, plot2, harmony_embeddings, all.genes, top10)


# subset and re-cluster PBS & GAS microglia only
Idents(object=dsmerged_microglia) <- dsmerged_microglia@meta.data$condition
dsmerged_microglia_PBS_GAS <- subset(x = dsmerged_microglia, idents = c("PBS",
       "GAS"),invert = FALSE)
```

```
DimPlot(dsmerged_microglia_PBS_GAS, reduction = "tsne",label=T)
dsmerged_microglia_PBS_GAS <- NormalizeData(dsmerged_microglia_PBS_GAS)
dsmerged_microglia_PBS_GAS <- FindVariableFeatures(dsmerged_microglia_PBS_GAS,
       selection.method = "vst", nfeatures = 3000)
top10 <- head(VariableFeatures(dsmerged_microglia_PBS_GAS), 10)
plot1 <- VariableFeaturePlot(dsmerged_microglia_PBS_GAS)
plot2 <- LabelPoints(plot = plot1, points = top10, repel = TRUE, xnudge = 0, ynudge =
       0)
plot2
all.genes <- rownames(dsmerged_microglia_PBS_GAS)
dsmerged_microglia_PBS_GAS <- ScaleData(dsmerged_microglia_PBS_GAS, features =
       all.genes)
dsmerged_microglia_PBS_GAS <- RunPCA(dsmerged_microglia_PBS_GAS, features =
       VariableFeatures(object = dsmerged_microglia_PBS_GAS))
print(dsmerged_microglia_PBS_GAS[["pca"]], dims = 1:5, nfeatures = 5)
VizDimLoadings(dsmerged_microglia_PBS_GAS, dims = 1:5, reduction = "pca")
ElbowPlot(dsmerged_microglia_PBS_GAS,ndims=50)
dsmerged_microglia_PBS_GAS <- FindNeighbors(dsmerged_microglia_PBS_GAS, dims = 1:32)
dsmerged_microglia_PBS_GAS <- FindClusters(dsmerged_microglia_PBS_GAS, resolution =
       0.4)
dsmerged_microglia_PBS_GAS <- RunTSNE(dsmerged_microglia_PBS_GAS, dims = 1:32)

Idents(object=dsmerged_microglia_PBS_GAS) <-
       dsmerged_microglia_PBS_GAS@meta.data$batch
DimPlot(dsmerged_microglia_PBS_GAS, reduction = "tsne",label=T)

# run harmony

dsmerged_microglia_PBS_GAS <- dsmerged_microglia_PBS_GAS %>%
RunHarmony("batch", plot_convergence = TRUE)

harmony_embeddings <- Embeddings(dsmerged_microglia_PBS_GAS, 'harmony')
harmony_embeddings[1:5, 1:5]

dsmerged_microglia_PBS_GAS <- dsmerged_microglia_PBS_GAS %>%
RunTSNE(reduction = "harmony", dims = 1:32) %>%
FindNeighbors(reduction = "harmony", dims = 1:32) %>%
FindClusters(resolution = 0.4) %>%
identity()

# confirming Ccl3 and Ccl4 expression is not due to ex vivo activation ---

exAM.markers <- c("Fos", "Jun", "Dusp1", "Hspa1a", "Hist1h1d", "Hist1h2ac", "Nfkbid",
       "Nfkbiz")
Idents(object=dsmerged_microglia_PBS_GAS) <-
       dsmerged_microglia_PBS_GAS@meta.data$condition
AverageExpression(dsmerged_microglia_PBS_GAS, features = exAM.markers)

dsmerged_microglia_PBS_GAS_new <- AddModuleScore(dsmerged_microglia_PBS_GAS, features
       = exAM.markers, name = "exAM_score")
FeaturePlot(dsmerged_microglia_PBS_GAS_new, features = "exAM_score1")
dsmerged_microglia_PBS_GAS_new$exAM_score <-
       dsmerged_microglia_PBS_GAS_new$exAM_score1

FeatureScatter(dsmerged_microglia_PBS_GAS_new, feature1 = "Ccl3", feature2 = "Ccl4",
       group.by = "inoculate")
```

```r
## code for scRNAseq analysis of perivascular macrophages ----

dsmerged_PBS_GAS <- readRDS("Directory/dsmerged_004_005_008_009_014_023_024.rds")

Idents(object=dsmerged_PBS_GAS) <- dsmerged_PBS_GAS@meta.data$cluster.names_all.cells
DimPlot(dsmerged_PBS_GAS, reduction = "tsne",label=T)
dsmerged_macrophages <- subset(x = dsmerged_PBS_GAS, idents = c("Macrophages"),invert
= FALSE)
DimPlot(dsmerged_macrophages, reduction = "tsne",label=T)

dsmerged_macrophages <- NormalizeData(dsmerged_macrophages)
dsmerged_macrophages <- FindVariableFeatures(dsmerged_macrophages, selection.method =
"vst", nfeatures = 3000)
top10 <- head(VariableFeatures(dsmerged_macrophages), 10)
plot1 <- VariableFeaturePlot(dsmerged_macrophages)
plot2 <- LabelPoints(plot = plot1, points = top10, repel = TRUE, xnudge = 0, ynudge =
0)
plot2
all.genes <- rownames(dsmerged_macrophages)
dsmerged_macrophages <- ScaleData(dsmerged_macrophages, features = all.genes)
dsmerged_macrophages <- RunPCA(dsmerged_macrophages, features =
VariableFeatures(object = dsmerged_macrophages))
print(dsmerged_macrophages[["pca"]], dims = 1:5, nfeatures = 5)
VizDimLoadings(dsmerged_macrophages, dims = 1:5, reduction = "pca")
ElbowPlot(dsmerged_macrophages,ndims=50)
dsmerged_macrophages <- FindNeighbors(dsmerged_macrophages, dims = 1:48)
dsmerged_macrophages <- FindClusters(dsmerged_macrophages, resolution = 1)
dsmerged_macrophages <- RunTSNE(dsmerged_macrophages, dims = 1:48)
DimPlot(dsmerged_macrophages, reduction = "tsne",label=T)
DimPlot(dsmerged_macrophages, reduction = "tsne",group.by="orig.ident")

immunemarkergenes <- c("Ptprc","Aif1","Tmem119", "Cd3e", "Cd4", "Cd8a", "Trdv4",
"Itgam", "Itgax","Cd19","Ms4a1","Ly6g")
moreimmunemarkers <- c("Ly6c2", "AceL", "Xcr1", "Cd209a", "Ccr9", "Nudt17", "Camp",
"Klrb1c", "Cd163l1", "Gata3")
PVMMarkers <- c("Cd163","Mrc1","Lyve1") # clusters 4 & 9 = BAMs

FeaturePlot(dsmerged_macrophages, features = PVMMarkers)

# remove mixed signature clusters

dsmerged_macrophages <- subset(x = dsmerged_macrophages, idents = c(10, 14, 15),invert
= TRUE)  # clusters 10 & 14 = microglia ; cluster 15 = T cells
DimPlot(dsmerged_macrophages, reduction = "tsne",label=T)

    # normalize & generate PCA
    dsmerged_macrophages <- NormalizeData(dsmerged_macrophages)
    dsmerged_macrophages <- FindVariableFeatures(dsmerged_macrophages,
selection.method = "vst", nfeatures = 3000)
    top10 <- head(VariableFeatures(dsmerged_macrophages), 10)
    plot1 <- VariableFeaturePlot(dsmerged_macrophages)
    plot2 <- LabelPoints(plot = plot1, points = top10, repel = TRUE, xnudge = 0,
ynudge = 0)
    plot2
    all.genes <- rownames(dsmerged_macrophages)
    dsmerged_macrophages <- ScaleData(dsmerged_macrophages, features = all.genes)
    dsmerged_macrophages <- RunPCA(dsmerged_macrophages, features =
VariableFeatures(object = dsmerged_macrophages))
    print(dsmerged_macrophages[["pca"]], dims = 1:5, nfeatures = 5)
    VizDimLoadings(dsmerged_macrophages, dims = 1:5, reduction = "pca")
    ElbowPlot(dsmerged_macrophages,ndims=50)
    dsmerged_macrophages <- FindNeighbors(dsmerged_macrophages, dims = 1:48)
    dsmerged_macrophages <- FindClusters(dsmerged_macrophages, resolution = 1)
    dsmerged_macrophages <- RunTSNE(dsmerged_macrophages, dims = 1:39)
    DimPlot(dsmerged_macrophages, reduction = "tsne",label=T)
    DimPlot(dsmerged_macrophages, reduction = "tsne",group.by="orig.ident")
```

```r
    rm(plot1, plot2, all.genes, top10)

    saveRDS(dsmerged_macrophages, file =
"Directory/dsmerged_macrophages_004_005_008_009_014_023_024.rds")


# perivascular macrophages ----

    PVMMarkers <- c("Cd163","Mrc1","Lyve1")
    BAMMarkers <- c("P2rx7","Mrc1","Folr2", "Nrp1", "Cd63", "Cd38", "Clec12a", "H2-
Ab")
    FeaturePlot(dsmerged_macrophages, features = PVMMarkers)


    # subset perivascular macrophages & run DE

        dsmerged_PVMs <- subset(x = dsmerged_macrophages, idents = c(5, 8, 10,
16),invert = FALSE)
        Idents(object=dsmerged_PVMs) <- dsmerged_PVMs@meta.data$inoculate
        DimPlot(dsmerged_PVMs, reduction = "tsne", label=T)
        dsmerged_PVMs <- FindMarkers(dsmerged_PVMs, ident.1 = "GAS", ident.2 = "PBS",
min.pct = 0, logfc.threshold = 0)
```

```
# code for scRNAseq analysis of NALT/OE after GAS infection ----

# import data files & merge
ds006.data <- Read10X(data.dir = "Directory/CW006/raw_feature_bc_matrix")
ds007.data <- Read10X(data.dir = "Directory/CW007/raw_feature_bc_matrix")
ds0019.data <- Read10X(data.dir = "Directory/CW019/raw_feature_bc_matrix")
ds0020.data <- Read10X(data.dir = "Directory/CW020/raw_feature_bc_matrix")

ds006 <- CreateSeuratObject(counts = ds006.data, project = "PBS", min.cells = 3,
       min.features = 200)
ds007 <- CreateSeuratObject(counts = ds007.data, project = "GAS", min.cells = 3,
       min.features = 200)
ds0019 <- CreateSeuratObject(counts = ds0019.data, project = "PBS", min.cells = 3,
       min.features = 200)
ds0020 <- CreateSeuratObject(counts = ds0020.data, project = "GAS", min.cells = 3,
       min.features = 200)

dsmerged_NALT <- merge(ds006, y = c(ds007, ds0019, ds0020), add.cell.ids = c("CW_006",
       "CW_007", "CW_019", "CW_020"))

rm(ds006, ds006.data, ds007, ds007.data, ds0019, ds0019.data, ds0020, ds0020.data)

#adding meta data

samplevec <- rep("CW006",nrow(dsmerged_NALT@meta.data))
samplevec[grep("CW_007",rownames(dsmerged_NALT@meta.data))] <- "CW007"
samplevec[grep("CW_019",rownames(dsmerged_NALT@meta.data))] <- "CW019"
samplevec[grep("CW_020",rownames(dsmerged_NALT@meta.data))] <- "CW020"
table(samplevec)
dsmerged_NALT@meta.data$sample=samplevec

# add batch to meta data
batchvec <- rep("batch B",nrow(dsmerged_NALT@meta.data))
batchvec[grep("CW_019",rownames(dsmerged_NALT@meta.data))] <- "batch L"
batchvec[grep("CW_020",rownames(dsmerged_NALT@meta.data))] <- "batch L"
table(batchvec)
dsmerged_NALT@meta.data$batch=batchvec

# add enrichment to meta data
enrichmentvec <- rep("live_FACS",nrow(dsmerged_NALT@meta.data))
enrichmentvec[grep("CW_019",rownames(dsmerged_NALT@meta.data))] <- "cd11b_FACS"
enrichmentvec[grep("CW_020",rownames(dsmerged_NALT@meta.data))] <- "cd11b_FACS"
table(enrichmentvec)
dsmerged_NALT@meta.data$enrichment=enrichmentvec

# add inoculate to meta data
inoculatevec <- rep("GAS",nrow(dsmerged_NALT@meta.data))
inoculatevec[grep("CW_006",rownames(dsmerged_NALT@meta.data))] <- "PBS"
inoculatevec[grep("CW_019",rownames(dsmerged_NALT@meta.data))] <- "PBS"
table(inoculatevec)
dsmerged_NALT@meta.data$inoculate=inoculatevec

# add tissue to meta data
tissuevec <- rep("NALT",nrow(dsmerged_NALT@meta.data))
table(tissuevec)
dsmerged_NALT@meta.data$tissue=tissuevec

# add genotype to meta data
genotypevec <- rep("WT",nrow(dsmerged_NALT@meta.data))
table(genotypevec)
dsmerged_NALT@meta.data$genotype=genotypevec

# add drug to meta data
drugvec <- rep("none",nrow(dsmerged_NALT@meta.data))
table(drugvec)
dsmerged_NALT@meta.data$drug=drugvec
```

```
# add condition to meta data
conditionvec <- rep("GAS",nrow(dsmerged_NALT@meta.data))
conditionvec[grep("CW_006",rownames(dsmerged_NALT@meta.data))] <- "PBS"
conditionvec[grep("CW_019",rownames(dsmerged_NALT@meta.data))] <- "PBS"
table(conditionvec)
dsmerged_NALT@meta.data$condition=conditionvec

rm(samplevec, batchvec, enrichmentvec, tissuevec, genotypevec, drugvec, inoculatevec,
       conditionvec)

# remove dead cells

dsmerged_NALT[["percent.mt"]] <- PercentageFeatureSet(dsmerged_NALT, pattern = "^mt-")
VlnPlot(dsmerged_NALT, features = c("nFeature_RNA", "nCount_RNA", "percent.mt"), ncol
       = 3)
dsmerged_NALT <- subset(dsmerged_NALT, subset = nCount_RNA > 1000 & nCount_RNA < 50000
       & percent.mt < 20)
VlnPlot(dsmerged_NALT, features = c("nFeature_RNA", "nCount_RNA", "percent.mt"), ncol
       = 3)

# normalize & generate PCA
dsmerged_NALT <- NormalizeData(dsmerged_NALT)
dsmerged_NALT <- FindVariableFeatures(dsmerged_NALT, selection.method = "vst",
       nfeatures = 3000)
top10 <- head(VariableFeatures(dsmerged_NALT), 10)
plot1 <- VariableFeaturePlot(dsmerged_NALT)
plot2 <- LabelPoints(plot = plot1, points = top10, repel = TRUE, xnudge = 0, ynudge =
       0)
plot2
all.genes <- rownames(dsmerged_NALT)
dsmerged_NALT <- ScaleData(dsmerged_NALT, features = all.genes)
dsmerged_NALT <- RunPCA(dsmerged_NALT, features = VariableFeatures(object =
       dsmerged_NALT))
print(dsmerged_NALT[["pca"]], dims = 1:5, nfeatures = 5)
VizDimLoadings(dsmerged_NALT, dims = 1:5, reduction = "pca")
ElbowPlot(dsmerged_NALT,ndims=50)
dsmerged_NALT <- FindNeighbors(dsmerged_NALT, dims = 1:48)
dsmerged_NALT <- FindClusters(dsmerged_NALT, resolution = 2)
dsmerged_NALT <- RunTSNE(dsmerged_NALT, dims = 1:48)
DimPlot(dsmerged_NALT, reduction = "tsne",label=T)
DimPlot(dsmerged_NALT, reduction = "tsne",group.by="inoculate")
rm(plot1, plot2, all.genes, top10)


# assign cell types

neuroglial.markers <- c("Snap25", "Dcx", "Omp", "Adcy3", "Gfap","Aqp4",
       "Frzb","Cldn5", "Pecam1", "Pdgfra", "Pdgfrb", "Atp13a5", "Col1a1", "Fbln1")
lymphoid.markers <- c("Ptprc", "Cd3e", "Cd4", "Cd8a", "Cd163l1", "Klrb1c", "Cd19",
       "Ms4a1")
myeloid.markers <- c("Ptprc", "Tmem119", "Aif1", "Itgam", "Cd68", "Ly6g", "Camp",
       "Ly6c2", "Itgax", "Xcr1", "Cd209a", "Ccr9", "Nudt17")
macrophage.vs.microglia.markers <- c("Ptprc", "Tmem119", "Cx3cr1", "P2ry12", "Itgam",
       "Aif1", "Hexb", "Plac8", "Mrc1", "Cd163", "Lyve1")
fibroblast.markers <- c("Col1a1", "Col1a2", "Col5a1", "Loxl1", "Lum", "Fbln1",
       "Fbln2", "Cd34", "Pdgfra")
schwann.cell.markers <-
       c("Sox10","Gap43","Fabp7","Mpz","Dhh","Ngfr","S100a1","S100b","Egr2","Mbp","Nca
       m1")
epithelial.markers <- c("Epcam", "Cd24a", "Itga6", "Ceacam1", "St6gal1", "Itgb4",
       "Il1r1", "Prom1", "Ddr2", "Krt19") # Itga6 = CD49f ; Ceacam1 = CD66a ; St6gal1
       = CD75 ; Itgb4 = CD104 ; Il1r1 = CD121a ; Prom1 = CD133 ; Ddr2 = CD167 ; Krt19
       = Cytokeratin
non.CNS.EC.markers  <- c("Pecam1", "Vwf", "Erg", "Cdh5", "Cav1", "Cldn1")
```

```
granulocyte.markers <- c("Itga2", "Fcer1a", "Cd200r3", "Itgam", "Ccr3", "Siglecf",
       "Ly6c2", "Cd66a", "Cd66b", "Cd66c", "Cd66d")

rm(neuroglial.markers, lymphoid.markers, myeloid.markers,
       macrophage.vs.microglia.markers, fibroblast.markers, schwann.cell.markers,
       epithelial.markers, non.CNS.EC.markers , granulocyte.markers)

# rename clusters

Idents(object=dsmerged_NALT) <- dsmerged_NALT@meta.data$RNA_snn_res.1
DimPlot(dsmerged_NALT, reduction = "tsne",label=T)
new.cluster.ids <- c("Neurons", "Neurons", "Neurons", "Neutrophils", "Neurons",
       "Neurons", "Monocytes", "Neurons", "Neurons", "Neurons", "Neutrophils",
       "Neutrophils", "Macrophages", "Neutrophils", "Dendritic cells", "Neurons",
       "Neutrophils", "CD4 T cells", "Monocytes", "Macrophages", "Neurons", "B cells",
       "Proliferating cells", "Neurons", "CD8 T cells", "Mixed signatures:
       Neurons/Epithelial", "B cells", "Mixed signatures: Neutrophils/Macrophages",
       "Epithelial cells", "Epithelial cells", "Erythropoietic cells", "Mixed
       signatures: Neutrophils/Endothelial cells", "Dendritic cells", "Dendritic
       cells", "Epithelial cells", "gamma delta T cells", "NK cells", "B cells",
       "Neurons", "Neurons", "Neutrophils", "Monocytes", "Epithelial cells",
       "Olfactory ensheathing cells", "Fibroblasts", "Mixed signatures: Dendritic
       cells/Endothelial cells", "Mixed signatures: Epithelial cells/Astrocytes",
       "Epithelial cells", "Epithelial cells", "Neurons", "Neurons", "Mixed
       signatures: Neurons/Macrophages", "Epithelial cells", "Mixed signatures:
       Granulocytes/Endothelial cells")
names(new.cluster.ids) <- levels(dsmerged_NALT)
dsmerged_NALT <- RenameIdents(dsmerged_NALT, new.cluster.ids)
DimPlot(dsmerged_NALT, reduction = "tsne",label=T)+ NoLegend()
dsmerged_NALT[["cluster.names_NALT"]] <- Idents(object = dsmerged_NALT) # stash
       identity
rm(new.cluster.ids)

# remove mixed signature clusters

dsmerged_NALT <- subset(x = dsmerged_NALT, idents = c("Erythropoietic cells",
       "Proliferating cells", "Mixed signatures: Neurons/Epithelial", "Mixed
       signatures: Neutrophils/Macrophages", "Mixed signatures:
       Neutrophils/Endothelial cells", "Mixed signatures: Dendritic cells/Endothelial
       cells", "Mixed signatures: Epithelial cells/Astrocytes", "Mixed signatures:
       Neurons/Macrophages", "Mixed signatures: Granulocytes/Endothelial cells"),
       invert = TRUE)
```

### Document S3: Scripts for processing of MERFISH data

```
## code for initial MERFISH analysis of olfactory bulb after GAS infection (UMass) ---
       -

# analysis run with R v 4.0.0 and Seurat 4.1.0.9005

# load programs
library(Seurat)
library(dplyr)
library(Matrix)
library(reticulate)
library(scCustomize)
library(tidyverse)
library(patchwork)
library(viridis)
library(qs)
library(sva)
library(sp)

options(scipen = 100)

#read in data. transpose so rows are genes and columns are cells. remove "blank"
       genes. remove volume < 50. add metadata for vol, X, y. normalize to volume
OB1 <- read.table(file = '**/sample1_merged_metadata_and_partitions.csv',sep=",",
       header = TRUE, row.names=1)
OB1 <- t(OB1)
Big.Cells <- which(OB1[464,]>= 50)
OB1 <- OB1[,Big.Cells]
volume<-OB1[464,]
center.x<-OB1[465,]
center.y<-OB1[466,]
OB1 <- OB1[1:391,]
OB1 <- OB1/volume
OB1_s <- CreateSeuratObject(counts = OB1)
Volumes <- data.frame(volume)
Center.x <- data.frame(center.x)
Center.y <- data.frame(center.y)
OB1_s@meta.data <- cbind(OB1_s@meta.data,Volumes)
OB1_s@meta.data <- cbind(OB1_s@meta.data,Center.x)
OB1_s@meta.data <- cbind(OB1_s@meta.data,Center.y)

#split object into 2 objects containing cells from PBS vs. GAS sections on the same
       coverslip
OB1.combined <- OB1_s
PBS1 <- subset(OB1.combined,subset = center.y < 7000)
PBS1 <- subset(PBS1,subset = center.y < 3540 | center.x < 2510)
GAS1 <- subset(OB1.combined,subset = center.y > 7000)
PBS1$group <- "PBS"
PBS1$sample <- "PBS_1"
GAS1$group <- "GAS"
GAS1$sample <- "GAS_1"

#repeat for samples 2-4
OB2 <- read.table(file = '**/sample2_merged_metadata_and_partitions.csv',sep=",",
       header = TRUE, row.names=1)
OB2 <- t(OB2)
Big.Cells <- which(OB2[464,]>= 50)
```

```
OB2 <- OB2[,Big.Cells]
volume<-OB2[464,]
center.x<-OB2[465,]
center.y<-OB2[466,]
OB2 <- OB2[1:391,]
OB2 <- OB2/volume
OB2_s <- CreateSeuratObject(counts = OB2)
Volumes <- data.frame(volume)
Center.x <- data.frame(center.x)
Center.y <- data.frame(center.y)
OB2_s@meta.data <- cbind(OB2_s@meta.data,Volumes)
OB2_s@meta.data <- cbind(OB2_s@meta.data,Center.x)
OB2_s@meta.data <- cbind(OB2_s@meta.data,Center.y)

OB2.combined <- OB2_s
PBS2 <- subset(OB2.combined,subset = center.y < 5600)
GAS2 <- subset(OB2.combined,subset = center.y > 7000)
GAS2 <- subset(GAS2,subset = center.y < 12284 | center.x < 2812)
PBS2$group <- "PBS"
PBS2$sample <- "PBS_2"
GAS2$group <- "GAS"
GAS2$sample <- "GAS_2"

OB3 <- read.table(file = '**/sample3_merged_metadata_and_partitions.csv',sep=",",
       header = TRUE, row.names=1)
OB3 <- t(OB3)
Big.Cells <- which(OB3[464,]>= 50)
OB3 <- OB3[,Big.Cells]
volume<-OB3[464,]
center.x<-OB3[465,]
center.y<-OB3[466,]
OB3 <- OB3[1:391,]
OB3 <- OB3/volume
OB3_s <- CreateSeuratObject(counts = OB3)
Volumes <- data.frame(volume)
Center.x <- data.frame(center.x)
Center.y <- data.frame(center.y)
OB3_s@meta.data <- cbind(OB3_s@meta.data,Volumes)
OB3_s@meta.data <- cbind(OB3_s@meta.data,Center.x)
OB3_s@meta.data <- cbind(OB3_s@meta.data,Center.y)

OB3.combined <- OB3_s
PBS3 <- subset(OB3.combined,subset = center.y > 7000)
GAS3 <- subset(OB3.combined,subset = center.y < 7000)
PBS3$group <- "PBS"
PBS3$sample <- "PBS_3"
GAS3$group <- "GAS"
GAS3$sample <- "GAS_3"

OB4 <- read.table(file = '**/sample4_merged_metadata_and_partitions.csv',sep=",",
       header = TRUE, row.names=1)
OB4 <- t(OB4)
Big.Cells <- which(OB4[464,]>= 50)
OB4 <- OB4[,Big.Cells]
volume<-OB4[464,]
center.x<-OB4[465,]
center.y<-OB4[466,]
```

```
OB4 <- OB4[1:391,]
OB4 <- OB4/volume
OB4_s <- CreateSeuratObject(counts = OB4)
Volumes <- data.frame(volume)
Center.x <- data.frame(center.x)
Center.y <- data.frame(center.y)
OB4_s@meta.data <- cbind(OB4_s@meta.data,Volumes)
OB4_s@meta.data <- cbind(OB4_s@meta.data,Center.x)
OB4_s@meta.data <- cbind(OB4_s@meta.data,Center.y)

OB4.combined <- OB4_s
PBS4 <- subset(OB4.combined,subset = center.x < 4500)
GAS4<- subset(OB4.combined,subset = center.x >4500)
PBS4$group <- "PBS"
PBS4$sample <- "PBS_4"
GAS4$group <- "GAS"
GAS4$sample <- "GAS_4"

##remove portions of GAS4 section that are not in the olfactory bulb
GAS4.x <- GAS4@meta.data[,5]
head(GAS4.x)
GAS4.y <- GAS4@meta.data[,6]
head(GAS4.y)

#X Y points defining the non-olfactory bulb regions in GAS4
GAS4_Rep1.x <-
       c(6003.06,6034.19,6091.03,6176.58,6245.24,6187.08,6077.35,5984.03,5983.12,6647.
       6,6835.52,6912.2,7359.27,7376.72,7473.33,6068.5,3939.8,3939.8,5915.31)
GAS4_Rep1.y <-
       c(3294.86,3526.63,3711.69,3976.82,4048.19,4501.78,4711.69,4914.02,4961.04,5080.
       32,5356.65,5389.6,5283.13,5912.29,6348.2,7000,7000,2631.15,3181)
GAS4_Rep2.x <-
       c(6503.67,6719.63,6778.14,6813.86,6763.36,6590.23,6495.63,6391.15,6397.21,6434.
       48)
GAS4_Rep2.y <-
       c(3044.73,3134.84,3190.46,3381.29,3537.53,3529.87,3444.28,3200.13,3103.89,3062.
       16)

##filter out cells from GAS contained within the non-OB polygon
GAS4_Rep1 <- point.in.polygon(GAS4.x, GAS4.y, GAS4_Rep1.x, GAS4_Rep1.y)
GAS4_Rep2 <- point.in.polygon(GAS4.x, GAS4.y, GAS4_Rep2.x, GAS4_Rep2.y)

GAS4.info <- cbind.data.frame(GAS4_Rep1,GAS4_Rep2)
head(GAS4.info, 50)
Region <- rowSums(GAS4.info)
GAS4.info2 <- GAS4.info %>% mutate(Region2 = if_else(Region > 0,"non_OB","OB"))
head(GAS4.info2, 50)
Region_all <-GAS4.info2[,3]

GAS4 <- AddMetaData(GAS4, metadata = Region_all, col.name = "Region")
GAS4 <- subset(GAS4,Region=="OB")

## merge objects for all samples
OB.combined <- merge(PBS1, y=c(GAS1, PBS2, GAS2, PBS3, GAS3,PBS4,GAS4))

# remove cells with nFeature_RNA < 11)
OB.combined <- subset(OB.combined,nFeature_RNA > 10)
```

```
#normalize data and find variable features (all transcripts are used as variable
       features)
OB.combined <-NormalizeData(OB.combined)
OB.combined <- FindVariableFeatures(OB.combined, selection.method = "vst", nfeatures =
       391)

#scale data and run PCA
all.genes <- rownames(OB.combined)
OB.combined <- ScaleData(OB.combined, features = all.genes)
OB.combined <- RunPCA(OB.combined, features = VariableFeatures(object =
       OB.combined),npcs = 50)

## define each coverslip as a batch for ComBat batch correction
batchname = OB.combined@meta.data$sample
batchid = rep(1,length(batchname))
batchid[batchname=="PBS_1"] = 1
batchid[batchname=="PBS_2"] = 2
batchid[batchname=="PBS_3"] = 3
batchid[batchname=="PBS_4"] = 4
batchid[batchname=="GAS_1"] = 1
batchid[batchname=="GAS_2"] = 2
batchid[batchname=="GAS_3"] = 3
batchid[batchname=="GAS_4"] = 4
names(batchid) = rownames(OB.combined@meta.data)
OB.combined <- AddMetaData(object = OB.combined, metadata = batchid, col.name =
       "batchid")
table(OB.combined@meta.data$batchid)

## perform ComBat batch correction
m = as.data.frame(as.matrix(OB.combined@assays$RNA@data))
m = m[rowSums(m)>0,]
com = ComBat(m, batchid, prior.plots=FALSE, par.prior=TRUE)

##replace data slot with ComBat batch corrected data
OB.combined@assays$RNA@data = Matrix(as.matrix(com),sparse=T)

##rescale data and rerun PCA
OB.combined = ScaleData(OB.combined)
OB.combined <- RunPCA(OB.combined, features = VariableFeatures(object =
       OB.combined),npcs = 50)

#test of number of significant Principal Components. 28 are used for further analysis.
OB.combined <- JackStraw(OB.combined, num.replicate = 100,dims=50)
OB.combined <- ScoreJackStraw(OB.combined, dims = 1:50)
JackStrawPlot(OB.combined, dims = 1:50)

##Clustering and linear dimensionality reduction
OB.combined <- FindNeighbors(OB.combined, dims = 1:28)
OB.combined <- FindClusters(OB.combined, resolution = 2.4)
OB.combined <- RunTSNE(OB.combined, dims = 1:28)

## Cell-type annotation of clusters
Idents(OB.combined) <- OB.combined@meta.data$seurat_clusters
new.cluster.ids <- c("Granule Cells","Granule Cells","Granule Cells","Granule
       Cells","Periglomerular Cells",
```

```
                  "Granule Cells","Granule Cells","Astrocytes","Periglomerular
        Cells","Granule Cells",
                  "Immature Periglomerular Cells","Granule Cells","Olfactory
        Ensheathing Cells","Astrocytes","Olfactory Ensheathing Cells",
                  "Endothelial Cells","External Tufted Cells","Mitral/Tufted
        Cells","Periglomerular Cells","Mitral/Tufted Cells",
                  "Meningeal Fibroblasts","Olfactory Ensheathing
        Cells","Neuroblasts","Somatostatin-immunoreactive Neurons","Astrocyte/Neuron
        Hybrids",
                  "Endothelial Cells","Endothelial/Neurons/Astrocyte
        Hybrids","Astrocytes/Granule Cell Hybrids","Pericytes","Oligodendrocytes",
                  "OPCs","Microglia","Microglia","External Tufted
        Cells","Microglia",
                  "Border-associated Macrophages","OEC/Neuron
        Hybrids","Endothelial/Pericyte Hybrids","Olfactory Ensheathing
        Cells","Macrophage/Granule Cell Hybrids",
                  "Periglomerular Cells","T
        Cells","Neutrophils","Astrocyte/Mitral/Tufted Hybrids","Artifacts")
names(new.cluster.ids) <- levels(OB.combined)
OB.combined <- RenameIdents(OB.combined, new.cluster.ids)
OB.combined$celltype <- Idents(OB.combined)

##data output
saveRDS(OB.combined,"220428_AgalliuLab_GAS_combat_AllCells_norm.rds")
```

```r
## code for MERFISH microglia analysis of T cell nearest neighbor distance (NND)
       (UMass) ----

# analysis run with R v 4.0.0 and Seurat 4.1.0.9005

## removal of cell clusters annotated as hybrids and reanalysis.
OB.NoHybrids <- subset(OB.combined, idents= c("Microglia","Granule
       Cells","Periglomerular Cells","Astrocytes","Olfactory Ensheathing
       Cells","Mitral/Tufted Cells","Neuroblasts","Immature Periglomerular
       Cells","Meningeal Fibroblasts","Oligodendrocytes","OPCs","Border-associated
       Macrophages","Endothelial Cells","Pericytes","Somatostatin-immunoreactive
       Neurons","T Cells","Neutrophils"))
OB.NoHybrids <- FindVariableFeatures(OB.NoHybrids, selection.method = "vst", nfeatures
       = 391)
all.genes <- rownames(OB.NoHybrids)
OB.NoHybrids <- ScaleData(OB.NoHybrids, features = all.genes)
OB.NoHybrids <- RunPCA(OB.NoHybrids, features = VariableFeatures(object =
       OB.NoHybrids),npcs = 50)
OB.NoHybrids <- FindNeighbors(OB.NoHybrids, dims = 1:28)
OB.NoHybrids <- FindClusters(OB.NoHybrids, resolution = 2.4)
OB.NoHybrids <- RunUMAP(OB.NoHybrids, dims = 1:28)
OB.NoHybrids <- RunTSNE(OB.NoHybrids, dims = 1:28)
DimPlot(OB.NoHybrids,reduction="tsne")

##output data file
saveRDS(OB.NoHybrids,"220428_AgalliuLab_GAS_combat_AllCells_NoHybrids.rds")

#### Microglia - T cell Nearest Neighbor Distance (NND) ###############

OB.NoHybrids <- readRDS("220428_AgalliuLab_GAS_combat_AllCells_NoHybrids.rds")

##for each sample, subset object to microglia and T Cells
PBS1.NND <- subset(OB.NoHybrids, subset = sample == "PBS_1" & celltype == "Microglia"
       | sample == "PBS_1" & celltype == "T Cells")
PBS1.Microglia <- subset(OB.NoHybrids, subset = sample == "PBS_1" & celltype ==
       "Microglia")

## generate .csv file of cell X Y position used for nearest neighbor python script
output_dataframe=data.frame(PBS1.NND$celltype)
output_dataframe=cbind(output_dataframe,PBS1.NND@meta.data$center.x)
output_dataframe=cbind(output_dataframe,PBS1.NND@meta.data$center.y)
output_dataframe <- cbind(rownames(output_dataframe), data.frame(output_dataframe,
       row.names=NULL))
colnames(output_dataframe) <- c('cell', 'cluster', 'x', 'y')
write.csv(output_dataframe, 'all_cells.csv', row.names=FALSE)

#run NND analysis using "python find_nearest_neighbors2.py"

### read output file of "python find_nearest_neighbors2.py" back into Rstudio and
       attach as metadata to microglia.
NND.Microglia.TCells.PBS1 <-read.csv("python_nearest_neighbor_outputs/PBS1 Microglia_T
       Cells_nearest_neighbors.csv",header=FALSE)
NNDs.MG.TCells <- NND.Microglia.TCells.PBS1[,3]
NND.MG.TCells <- data.frame(NNDs.MG.TCells)
PBS1.Microglia@meta.data <- cbind(PBS1.Microglia@meta.data,NND.MG.TCells)
```

```
##repeat for sample 2-8
PBS2.NND <- subset(OB.NoHybrids, subset = sample == "PBS_2" & celltype == "Microglia"
       | sample == "PBS_2" & celltype == "T Cells")
PBS2.Microglia <- subset(OB.NoHybrids, subset = sample == "PBS_2" & celltype ==
       "Microglia")

output_dataframe=data.frame(PBS2.NND$celltype)
output_dataframe=cbind(output_dataframe,PBS2.NND@meta.data$center.x)
output_dataframe=cbind(output_dataframe,PBS2.NND@meta.data$center.y)
output_dataframe <- cbind(rownames(output_dataframe), data.frame(output_dataframe,
       row.names=NULL))
colnames(output_dataframe) <- c('cell', 'cluster', 'x', 'y')
write.csv(output_dataframe, 'all_cells.csv', row.names=FALSE)

NND.Microglia.TCells.PBS2 <-read.csv("python_nearest_neighbor_outputs/PBS2 Microglia_T
       Cells_nearest_neighbors.csv",header=FALSE)
NNDs.MG.TCells <- NND.Microglia.TCells.PBS2[,3]
NND.MG.TCells <- data.frame(NNDs.MG.TCells)
PBS2.Microglia@meta.data <- cbind(PBS2.Microglia@meta.data,NND.MG.TCells)

PBS3.NND <- subset(OB.NoHybrids, subset = sample == "PBS_3" & celltype == "Microglia"
       | sample == "PBS_3" & celltype == "T Cells")
PBS3.Microglia <- subset(OB.NoHybrids, subset = sample == "PBS_3" & celltype ==
       "Microglia")

output_dataframe=data.frame(PBS3.NND$celltype)
output_dataframe=cbind(output_dataframe,PBS3.NND@meta.data$center.x)
output_dataframe=cbind(output_dataframe,PBS3.NND@meta.data$center.y)
output_dataframe <- cbind(rownames(output_dataframe), data.frame(output_dataframe,
       row.names=NULL))
colnames(output_dataframe) <- c('cell', 'cluster', 'x', 'y')
write.csv(output_dataframe, 'all_cells.csv', row.names=FALSE)

NND.Microglia.TCells.PBS3 <-read.csv("python_nearest_neighbor_outputs/PBS3 Microglia_T
       Cells_nearest_neighbors.csv",header=FALSE)
NNDs.MG.TCells <- NND.Microglia.TCells.PBS3[,3]
NND.MG.TCells <- data.frame(NNDs.MG.TCells)
PBS3.Microglia@meta.data <- cbind(PBS3.Microglia@meta.data,NND.MG.TCells)

PBS4.NND <- subset(OB.NoHybrids, subset = sample == "PBS_4" & celltype == "Microglia"
       | sample == "PBS_4" & celltype == "T Cells")
PBS4.Microglia <- subset(OB.NoHybrids, subset = sample == "PBS_4" & celltype ==
       "Microglia")
output_dataframe=data.frame(PBS4.NND$celltype)
output_dataframe=cbind(output_dataframe,PBS4.NND@meta.data$center.x)
output_dataframe=cbind(output_dataframe,PBS4.NND@meta.data$center.y)
output_dataframe <- cbind(rownames(output_dataframe), data.frame(output_dataframe,
       row.names=NULL))
colnames(output_dataframe) <- c('cell', 'cluster', 'x', 'y')
write.csv(output_dataframe, 'all_cells.csv', row.names=FALSE)

NND.Microglia.TCells.PBS4 <-read.csv("python_nearest_neighbor_outputs/PBS4 Microglia_T
       Cells_nearest_neighbors.csv",header=FALSE)
NNDs.MG.TCells <- NND.Microglia.TCells.PBS4[,3]
NND.MG.TCells <- data.frame(NNDs.MG.TCells)
PBS4.Microglia@meta.data <- cbind(PBS4.Microglia@meta.data,NND.MG.TCells)
```

```
GAS1.NND <- subset(OB.NoHybrids, subset = sample == "GAS_1" & celltype == "Microglia"
       | sample == "GAS_1" & celltype == "T Cells")
GAS1.Microglia <- subset(OB.NoHybrids, subset = sample == "GAS_1" & celltype ==
       "Microglia")

output_dataframe=data.frame(GAS1.NND$celltype)
output_dataframe=cbind(output_dataframe,GAS1.NND@meta.data$center.x)
output_dataframe=cbind(output_dataframe,GAS1.NND@meta.data$center.y)
output_dataframe <- cbind(rownames(output_dataframe), data.frame(output_dataframe,
       row.names=NULL))
colnames(output_dataframe) <- c('cell', 'cluster', 'x', 'y')
write.csv(output_dataframe, 'all_cells.csv', row.names=FALSE)

NND.Microglia.TCells.GAS1 <-read.csv("python_nearest_neighbor_outputs/GAS1 Microglia_T
       Cells_nearest_neighbors.csv",header=FALSE)
NNDs.MG.TCells <- NND.Microglia.TCells.GAS1[,3]
NND.MG.TCells <- data.frame(NNDs.MG.TCells)
GAS1.Microglia@meta.data <- cbind(GAS1.Microglia@meta.data,NND.MG.TCells)

GAS2.NND <- subset(OB.NoHybrids, subset = sample == "GAS_2" & celltype == "Microglia"
       | sample == "GAS_2" & celltype == "T Cells")
GAS2.Microglia <- subset(OB.NoHybrids, subset = sample == "GAS_2" & celltype ==
       "Microglia")

output_dataframe=data.frame(GAS2.NND$celltype)
output_dataframe=cbind(output_dataframe,GAS2.NND@meta.data$center.x)
output_dataframe=cbind(output_dataframe,GAS2.NND@meta.data$center.y)
output_dataframe <- cbind(rownames(output_dataframe), data.frame(output_dataframe,
       row.names=NULL))
colnames(output_dataframe) <- c('cell', 'cluster', 'x', 'y')
write.csv(output_dataframe, 'all_cells.csv', row.names=FALSE)

NND.Microglia.TCells.GAS2 <-read.csv("python_nearest_neighbor_outputs/GAS2 Microglia_T
       Cells_nearest_neighbors.csv",header=FALSE)
NNDs.MG.TCells <- NND.Microglia.TCells.GAS2[,3]
NND.MG.TCells <- data.frame(NNDs.MG.TCells)
GAS2.Microglia@meta.data <- cbind(GAS2.Microglia@meta.data,NND.MG.TCells)

GAS3.NND <- subset(OB.NoHybrids, subset = sample == "GAS_3" & celltype == "Microglia"
       | sample == "GAS_3" & celltype == "T Cells")
GAS3.Microglia <- subset(OB.NoHybrids, subset = sample == "GAS_3" & celltype ==
       "Microglia")

output_dataframe=data.frame(GAS3.NND$celltype)
output_dataframe=cbind(output_dataframe,GAS3.NND@meta.data$center.x)
output_dataframe=cbind(output_dataframe,GAS3.NND@meta.data$center.y)
output_dataframe <- cbind(rownames(output_dataframe), data.frame(output_dataframe,
       row.names=NULL))
colnames(output_dataframe) <- c('cell', 'cluster', 'x', 'y')
write.csv(output_dataframe, 'all_cells.csv', row.names=FALSE)

NND.Microglia.TCells.GAS3 <-read.csv("python_nearest_neighbor_outputs/GAS3 Microglia_T
       Cells_nearest_neighbors.csv",header=FALSE)
NNDs.MG.TCells <- NND.Microglia.TCells.GAS3[,3]
NND.MG.TCells <- data.frame(NNDs.MG.TCells)
GAS3.Microglia@meta.data <- cbind(GAS3.Microglia@meta.data,NND.MG.TCells)
```

```
GAS4.NND <- subset(OB.NoHybrids, subset = sample == "GAS_4" & celltype == "Microglia"
       | sample == "GAS_4" & celltype == "T Cells")
GAS4.Microglia <- subset(OB.NoHybrids, subset = sample == "GAS_4" & celltype ==
       "Microglia")
output_dataframe=data.frame(GAS4.NND$celltype)
output_dataframe=cbind(output_dataframe,GAS4.NND@meta.data$center.x)
output_dataframe=cbind(output_dataframe,GAS4.NND@meta.data$center.y)
output_dataframe <- cbind(rownames(output_dataframe), data.frame(output_dataframe,
       row.names=NULL))
colnames(output_dataframe) <- c('cell', 'cluster', 'x', 'y')
write.csv(output_dataframe, 'all_cells.csv', row.names=FALSE)

NND.Microglia.TCells.GAS4 <-read.csv("python_nearest_neighbor_outputs/GAS4 Microglia_T
       Cells_nearest_neighbors.csv",header=FALSE)
NNDs.MG.TCells <- NND.Microglia.TCells.GAS4[,3]
NND.MG.TCells <- data.frame(NNDs.MG.TCells)
GAS4.Microglia@meta.data <- cbind(GAS4.Microglia@meta.data,NND.MG.TCells)

##merge microglia objects containing NND metadata for all 8 samples
Microglia <-merge(PBS1.Microglia,y=
       c(PBS2.Microglia,PBS3.Microglia,PBS4.Microglia,GAS1.Microglia,GAS2.Microglia,GA
       S3.Microglia,GAS4.Microglia))

##output data file
saveRDS(Microglia,"220921_AgalliuLab_GAS_combat_MicrogliaOnly_TCellNNDs.rds")
```

```r
## code for additional MERFISH analysis of olfactory bulb after GAS infection
       (Columbia) ----

# analysis run with R v 4.0.2 and Seurat 4.0.2

# load programs
library(Seurat)
library(dplyr)
library(ggplot2)

# open file
MERFISH_all_includes_mixed_signature <- readRDS("Directory/MERFISH/Data/RDS
       files/220428_AgalliuLab_GAS_combat_AllCells_norm.rds")# RDS file received from
       UMass; already batch-normalized through ComBat
DimPlot(MERFISH_all_includes_mixed_signature, reduction = "tsne", label = T)

# reformat batch ID in metadata
Idents(object=MERFISH_all_includes_mixed_signature) <-
       MERFISH_all_includes_mixed_signature@meta.data$sample
DimPlot(MERFISH_all_includes_mixed_signature, reduction = "tsne",label=T)
new.cluster.ids <- c("batch 1", "batch 1", "batch 2", "batch 2", "batch 3", "batch 3",
       "batch 4", "batch 4")
names(new.cluster.ids) <- levels(MERFISH_all_includes_mixed_signature)
MERFISH_all_includes_mixed_signature <-
       RenameIdents(MERFISH_all_includes_mixed_signature, new.cluster.ids)
DimPlot(MERFISH_all_includes_mixed_signature, reduction = "tsne",label=T)+ NoLegend()
MERFISH_all_includes_mixed_signature[["batch"]] <- Idents(object =
       MERFISH_all_includes_mixed_signature) # stash identity
MERFISH_all_includes_mixed_signature$batchid <- NULL # remove batch meta data from
       UMass

# modify cell type in metadata
Idents(object=MERFISH_all_includes_mixed_signature) <-
       MERFISH_all_includes_mixed_signature@meta.data$celltype
DimPlot(MERFISH_all_includes_mixed_signature, reduction = "tsne",label=T)
new.cluster.ids <- c("Neurons", "Neurons", "Astrocytes", "Neurons", "Olfactory
       ensheathing cells", "Endothelial cells", "Neurons", "Neurons", "Fibroblasts",
       "Neurons", "Neurons", "Mixed signature: Astrocytes/Neurons", "Mixed signature:
       Astrocytes/Neurons/Endothelial cells", "Mixed signature: Astrocytes/Neurons",
       "Pericytes", "Oligodendrocytes", "OPCs", "Microglia", "Macrophages", "Mixed
       signature: Olfactory ensheathing cells/Neurons", "Mixed signature: Endothelial
       cells/Pericytes", "Mixed signature: Macrophages/Neurons", "T cells",
       "Neutrophils", "Mixed signature: Astrocytes/Neurons", "Artifacts")
names(new.cluster.ids) <- levels(MERFISH_all_includes_mixed_signature)
MERFISH_all_includes_mixed_signature <-
       RenameIdents(MERFISH_all_includes_mixed_signature, new.cluster.ids)
DimPlot(MERFISH_all_includes_mixed_signature, reduction = "tsne",label=T)
MERFISH_all_includes_mixed_signature[["MERFISH.cluster.names"]] <- Idents(object =
       MERFISH_all_includes_mixed_signature) # stash identity

# remove mixed-signature clusters
MERFISH_all <- subset(x = MERFISH_all_includes_mixed_signature, idents = c("Mixed
       signature: Astrocytes/Neurons", "Mixed signature: Endothelial cells/Pericytes",
       "Mixed signature: Macrophages/Neurons", "Mixed signature: Olfactory ensheathing
       cells/Neurons", "Mixed signature: Astrocytes/Neurons/Endothelial cells",
       "Artifacts"),invert = TRUE)
DimPlot(MERFISH_all, reduction = "tsne",label=T)+ NoLegend()
```

```
# add neuronal subtype to metadata

Idents(object=MERFISH_all) <- MERFISH_all@meta.data$seurat_clusters
DimPlot(MERFISH_all, reduction = "tsne",label=T)
new.cluster.ids <- c("Granule cells", "Periglomerular cells", "NA", "Periglomerular
       cells", "NA", "NA", "External tufted cells", "Mitral/Tufted cells", "NA",
       "Neuroblasts", "Somatostatin-immunoreactive neurons", "NA", "NA", "NA", "NA",
       "NA", "NA", "NA")
names(new.cluster.ids) <- levels(MERFISH_all)
MERFISH_all <- RenameIdents(MERFISH_all, new.cluster.ids)
DimPlot(MERFISH_all, reduction = "tsne",label=T)
MERFISH_all[["MERFISH.neuronal.subtype"]] <- Idents(object = MERFISH_all) # stash
       identity

MERFISH_all$inoculate <- MERFISH_all$group
MERFISH_all$group <- NULL # remove meta data from UMass
MERFISH_all$celltype <- NULL # remove meta data from UMass
MERFISH_all$seurat.cluster <- MERFISH_all$seurat_clusters
MERFISH_all$seurat_clusters <- NULL # remove meta data from UMass

saveRDS(MERFISH_all, file = "Directory/Manuscript/Data/MERFISH_all.rds")

# EC differential expression by biological replicate

Idents(object=MERFISH_all) <- MERFISH_all@meta.data$MERFISH.cluster.names
MERFISH_ECs <- subset(x = MERFISH_all, idents = c("Endothelial cells"),invert = FALSE)
DimPlot(MERFISH_ECs, reduction = "tsne")

 # Note: ComBat-corrected expression values in "data"; original raw values
       (counts/m^3) - which were log-normalized via Seurat - are in "counts"

merfish.probes <- c("Abca7", "Abcc3", "Abcg2", "Ablim1", "Actb", "Acvrl1", "Adam10",
       "Adam17", "Adgrf5", "Adgrl4", "Adora1", "Aff3", "Ago4", "Agt", "Ahr", "Akap12",
       "Aldoc", "Anxa1", "Ap2m1", "Aqp4", "Arc", "Arg1", "Arhgap29", "Arl15", "Arpc2",
       "Atmin", "Atp10a", "Axl", "Baiap2l1", "Bard1", "Bin1", "Birc5", "Bmp6",
       "Brca1", "Btk", "C1qa", "C1qb", "C1qbp", "C1qc", "C3", "C3ar1", "C4a", "C5ar1",
       "Cald1", "Casp7", "Casp8", "Cass4", "Ccl2", "Ccl22", "Ccl3", "Ccr2", "Cd14",
       "Cd163", "Cd27", "Cd33", "Cd3e", "Cd4", "Cd47", "Cd68", "Cd72", "Cd74",
       "Cd79a", "Cd84", "Cd86", "Cd8a", "Cdh5", "Cdh9", "Cemip", "Cenpa", "Cgnl1",
       "Chek2", "Chit1", "Cldn5", "Clec7a", "Clic4", "Clu", "Cmtm8", "Cobll1",
       "Col6a3", "Cotl1", "Crim1", "Csad", "Csf1r", "Csf2", "Csf2ra", "Csf2rb",
       "Cspg4", "Cstb", "Ctgf", "Ctsb", "Cx3cr1", "Cxcl10", "Cyr61", "Dach1", "Dapk1",
       "Dclre1a", "Ddx58", "Des", "Dlc1", "Dna2", "Dock2", "Dock9", "Dusp1", "E2f1",
       "Ebf1", "Ebi3", "Ece1", "Edn1", "Edn3", "Efnb2", "Egfl7", "Egfr", "Egr1",
       "Elovl7", "Emcn", "Emp1", "Eng", "Enpp6", "Entpd1", "Epas1", "Epb41l4a",
       "Epha1", "Erg", "Esam", "Esyt2", "Fancd2", "Fbrs", "Fbxw17", "Fcer1g", "Fcgr1",
       "Fcgrt", "Fcrls", "Fgd2", "Flcn", "Flnb", "Flt1", "Flt3", "Flt4", "Fn1",
       "Folr2", "Fos", "Foxj1", "Foxp1", "Ftsj3", "Gad1", "Gad2", "Galnt18", "Gbp2",
       "Gfer", "Gna13", "Gna15", "Gpi1", "Gpr183", "Gpr34", "Gpr84", "Grb10", "Grn",
       "Gusb","H2.D1", "Hcar2", "Heg1", "Hells", "Hexb", "Hmgb2", "Hmox1", "Trem2",
       "Icam1", "Ifi30", "Ifih1", "Igsf6", "Il1a", "Il1b", "Il1rl2", "Il1rn", "Il21r",
       "Il2rg", "Il3ra", "Impact", "Inpp5d", "Irak4", "Irf7", "Irf8", "Itga1",
       "Itga2", "Itga6", "Itgae", "Itgal", "Itgam", "Itgax", "Itgb1", "Itgb3",
       "Itgb5", "Itm2a", "Jcad", "Jun", "Kif26a", "Kit", "Lacc1", "Lair1", "Lama2",
       "Lcn2", "Ldb2", "Ldlrad3", "Lef1", "Lfng", "Lgals3", "Lgmn", "Lig1", "Liph",
       "Lmnb1", "Lrch3", "Lrp1", "Lrrc3", "Ly6g", "Ly9", "Lyn", "Lyve1", "Lyz2",
```

```
    "Mb21d1", "Mcm2", "Mcm5", "Mcm6", "Mctp1", "Mecom", "Mef2c", "Meg3", "Mertk",
    "Mfsd6l", "Mgat1", "Mki67", "Mkl2", "Mmp2", "Mmp9", "Mpnd", "Mrc1", "Ms4a1",
    "Ms4a6d", "Ms4a7", "Msn", "Mvp", "Mybl2", "Myrip", "Nampt", "Napsa", "Ncaph",
    "Ncf1", "Nckap1l", "Nebl", "Nfib", "Nlrp3", "Nos3", "Nostrin", "Nrm", "Ocln",
    "Olfm2", "Olfml3", "Osm", "Osmr", "P2ry12", "Palmd", "Pam", "Pard3", "Pcna",
    "Pde10a", "Pde4d", "Pdgfra", "Pdgfrb", "Pdlim5", "Pdpn", "Pdzrn3", "Pecam1",
    "Picalm", "Pik3cg", "Pla2g4a", "Plac8", "Plcb4", "Plcg2", "Plekha6", "Plekhg1",
    "Plp1", "Plpp1", "Pltp", "Plxdc2", "Plxna2", "Podxl", "Pole", "Ppfibp1",
    "Prdx5", "Prickle2", "Prkg1", "Pros1", "Psmb8", "Ptk2b", "Ptpn6", "Ptprc",
    "Ptprg", "Ptprj", "Ptprm", "Pvalb", "Qars", "Rad23b", "Rad51", "Rae1",
    "Rapgef4", "Rbms3", "Rngtt", "Rora", "Rorb", "Rrm2", "Rsad2", "Rundc3b",
    "Sall1", "Sardh", "Sash1", "Sdf4", "Sell", "Serpina3n", "Serpine1", "Serpinf1",
    "Siglecf", "Slamf8", "Slamf9", "Slc17a6", "Slc1a1", "Slc39a10", "Slc40a1",
    "Slc4a4", "Slc7a1", "Slco2b1", "Slfn8", "Smc3", "Snx2", "Sorbs1", "Sorbs2",
    "Sox10", "Sox2", "Sox9", "Spi1", "Spp1", "Sptbn1", "Srgn", "Sst", "St8sia6",
    "Stat1", "Syk", "Syne1", "Syne2", "Tacc1", "Tead1", "Tek", "Tgfa", "Tgfbi",
    "Tgfbr1", "Tgfbr2", "Tgm2", "Thsd4", "Timeless", "Timp2", "Timp3", "Tlr2",
    "Tlr4", "Tmem119", "Tmem173", "Tmtc1", "Tnfrsf11a", "Tnfrsf1a", "Tnfsf13b",
    "Top2a", "Trem1", "Trem2", "Trim47", "Tshz2", "Tspan33", "Ttll12", "Ttr",
    "Txnrd1", "Tyrobp", "Unc13b", "Ung", "Utrn", "Vac14", "Vav1", "Vcl", "Vegfa",
    "Vim", "Vip", "Vtn", "Vwf", "Was", "Wwtr1", "Xpo1", "Zbp1", "Zbtb46")

MERFISH_ECs_Original_LogNormalized_AvgExpression <- AverageExpression(MERFISH_ECs,
    features = merfish.probes, slot = "counts")
write.csv(MERFISH_ECs_Original_LogNormalized_AvgExpression, file =
    "Directory/MERFISH/MERFISH_ECs_Original_LogNormalized_AvgExpression_bySample.cs
    v")

# add distance to T cell to microglia

MicrogliaOnly_TCellNNDs <- readRDS("Directory/MERFISH/Data/RDS
    files/220921_AgalliuLab_GAS_combat_MicrogliaOnly_TCellNNDs.rds")
DimPlot(MicrogliaOnly_TCellNNDs)

DistanceToTCell <- MicrogliaOnly_TCellNNDs[["NNDs.MG.TCells"]]
MERFISH_microglia <- AddMetaData(MERFISH_microglia, metadata = DistanceToTCell,
    col.name = "T_cell_distance")


 plot.xy.gene.minimal <- function(object, gene){

 object.df <- data.frame(object$center.x, object$center.y, FetchData(object = object,
    vars = gene, slot = "counts"))
 x <- object$center.x
 y <- object$center.y
 Expression <- object.df[,3]
 mid <- mean(Expression)
 ggplot(object.df, aes(x, y, colour = Expression)) + geom_point(size = 2.5) +
    ggtitle(paste(gene)) + theme_linedraw() + scale_color_gradient(limits =
    c(5,150), na.value = "gray90", high = "light gray", low = "#1e0bfe", labs(title
    = "Distance\nto nearest\nT cell (mm)")) + theme(plot.title = element_blank(),
    axis.title = element_blank(), panel.grid = element_blank(), panel.border =
    element_rect(color = "white"), legend.text = element_text(size = 12),
    axis.ticks = element_blank(), axis.text = element_blank(), legend.title =
    element_text(size = 14, vjust = 5))
 }
```

```
plot.xy.gene.minimal(MERFISH_microglia_GAS_1, "T_cell_distance")
```

```
## code for adding olfactory bulb region to MERFISH data (Columbia) ----

# analysis run with R v 4.0.2 and Seurat 4.0.2

# export coordinate plot of all cells in sample
DimPlot(MERFISH_all)
Idents(object=MERFISH_all) <- MERFISH_all@meta.data$sample
MERFISH_GAS_1 <- subset(x = MERFISH_all, idents = c("GAS_1"),invert = FALSE)

plot.xy.cluster <- function(object){

object.df <- data.frame(object$center.x, object$center.y, object@active.ident)
x <- object$center.x
y <- object$center.y
group <- object@active.ident
ggplot(object.df, aes(x, y, color = group)) + geom_point(size = 0.1) +
       theme_linedraw() + NoLegend()
}

plot.xy.cluster(MERFISH_GAS_1)

ggsave(
filename = "DimPlot_MERFISH_GAS_4_SpatialPlot.jpeg",
plot = last_plot(),
device = "jpeg",
path = "Directory/MERFISH/Assigning microglia to regions",
scale = 1,
width = 5,
height = 5,
units = "in",
dpi = 300,
bg = "transparent"
)

# subset on microglia only

Idents(object=MERFISH_GAS_1) <- MERFISH_GAS_1@meta.data$MERFISH.cluster.names
DimPlot(MERFISH_GAS_1)
MERFISH_GAS_1_microglia <- subset(x = MERFISH_GAS_1, idents = c("Microglia"),invert =
       FALSE)
DimPlot(MERFISH_GAS_1_microglia)

# export microglia coordinates

microglia.GAS_1.info <- MERFISH_GAS_1_microglia@meta.data[c(5, 6, 8, 13)]
head(microglia.GAS_1.info)

microglia_GAS_1.x <- MERFISH_GAS_1_microglia@meta.data[,5]
head(microglia_GAS_1.x)
microglia_GAS_1.y <- MERFISH_GAS_1_microglia@meta.data[,6]
head(microglia_GAS_1.y)

# regional polygon coordinates

# generated using mobilefish.com

granular_GAS_1L.x <- c(1676, 1764, 1829, 1911, 2007, 2077, 2178, 2287, 2409, 2505, 2606, 2676,
       2702, 2720, 2694, 2650, 2601, 2536, 2466, 2388, 2305, 2248, 2199, 2152, 2116, 2108, 2077,
```

```
       2046, 2012, 1968, 1930, 1880, 1816, 1740, 1637, 1549, 1466, 1383, 1331, 1300, 1292, 1313,
       1362, 1388, 1414, 1453, 1471, 1497, 1515, 1528, 1523, 1536, 1575, 1624)
granular_GAS_1L.y <- c(15861, 15894, 15917, 15942, 15964, 15978, 15992, 16003, 16011, 16011,
       15984, 15922, 15847, 15744, 15663, 15594, 15546, 15499, 15463, 15415, 15374, 15335,
       15279, 15218, 15148, 15073, 15003, 14928, 14853, 14786, 14702, 14655, 14608, 14566,
       14546, 14546, 14566, 14613, 14688, 14772, 14861, 14936, 14998, 15053, 15128, 15204,
       15271, 15354, 15443, 15546, 15622, 15702, 15772, 15814)

granular_GAS_1R.x <- c(2981, 3007, 3064, 3152, 3256, 3357, 3453, 3536, 3593, 3658, 3676, 3683,
       3670, 3645, 3588, 3523, 3427, 3339, 3274, 3186, 3095, 3033, 2963, 2893, 2829, 2733, 2650,
       2575, 2518, 2479, 2445, 2445, 2471, 2541, 2593, 2658, 2727, 2797, 2854, 2893, 2924, 2950)
granular_GAS_1R.y <- c(14839, 14936, 15025, 15073, 15081, 15053, 14992, 14917, 14828, 14711,
       14599, 14485, 14360, 14251, 14156, 14073, 13984, 13922, 13867, 13814, 13738, 13669,
       13599, 13538, 13485, 13477, 13524, 13594, 13663, 13744, 13853, 13942, 14053, 14134,
       14195, 14257, 14312, 14354, 14435, 14518, 14599, 14711)

EPL_GAS_1L.x <-c(1326, 1344, 1362, 1401, 1445, 1510, 1624, 1740, 1860, 1981, 2103, 2235, 2375,
       2479, 2601, 2715, 2784, 2829, 2823, 2803, 2759, 2689, 2601, 2523, 2427, 2339, 2274, 2248,
       2212, 2165, 2108, 2051, 1987, 1917, 1847, 1746, 1650, 1562, 1471, 1375, 1305, 1235, 1191,
       1147, 1126, 1139, 1173, 1204, 1230, 1256, 1287)
EPL_GAS_1L.y <-c(15499, 15599, 15730, 15853, 15936, 16003, 16059, 16128, 16170, 16176, 16181,
       16209, 16204, 16148, 16087, 16017, 15922, 15800, 15669, 15574, 15485, 15401, 15335,
       15271, 15204, 15134, 15045, 14956, 14819, 14738, 14663, 14574, 14524, 14471, 14410,
       14374, 14360, 14368, 14410, 14449, 14499, 14552, 14613, 14711, 14805, 14908, 15025,
       15115, 15209, 15293, 15396)

EPL_GAS_1R.x <-c(3880, 3886, 3867, 3841, 3797, 3777, 3772, 3715, 3650, 3606, 3531, 3453, 3396,
       3331, 3248, 3165, 3103, 3007, 2924, 2829, 2720, 2624, 2523, 2471, 2414, 2370, 2331, 2326,
       2326, 2339, 2352, 2396, 2409, 2445, 2497, 2567, 2632, 2681, 2740, 2797, 2810, 2816, 2860,
       2930, 2994, 3077, 3173, 3274, 3370, 3458, 3541, 3611, 3694, 3766, 3816, 3854)
EPL_GAS_1R.y <-c(14627, 14518, 14429, 14326, 14257, 14181, 14067, 13970, 13908, 13839, 13777,
       13725, 13663, 13585, 13538, 13471, 13410, 13368, 13340, 13298, 13312, 13340, 13382,
       13443, 13505, 13580, 13669, 13764, 13867, 13978, 14067, 14142, 14237, 14321, 14388,
       14443, 14510, 14594, 14663, 14758, 14853, 14950, 15053, 15106, 15148, 15162, 15181,
       15195, 15204, 15181, 15128, 15059, 14998, 14903, 14805, 14716)

glomerular_GAS_1L.x <-c(2694, 2619, 2554, 2497, 2445, 2396, 2352, 2305, 2261, 2212, 2139, 2056,
       1981, 1911, 1816, 1668, 1588, 1479, 1388, 1269, 1209, 1126, 1103, 1051, 1025, 994, 1000,
       1000, 1020, 1038, 1069, 1103, 1108, 1147, 1178, 1199, 1217, 1235, 1269, 1292, 1339, 1401,
       1497, 1567, 1694, 1803, 1893, 2038, 2108, 2222, 2331, 2409, 2505, 2588, 2645, 2720, 2764,
       2816, 2860, 2886, 2917, 2943, 2955, 2976, 2976, 2943, 2911, 2867, 2829, 2784)
glomerular_GAS_1L.y <-c(15195, 15128, 15045, 14978, 14889, 14786, 14702, 14627, 14574, 14477,
       14396, 14340, 14284, 14237, 14218, 14209, 14209, 14232, 14245, 14293, 14340, 14443,
       14524, 14613, 14669, 14744, 14839, 14942, 15067, 15156, 15245, 15340, 15421, 15491,
       15566, 15641, 15744, 15839, 15922, 16011, 16092, 16156, 16209, 16265, 16284, 16307,
       16312, 16354, 16354, 16346, 16346, 16335, 16298, 16271, 16232, 16156, 16101, 16017,
       15964, 15889, 15828, 15758, 15674, 15594, 15518, 15449, 15410, 15340, 15265, 15232)

glomerular_GAS_1R.x <-c(3880, 3886, 3867, 3841, 3797, 3777, 3772, 3766, 3810, 3772, 3759, 3715,
       3645, 3601, 3536, 3484, 3414, 3344, 3282, 3217, 3142, 3059, 2950, 2854, 2772, 2670, 2567,
       2445, 2375, 2313, 2256, 2243, 2204, 2173, 2160, 2160, 2186, 2204, 2235, 2292, 2344, 2388,
       2435, 2471, 2510, 2554, 2611, 2670, 2740, 2803, 2873, 2955, 3038, 3152, 3269, 3435, 3536,
       3606, 3676, 3766, 3829, 3860, 3893, 3906)
glomerular_GAS_1R.y <-c(14627, 14518, 14429, 14326, 14257, 14181, 14067, 13984, 13894, 13805,
       13738, 13655, 13613, 13546, 13477, 13421, 13346, 13293, 13257, 13237, 13204, 13176,
       13167, 13167, 13181, 13209, 13251, 13293, 13354, 13429, 13518, 13608, 13688, 13758,
       13853, 13950, 14025, 14115, 14195, 14271, 14340, 14415, 14491, 14574, 14674, 14758,
       14853, 14950, 15031, 15092, 15176, 15271, 15307, 15326, 15321, 15279, 15237, 15167,
       15101, 15039, 14970, 14875, 14772, 14697)

# find layer yes/no using point in polygon

library(sp)

granular_GAS_1L <- point.in.polygon(microglia_GAS_1.x, microglia_GAS_1.y,
       granular_GAS_1L.x, granular_GAS_1L.y)
granular_GAS_1R <- point.in.polygon(microglia_GAS_1.x, microglia_GAS_1.y,
       granular_GAS_1R.x, granular_GAS_1R.y)

EPL_GAS_1L <- point.in.polygon(microglia_GAS_1.x, microglia_GAS_1.y, EPL_GAS_1L.x,
       EPL_GAS_1L.y)
```

```
EPL_GAS_1R <- point.in.polygon(microglia_GAS_1.x, microglia_GAS_1.y, EPL_GAS_1R.x,
       EPL_GAS_1R.y)

glomerular_GAS_1L <- point.in.polygon(microglia_GAS_1.x, microglia_GAS_1.y,
       glomerular_GAS_1L.x, glomerular_GAS_1L.y)
glomerular_GAS_1R <- point.in.polygon(microglia_GAS_1.x, microglia_GAS_1.y,
       glomerular_GAS_1R.x, glomerular_GAS_1R.y)

microglia.GAS_1.info <- cbind.data.frame(microglia.GAS_1.info, granular_GAS_1L,
       granular_GAS_1R, EPL_GAS_1L, EPL_GAS_1R, glomerular_GAS_1L, glomerular_GAS_1R)
head(microglia.GAS_1.info, 50)

# assign layer identity

microglia.GAS_1_granular <- filter(microglia.GAS_1.info, granular_GAS_1L == 1 |
       granular_GAS_1R == 1)
head(microglia.GAS_1_granular, 100)

microglia.GAS_1_EPL <- filter(microglia.GAS_1.info, EPL_GAS_1L == 1 | EPL_GAS_1R == 1,
       granular_GAS_1L == 0, granular_GAS_1R == 0)
head(microglia.GAS_1_EPL, 100)

microglia.GAS_1_glomerular <- filter(microglia.GAS_1.info, glomerular_GAS_1L == 1 |
       glomerular_GAS_1R == 1, EPL_GAS_1L == 0, EPL_GAS_1R == 0, granular_GAS_1L == 0,
       granular_GAS_1R == 0)
head(microglia.GAS_1_glomerular, 100)

microglia.GAS_1_outer <- filter(microglia.GAS_1.info, glomerular_GAS_1L == 0,
       glomerular_GAS_1R == 0, EPL_GAS_1L == 0, EPL_GAS_1R == 0, granular_GAS_1L == 0,
       granular_GAS_1R == 0)
head(microglia.GAS_1_outer, 100)

microglia.GAS_1_granular <- microglia.GAS_1_granular %>% mutate(layer =
       rep("granular"))
microglia.GAS_1_EPL <- microglia.GAS_1_EPL %>% mutate(layer =
       rep("external_plexiform"))
microglia.GAS_1_glomerular <- microglia.GAS_1_glomerular %>% mutate(layer =
       rep("glomerular"))
microglia.GAS_1_outer <- microglia.GAS_1_outer %>% mutate(layer = rep("outer"))

microglia.GAS_1 <- rbind(microglia.GAS_1_granular, microglia.GAS_1_EPL,
       microglia.GAS_1_glomerular, microglia.GAS_1_outer)
head(microglia.GAS_1)

layer_df_GAS_1 <- microglia.GAS_1 %>% select(0, 11)
head(layer_df_GAS_1)

MERFISH_GAS_1_microglia <- AddMetaData(MERFISH_GAS_1_microglia, metadata =
       layer_df_GAS_1, col.name = "layer")
plot.xy.cluster(MERFISH_GAS_1_microglia)
Idents(object=MERFISH_GAS_1_microglia) <- MERFISH_GAS_1_microglia@meta.data$layer
plot.xy.cluster(MERFISH_GAS_1_microglia)

rm(layer_df_GAS_1, MERFISH_GAS_1, microglia.GAS_1, microglia.GAS_1.info,
       microglia.GAS_1_EPL, microglia.GAS_1_glomerular, microglia.GAS_1_granular,
       microglia.GAS_1_outer, microglia_GAS_1.x, microglia_GAS_1.y)
```

```r
rm(granular_GAS_1L, granular_GAS_1L.x, granular_GAS_1L.y, granular_GAS_1R,
       granular_GAS_1R.x, granular_GAS_1R.y)
rm(EPL_GAS_1L, EPL_GAS_1L.x, EPL_GAS_1L.y, EPL_GAS_1R, EPL_GAS_1R.x, EPL_GAS_1R.y)
rm(glomerular_GAS_1L, glomerular_GAS_1L.x, glomerular_GAS_1L.y, glomerular_GAS_1R,
       glomerular_GAS_1R.x, glomerular_GAS_1R.y)

# repeat for other samples

# regional polygon coordinates for PBS 1

granular_PBS_1L.x <- c(1271, 1398, 1519, 1665, 1786, 1880, 2000, 2095, 2160, 2170, 2095, 2023,
       1913, 1776, 1655, 1528, 1414, 1320, 1265, 1199, 1118, 1046, 952, 831, 744, 688, 688, 711,
       750, 783, 815, 871, 936, 1001, 1079, 1160)
granular_PBS_1L.y <- c(4100, 4143, 4173, 4182, 4173, 4128, 4091, 4025, 3938, 3826, 3745, 3685,
       3625, 3594, 3558, 3537, 3477, 3417, 3350, 3278, 3212, 3158, 3091, 3085, 3131, 3203, 3299,
       3396, 3486, 3552, 3640, 3736, 3832, 3923, 3989, 4040)
granular_PBS_1R.x <- c(3059, 3137, 3241, 3368, 3472, 3554, 3586, 3586, 3593, 3593, 3586, 3593,
       3586, 3577, 3531, 3489, 3443, 3362, 3290, 3202, 3104, 2984, 2896, 2808, 2704, 2626, 2538,
       2450, 2391, 2391, 2408, 2456, 2489, 2544, 2593, 2648, 2720, 2785, 2841, 2889, 2955)
granular_PBS_1R.y <- c(3131, 3158, 3188, 3188, 3197, 3197, 3121, 3040, 2959, 2847, 2787, 2715,
       2625, 2513, 2411, 2335, 2284, 2239, 2233, 2233, 2233, 2233, 2218, 2188, 2188, 2167, 2167,
       2197, 2248, 2344, 2441, 2498, 2567, 2634, 2700, 2766, 2817, 2878, 2944, 3019, 3085)
EPL_PBS_1L.x <- c(1255, 1343, 1414, 1535, 1655, 1760, 1831, 1903, 2017, 2121, 2215, 2258, 2281,
       2287, 2281, 2264, 2215, 2144, 2072, 2000, 1919, 1847, 1776, 1665, 1577, 1479, 1447, 1385,
       1326, 1271, 1206, 1144, 1079, 1007, 919, 831, 734, 656, 584, 535, 512, 503, 519, 535,
       558, 568, 600, 662, 704, 766, 809, 864, 903, 975, 1040, 1144, 1193)
EPL_PBS_1L.y <- c(4350, 4426, 4477, 4507, 4498, 4492, 4432, 4359, 4320, 4293, 4212, 4121, 4010,
       3914, 3811, 3721, 3655, 3588, 3537, 3492, 3441, 3402, 3390, 3359, 3359, 3350, 3293, 3233,
       3167, 3115, 3070, 3025, 2974, 2929, 2908, 2908, 2929, 2974, 3025, 3106, 3197, 3269, 3350,
       3432, 3507, 3588, 3676, 3721, 3787, 3847, 3914, 3980, 4040, 4128, 4197, 4233, 4278)
EPL_PBS_1R.x <- c(2873, 2961, 3033, 3121, 3202, 3306, 3433, 3547, 3681, 3739, 3769, 3795, 3850,
       3883, 3883, 3857, 3857, 3857, 3840, 3840, 3827, 3785, 3739, 3674, 3577, 3498, 3394, 3290,
       3042, 2929, 2818, 2714, 2593, 2495, 2385, 2281, 2225, 2215, 2199, 2209, 2232, 2248, 2287,
       2329, 2391, 2440, 2489, 2544, 2609, 2665, 2730, 2792)
EPL_PBS_1R.y <- c(3224, 3293, 3350, 3381, 3396, 3402, 3411, 3402, 3381, 3320, 3224, 3106, 3004,
       2878, 2781, 2670, 2573, 2492, 2381, 2293, 2203, 2115, 2046, 1980, 1959, 1959, 1959, 1959,
       1974, 1959, 1953, 1938, 1938, 1929, 1944, 1989, 2070, 2143, 2218, 2293, 2365, 2462, 2528,
       2588, 2664, 2721, 2796, 2868, 2938, 2995, 3055, 3121)
glomerular_PBS_1L.x <- c(1919, 2023, 2127, 2241, 2297, 2346, 2401, 2434, 2456, 2489, 2512, 2479,
       2440, 2346, 2225, 2095, 1968, 1864, 1727, 1616, 1528, 1440, 1369, 1287, 1193, 1072, 942,
       822, 672, 568, 480, 431, 366, 333, 304, 304, 320, 327, 333, 376, 437, 480, 542, 591, 639,
       678, 718, 792, 864, 926, 1001, 1063, 1112, 1199, 1326, 1447, 1551, 1688, 1847)
glomerular_PBS_1L.y <- c(4507, 4486, 4462, 4411, 4344, 4278, 4218, 4167, 4070, 3944, 3826, 3676,
       3528, 3426, 3359, 3329, 3293, 3218, 3173, 3100, 3025, 2959, 2899, 2847, 2772, 2721, 2736,
       2772, 2781, 2811, 2878, 2944, 3010, 3070, 3173, 3233, 3329, 3402, 3486, 3567, 3634, 3691,
       3772, 3878, 3950, 4034, 4106, 4188, 4254, 4335, 4375, 4456, 4513, 4573, 4603, 4609, 4609,
       4603, 4567)
glomerular_PBS_1R.x <- c(3114, 3192, 3280, 3384, 3515, 3619, 3697, 3811, 3922, 4019, 4065, 4098,
       4137, 4146, 4153, 4146, 4114, 4091, 4075, 4042, 3987, 3922, 3840, 3762, 3635, 3498, 3394,
       3280, 3176, 3082, 2945, 2834, 2736, 2616, 2538, 2456, 2375, 2287, 2215, 2154, 2072, 2056,
       2040, 2049, 2066, 2121, 2170, 2209, 2264, 2329, 2368, 2417, 2495, 2570, 2642, 2730, 2808,
       2896, 2977, 3059)
glomerular_PBS_1R.y <- c(3507, 3528, 3543, 3543, 3522, 3477, 3426, 3359, 3284, 3167, 3034, 2929,
       2817, 2721, 2609, 2507, 2396, 2293, 2167, 2055, 1980, 1899, 1811, 1745, 1745, 1721, 1685,
       1664, 1664, 1670, 1691, 1685, 1634, 1649, 1706, 1760, 1826, 1878, 1953, 2019, 2076, 2152,
       2239, 2308, 2396, 2456, 2543, 2640, 2721, 2766, 2856, 2959, 3019, 3091, 3158, 3263, 3329,
       3390, 3447, 3477)

# regional polygon coordinates for GAS 2

granular_GAS_2L.x <- c(752, 844, 957, 1055, 1165, 1248, 1331, 1438, 1536, 1624, 1716, 1805, 1903,
       1985, 2077, 2160, 2236, 2319, 2392, 2460, 2573, 2640, 2695, 2701, 2671, 2604, 2527, 2460,
       2371, 2288, 2221, 2145, 2062, 1985, 1912, 1814, 1738, 1670, 1597, 1514, 1429, 1324, 1242,
       1144, 1061, 985, 902, 835, 768, 706, 654, 624, 608, 596, 596, 624, 679)
granular_GAS_2L.y <- c(14669, 14711, 14736, 14736, 14736, 14711, 14695, 14711, 14685, 14659,
       14643, 14633, 14618, 14618, 14643, 14685, 14757, 14809, 14860, 14860, 14860, 14772,
       14643, 14504, 14355, 14287, 14277, 14267, 14226, 14174, 14138, 14086, 14024, 13973,
       13921, 13885, 13844, 13797, 13746, 13694, 13679, 13653, 13617, 13606, 13591, 13606,
       13617, 13653, 13694, 13771, 13844, 13962, 14086, 14251, 14391, 14504, 14592)
```

```r
granular_GAS_2R.x <- c(2454, 2552, 2634, 2717, 2799, 2891, 2980, 3072, 3154, 3237, 3289, 3356,
        3408, 3454, 3460, 3445, 3402, 3350, 3289, 3212, 3154, 3063, 2986, 2913, 2836, 2753, 2671,
        2613, 2552, 2475, 2392, 2325, 2258, 2190, 2114, 2041, 1964, 1888, 1820, 1753, 1670, 1603,
        1551, 1490, 1453, 1438, 1468, 1542, 1609, 1695, 1790, 1866, 1933, 2041, 2114, 2197, 2279,
        2362)
granular_GAS_2R.y <- c(10887, 10928, 10964, 11016, 11068, 11119, 11119, 11145, 11130, 11119,
        11078, 11006, 10939, 10825, 10675, 10557, 10443, 10381, 10294, 10216, 10154, 10113,
        10077, 10051, 10025, 10041, 10041, 10067, 10092, 10103, 10165, 10216, 10268, 10304,
        10330, 10345, 10345, 10330, 10330, 10345, 10345, 10330, 10381, 10433, 10536, 10634,
        10737, 10825, 10866, 10902, 10902, 10913, 10902, 10887, 10877, 10866, 10866, 10866)
EPL_GAS_2L.x <- c(618, 679, 761, 872, 994, 1107, 1220, 1324, 1444, 1536, 1640, 1753, 1875, 1973,
        2068, 2138, 2212, 2319, 2423, 2527, 2619, 2695, 2738, 2763, 2769, 2753, 2723, 2656, 2597,
        2521, 2454, 2362, 2279, 2206, 2099, 1995, 1881, 1777, 1679, 1566, 1459, 1370, 1248, 1113,
        970, 865, 761, 679, 624, 566, 498, 474, 483, 498, 541)
EPL_GAS_2L.y <- c(14747, 14809, 14886, 14912, 14938, 14963, 14963, 14963, 14963, 14963, 14948,
        14922, 14886, 14896, 14948, 15000, 15041, 15087, 15154, 15165, 15041, 14922, 14757,
        14633, 14478, 14329, 14189, 14164, 14164, 14112, 14035, 13973, 13921, 13885, 13833,
        13782, 13730, 13679, 13606, 13539, 13493, 13441, 13441, 13441, 13441, 13477, 13555,
        13632, 13704, 13818, 13962, 14138, 14380, 14530, 14659)
EPL_GAS_2R.x <- c(2613, 2708, 2799, 2897, 3001, 3124, 3222, 3326, 3393, 3485, 3522, 3515, 3500,
        3454, 3387, 3320, 3252, 3182, 3078, 2995, 2903, 2815, 2708, 2588, 2475, 2356, 2212, 2108,
        1979, 1875, 1738, 1597, 1520, 1401, 1324, 1257, 1227, 1272, 1355, 1444, 1551, 1640, 1707,
        1829, 1927, 2025, 2145, 2249, 2356, 2460, 2527)
EPL_GAS_2R.y <- c(11320, 11372, 11372, 11372, 11357, 11320, 11284, 11181, 11119, 11006, 10789,
        10650, 10443, 10278, 10154, 10051, 9963, 9901, 9834, 9809, 9772, 9772, 9783, 9809, 9850,
        9876, 9963, 10000, 10041, 10041, 10092, 10139, 10190, 10278, 10417, 10510, 10686, 10825,
        10928, 10990, 11052, 11078, 11119, 11130, 11145, 11145, 11119, 11145, 11155, 11207,
        11243)
glomerular_GAS_2L.x <- c(1499, 1664, 1783, 1897, 2010, 2138, 2249, 2347, 2454, 2527, 2649, 2695,
        2769, 2769, 2784, 2747, 2695, 2613, 2552, 2490, 2334, 2242, 2138, 2010, 1875, 1722, 1572,
        1416, 1263, 1113, 957, 798, 679, 581, 498, 468, 422, 422, 437, 443, 483, 541, 602, 670,
        761, 859, 979, 1098, 1220, 1346, 1429)
glomerular_GAS_2L.y <- c(15294, 15227, 15190, 15180, 15180, 15201, 15242, 15319, 15319, 15242,
        15077, 14922, 14736, 14556, 14391, 14303, 14200, 14050, 13947, 13869, 13730, 13668,
        13581, 13493, 13426, 13364, 13302, 13250, 13235, 13209, 13250, 13302, 13452, 13565,
        13730, 13921, 14086, 14287, 14442, 14633, 14809, 14989, 15113, 15190, 15242, 15216,
        15216, 15216, 15242, 15252, 15252)
glomerular_GAS_2R.x <- c(1820, 1927, 2025, 2160, 2279, 2417, 2567, 2701, 2845, 2980, 3124, 3243,
        3335, 3378, 3445, 3515, 3515, 3558, 3589, 3552, 3500, 3417, 3350, 3222, 3108, 2934, 2793,
        2695, 2567, 2408, 2264, 2129, 2025, 1888, 1738, 1597, 1499, 1361, 1288, 1175, 1113, 1083,
        1092, 1129, 1175, 1288, 1386, 1526, 1624, 1716)
glomerular_GAS_2R.y <- c(11372, 11398, 11449, 11460, 11460, 11475, 11537, 11573, 11589, 11589,
        11573, 11511, 11424, 11331, 11217, 11052, 10866, 10650, 10443, 10294, 10165, 10000, 9809,
        9711, 9607, 9571, 9530, 9556, 9597, 9633, 9669, 9695, 9736, 9762, 9798, 9850, 9912, 9974,
        10051, 10165, 10330, 10510, 10737, 10902, 11078, 11171, 11284, 11331, 11346, 11372)

# regional polygon coordinates for PBS 2

granular_PBS_2L.x <- c(1965, 2117, 2221, 2319, 2418, 2516, 2600, 2668, 2755, 2827, 2915, 2995,
        3067, 3154, 3147, 3117, 3049, 2982, 2920, 2847, 2780, 2713, 2645, 2578, 2521, 2448, 2374,
        2294, 2226, 2154, 2079, 1995, 1920, 1865, 1810, 1773, 1738, 1700, 1688, 1683, 1718, 1773,
        1852, 1915)
granular_PBS_2L.y <- c(2915, 2907, 2894, 2864, 2822, 2792, 2771, 2729, 2686, 2656, 2605, 2571,
        2521, 2448, 2342, 2249, 2206, 2164, 2143, 2121, 2113, 2092, 2049, 2028, 1985, 1947, 1926,
        1913, 1892, 1913, 1947, 1998, 2070, 2143, 2219, 2334, 2414, 2508, 2605, 2707, 2780, 2801,
        2856, 2864)
granular_PBS_2R.x <- c(2431, 2528, 2625, 2713, 2805, 2885, 2995, 3099, 3179, 3246, 3314, 3386,
        3483, 3563, 3638, 3680, 3693, 3680, 3650, 3576, 3503, 3411, 3336, 3251, 3167, 3079, 2982,
        2895, 2827, 2743, 2650, 2583, 2466, 2406, 2326, 2264, 2204, 2154, 2137, 2137, 2167, 2221,
        2264, 2301, 2351, 2394)
granular_PBS_2R.y <- c(4996, 5026, 5056, 5056, 5068, 5056, 5047, 5005, 4954, 4903, 4839, 4805,
        4775, 4712, 4652, 4546, 4453, 4359, 4283, 4253, 4245, 4253, 4266, 4274, 4295, 4317, 4304,
        4304, 4304, 4295, 4283, 4295, 4295, 4295, 4295, 4346, 4380, 4431, 4516, 4618, 4712, 4784,
        4818, 4869, 4920, 4962)
EPL_PBS_2L.x <- c(2907, 3007, 3117, 3214, 3301, 3406, 3503, 3608, 3675, 3768, 3870, 3915, 3895,
        3847, 3790, 3718, 3655, 3571, 3473, 3374, 3306, 3234, 3159, 3054, 2927, 2810, 2713, 2620,
        2516, 2441, 2319, 2234, 2142, 2062, 2012, 2007, 2032, 2062, 2112, 2179, 2246, 2331, 2423,
        2511, 2600, 2688, 2768, 2852)
EPL_PBS_2L.y <- c(5276, 5268, 5225, 5183, 5119, 5017, 4941, 4869, 4784, 4690, 4567, 4419, 4304,
        4181, 4109, 4053, 4024, 4015, 4015, 4015, 4066, 4130, 4109, 4109, 4109, 4147, 4147, 4138,
        4138, 4130, 4147, 4168, 4223, 4295, 4397, 4554, 4690, 4826, 4911, 5005, 5056, 5132, 5183,
        5213, 5247, 5247, 5255, 5268)
```

```r
EPL_PBS_2R.x <- c(2680, 2768, 2865, 2957, 3042, 3134, 3226, 3306, 3361, 3386, 3386, 3344, 3276,
        3214, 3154, 3099, 3032, 2952, 2852, 2793, 2693, 2620, 2546, 2453, 2381, 2301, 2209, 2117,
        2032, 1940, 1877, 1810, 1738, 1675, 1613, 1583, 1566, 1546, 1536, 1541, 1558, 1620, 1670,
        1725, 1785, 1847, 1927, 2037, 2117, 2214, 2284, 2374, 2461, 2533, 2588)
EPL_PBS_2R.y <- c(2886, 2864, 2835, 2801, 2771, 2741, 2665, 2605, 2508, 2406, 2300, 2206, 2164,
        2113, 2070, 2041, 1998, 1977, 1947, 1926, 1892, 1833, 1799, 1769, 1705, 1676, 1663, 1663,
        1684, 1748, 1799, 1850, 1926, 1998, 2083, 2185, 2312, 2427, 2563, 2678, 2771, 2835, 2928,
        2979, 3000, 3009, 3030, 3043, 3030, 3022, 3000, 2971, 2949, 2915, 2907)
glomerular_PBS_2L.x <- c(2895, 2987, 3092, 3201, 3306, 3416, 3503, 3600, 3680, 3748, 3840, 3907,
        3982, 4079, 4142, 4177, 4154, 4084, 4012, 3945, 3870, 3790, 3718, 3650, 3558, 3478, 3374,
        3294, 3209, 3079, 2995, 2902, 2815, 2705, 2613, 2516, 2431, 2344, 2264, 2159, 2079, 2000,
        1932, 1860, 1840, 1840, 1847, 1872, 1920, 1970, 2024, 2067, 2137, 2197, 2251, 2319, 2394,
        2466, 2546, 2633, 2700, 2755, 2827)
glomerular_PBS_2L.y <- c(5506, 5476, 5421, 5349, 5306, 5234, 5183, 5111, 5047, 4983, 4890, 4826,
        4733, 4610, 4504, 4389, 4245, 4147, 4087, 3981, 3896, 3837, 3773, 3731, 3701, 3659, 3637,
        3637, 3637, 3688, 3709, 3743, 3752, 3760, 3794, 3837, 3879, 3896, 3952, 4024, 4087, 4138,
        4253, 4380, 4516, 4682, 4805, 4911, 5005, 5098, 5170, 5225, 5289, 5340, 5370, 5404, 5455,
        5476, 5497, 5484, 5484, 5484, 5484)
glomerular_PBS_2R.x <- c(2578, 2688, 2780, 2902, 3019, 3104, 3172, 3226, 3326, 3381, 3436, 3508,
        3533, 3528, 3448, 3336, 3251, 3154, 3062, 2982, 2902, 2798, 2700, 2608, 2491, 2399, 2314,
        2197, 2092, 1987, 1897, 1780, 1713, 1638, 1566, 1503, 1456, 1431, 1389, 1381, 1381, 1381,
        1411, 1448, 1503, 1541, 1596, 1663, 1725, 1810, 1885, 1977, 2074, 2184, 2289, 2374, 2448,
        2498)
glomerular_PBS_2R.y <- c(3157, 3123, 3073, 3043, 2971, 2907, 2886, 2864, 2780, 2707, 2584, 2436,
        2270, 2083, 1956, 1905, 1841, 1769, 1684, 1642, 1548, 1455, 1391, 1383, 1340, 1340, 1340,
        1332, 1340, 1349, 1404, 1455, 1527, 1599, 1684, 1769, 1884, 2028, 2164, 2300, 2457, 2584,
        2699, 2780, 2864, 2971, 3043, 3094, 3145, 3166, 3179, 3200, 3208, 3208, 3221, 3208, 3187,
        3157)

# regional polygon coordinates for GAS 3

granular_GAS_3L.x <- c(4201, 4274, 4331, 4393, 4464, 4540, 4616, 4692, 4762, 4854, 4949, 5006,
        5055, 5069, 5090, 5088, 5052, 5014, 4944, 4895, 4846, 4789, 4727, 4670, 4624, 4569, 4510,
        4423, 4361, 4287, 4220, 4149, 4089, 4005, 3913, 3823, 3726, 3669, 3623, 3579, 3552, 3547,
        3566, 3593, 3642, 3696, 3761, 3832, 3913, 3983, 4040, 4100, 4144)
granular_GAS_3L.y <- c(2566, 2625, 2680, 2722, 2763, 2814, 2826, 2839, 2835, 2793, 2742, 2688,
        2600, 2516, 2415, 2311, 2214, 2160, 2118, 2055, 1988, 1946, 1883, 1807, 1753, 1719, 1665,
        1631, 1589, 1577, 1548, 1535, 1527, 1518, 1518, 1556, 1615, 1698, 1774, 1858, 1916, 2034,
        2147, 2244, 2336, 2415, 2453, 2495, 2516, 2537, 2537, 2529, 2495)
granular_GAS_3R.x <- c(3970, 4040, 4127, 4214, 4290, 4366, 4447, 4559, 4673, 4768, 4873, 4914,
        4930, 4941, 4922, 4895, 4841, 4781, 4721, 4643, 4567, 4491, 4415, 4350, 4274, 4176, 4103,
        4038, 3970, 3913, 3832, 3753, 3682, 3623, 3579, 3530, 3498, 3490, 3495, 3517, 3547, 3579,
        3609, 3647, 3699, 3734, 3775, 3832, 3872, 3921)
granular_GAS_3R.y <- c(4193, 4206, 4223, 4223, 4223, 4185, 4126, 4076, 4013, 4000, 3979, 3900,
        3816, 3707, 3615, 3539, 3472, 3430, 3409, 3397, 3388, 3384, 3384, 3388, 3376, 3355, 3321,
        3292, 3279, 3271, 3258, 3254, 3271, 3292, 3329, 3405, 3485, 3568, 3652, 3732, 3807, 3862,
        3912, 3967, 4013, 4055, 4093, 4126, 4151, 4172)
EPL_GAS_3L.x <- c(4192, 4274, 4361, 4428, 4483, 4540, 4610, 4678, 4754, 4838, 4914, 4971, 5020,
        5082, 5137, 5172, 5207, 5250, 5248, 5256, 5242, 5237, 5212, 5139, 5101, 5052, 4990, 4917,
        4852, 4789, 4757, 4708, 4645, 4588, 4510, 4423, 4344, 4276, 4184, 4127, 4065, 3970, 3913,
        3856, 3794, 3726, 3658, 3601, 3539, 3484, 3446, 3411, 3392, 3392, 3419, 3460, 3503, 3593,
        3650, 3707, 3783, 3878, 3948, 4002, 4081, 4127)
EPL_GAS_3L.y <- c(2805, 2818, 2860, 2893, 2927, 2977, 2986, 2990, 3011, 2998, 2965, 2914, 2851,
        2797, 2738, 2680, 2642, 2571, 2491, 2374, 2298, 2193, 2105, 2034, 1979, 1916, 1862, 1803,
        1782, 1711, 1657, 1610, 1539, 1518, 1497, 1464, 1418, 1409, 1409, 1405, 1422, 1438, 1438,
        1438, 1451, 1464, 1506, 1539, 1581, 1665, 1749, 1837, 1967, 2072, 2193, 2290, 2407, 2503,
        2550, 2612, 2633, 2663, 2680, 2709, 2738, 2763)
EPL_GAS_3R.x <- c(3796, 3867, 3927, 3989, 4046, 4095, 4152, 4228, 4295, 4366, 4437, 4534, 4602,
        4681, 4735, 4789, 4865, 4936, 5004, 5061, 5118, 5126, 5131, 5104, 5082, 5039, 4993, 4944,
        4887, 4816, 4732, 4651, 4580, 4499, 4409, 4336, 4247, 4176, 4095, 4002, 3921, 3815, 3704,
        3615, 3539, 3468, 3414, 3370, 3338, 3343, 3370, 3397, 3433, 3476, 3536, 3587, 3634, 3690,
        3739)
EPL_GAS_3R.y <-c(4365, 4378, 4399, 4420, 4432, 4441, 4449, 4449, 4441, 4420, 4374, 4340, 4290,
        4235, 4181, 4172, 4151, 4139, 4093, 4030, 3975, 3862, 3753, 3644, 3577, 3514, 3443, 3397,
        3342, 3321, 3292, 3254, 3225, 3195, 3195, 3183, 3149, 3162, 3141, 3095, 3086, 3082, 3086,
        3116, 3149, 3195, 3288, 3388, 3493, 3631, 3766, 3879, 3988, 4072, 4160, 4214, 4264, 4298,
        4332)
glomerular_GAS_3L.x <- c(4005, 4065, 4127, 4184, 4255, 4344, 4428, 4521, 4607, 4678, 4768, 4846,
        4909, 4976, 5033, 5096, 5153, 5207, 5256, 5297, 5348, 5375, 5389, 5416, 5394, 5367, 5348,
        5299, 5242, 5174, 5123, 5063, 4993, 4930, 4873, 4811, 4762, 4700, 4632, 4559, 4483, 4380,
        4309, 4228, 4144, 4054, 3962, 3878, 3788, 3718, 3647, 3587, 3530, 3476, 3425, 3379, 3357,
        3324, 3303, 3275, 3275, 3281, 3300, 3335, 3357, 3397, 3503, 3547, 3587, 3636, 3696, 3756,
        3796, 3851, 3908, 3962)
```

```
glomerular_GAS_3L.y <- c(2881, 2910, 2927, 2948, 2977, 3011, 3053, 3082, 3107, 3116, 3120, 3128,
        3128, 3095, 3065, 3023, 2977, 2893, 2797, 2738, 2675, 2600, 2516, 2394, 2256, 2126, 2034,
        1937, 1849, 1774, 1690, 1602, 1527, 1472, 1418, 1363, 1321, 1288, 1258, 1216, 1195, 1170,
        1162, 1162, 1162, 1174, 1191, 1225, 1237, 1258, 1279, 1313, 1342, 1418, 1493, 1577, 1652,
        1732, 1824, 1916, 2051, 2130, 2223, 2290, 2407, 2508, 2587, 2667, 2730, 2755, 2772, 2776,
        2784, 2797, 2805, 2847)
glomerular_GAS_3R.x <- c(3696, 3745, 3794, 3859, 3918, 3975, 4051, 4157, 4298, 4409, 4569, 4692,
        4803, 4895, 4985, 5063, 5166, 5242, 5278, 5297, 5297, 5283, 5248, 5215, 5174, 5109, 5061,
        4966, 4887, 4784, 4700, 4632, 4567, 4496, 4420, 4358, 4287, 4225, 4171, 4114, 4040, 3975,
        3880, 3794, 3690, 3609, 3517, 3449, 3370, 3313, 3275, 3259, 3265, 3300, 3335, 3387, 3454,
        3509, 3566, 3623)
glomerular_GAS_3R.y <- c(4516, 4529, 4550, 4558, 4612, 4642, 4654, 4646, 4592, 4537, 4512, 4491,
        4449, 4420, 4340, 4264, 4168, 4084, 3954, 3862, 3774, 3686, 3594, 3506, 3397, 3309, 3246,
        3170, 3128, 3120, 3120, 3116, 3095, 3044, 3011, 2990, 2969, 2965, 2965, 2956, 2923, 2914,
        2914, 2902, 2902, 2944, 2986, 3044, 3120, 3233, 3363, 3548, 3728, 3912, 4051, 4181, 4264,
        4353, 4394, 4441)

# regional polygon coordinates for PBS 3

granular_PBS_3L.x <- c(1774, 1886, 1989, 2132, 2250, 2357, 2475, 2596, 2693, 2782, 2857, 2930,
        3012, 3027, 3006, 2982, 2900, 2833, 2766, 2678, 2602, 2536, 2454, 2363, 2290, 2208, 2111,
        2014, 1932, 1850, 1753, 1701, 1631, 1574, 1534, 1498, 1476, 1467, 1534, 1616, 1686)
granular_PBS_3L.y <-c(11294, 11303, 11294, 11303, 11308, 11344, 11357, 11377, 11377, 11377,
        11357, 11322, 11256, 11168, 11086, 10998, 10957, 10910, 10877, 10822, 10776, 10729,
        10674, 10633, 10581, 10553, 10545, 10545, 10553, 10594, 10633, 10682, 10743, 10803,
        10872, 10979, 11086, 11154, 11234, 11270, 11289)
granular_PBS_3R.x <- c(970, 1073, 1155, 1246, 1334, 1452, 1549, 1631, 1737, 1880, 1968, 2035,
        2065, 2050, 2014, 1953, 1871, 1789, 1656, 1574, 1461, 1401, 1319, 1222, 1134, 1043, 924,
        833, 818, 827, 812, 791, 797, 833, 909)
granular_PBS_3R.y <- c(12552, 12604, 12631, 12659, 12686, 12713, 12727, 12741, 12741, 12733,
        12686, 12612, 12505, 12390, 12288, 12214, 12145, 12112, 12079, 12024, 11983, 11942,
        11890, 11835, 11775, 11769, 11775, 11830, 11923, 12011, 12093, 12206, 12315, 12403,
        12497)
EPL_PBS_3L.x <- c(2423, 2551, 2654, 2736, 2879, 2997, 3103, 3191, 3231, 3273, 3282, 3191, 3134,
        3043, 2945, 2833, 2693, 2618, 2551, 2460, 2335, 2229, 2080, 1968, 1826, 1713, 1646, 1522,
        1431, 1355, 1334, 1319, 1334, 1355, 1446, 1574, 1737, 1865, 1998, 2126, 2238, 2326)
EPL_PBS_3L.y <- c(11497, 11497, 11497, 11497, 11506, 11456, 11404, 11335, 11242, 11141, 11045,
        10979, 10918, 10836, 10776, 10696, 10600, 10506, 10452, 10391, 10345, 10270, 10262,
        10276, 10309, 10358, 10424, 10493, 10608, 10734, 10877, 11006, 11119, 11228, 11357,
        11432, 11470, 11478, 11484, 11484, 11478, 11478)
EPL_PBS_3R.x <- c(1498, 1589, 1671, 1774, 1850, 1938, 2029, 2065, 2132, 2193, 2229, 2250, 2250,
        2184, 2111, 2044, 1917, 1835, 1728, 1640, 1543, 1452, 1355, 1258, 1134, 1012, 848, 751,
        678, 618, 557, 514, 499, 499, 514, 551, 618, 693, 791, 873, 976, 1088, 1206, 1313, 1401)
EPL_PBS_3R.y <-c(12969, 12969, 12955, 12922, 12889, 12853, 12793, 12741, 12681, 12604, 12524,
        12403, 12307, 12228, 12159, 12093, 12044, 11983, 11937, 11895, 11849, 11788, 11755,
        11742, 11706, 11706, 11742, 11780, 11895, 11992, 12066, 12173, 12315, 12409, 12491,
        12584, 12667, 12727, 12760, 12793, 12842, 12895, 12922, 12963, 12963)
glomerular_PBS_3L.x <- c(2335, 2445, 2572, 2721, 2863, 2991, 3140, 3258, 3355, 3431, 3492, 3543,
        3543, 3492, 3385, 3267, 3161, 3058, 2909, 2766, 2669, 2490, 2378, 2244, 2014, 1841, 1686,
        1534, 1425, 1288, 1222, 1176, 1149, 1161, 1222, 1304, 1395, 1534, 1671, 1865, 2014, 2162)
glomerular_PBS_3L.y <- c(11681, 11673, 11673, 11673, 11668, 11659, 11607, 11552, 11470, 11368,
        11270, 11127, 10998, 10877, 10770, 10674, 10608, 10553, 10438, 10323, 10235, 10183,
        10108, 10026, 9999, 10040, 10095, 10188, 10303, 10446, 10581, 10729, 10896, 11045, 11206,
        11330, 11437, 11539, 11594, 11626, 11654, 11673)
glomerular_PBS_3R.x <- c(1215, 1355, 1513, 1701, 1880, 1998, 2111, 2250, 2348, 2490, 2566, 2581,
        2566, 2475, 2378, 2296, 2193, 2080, 1953, 1795, 1662, 1543, 1431, 1297, 1149, 982, 797,
        693, 581, 484, 402, 311, 296, 305, 335, 417, 499, 669, 818, 954, 1058)
glomerular_PBS_3R.y <- c(13158, 13186, 13186, 13158, 13131, 13079, 13010, 12941, 12875, 12788,
        12700, 12571, 12395, 12247, 12145, 12085, 12019, 11923, 11830, 11761, 11687, 11626,
        11572, 11511, 11506, 11506, 11552, 11618, 11714, 11808, 11923, 12085, 12247, 12450,
        12612, 12779, 12922, 13010, 13079, 13111, 13131)

# regional polygon coordinates for GAS 4

granular_GAS_4L.x <- c(6639, 6713, 6788, 6860, 6950, 7057, 7153, 7243, 7323, 7425, 7521, 7631,
        7721, 7787, 7846, 7888, 7903, 7882, 7852, 7837, 7793, 7712, 7616, 7536, 7485, 7425, 7500,
        7574, 7691, 7808, 7897, 7993, 7969, 7918, 7867, 7787, 7712, 7631, 7559, 7470, 7374, 7272,
        7183, 7102, 7006, 6911, 6815, 6698, 6618, 6528, 6447, 6382, 6325, 6271, 6220, 6169, 6154,
        6154, 6169, 6175, 6220, 6250, 6316, 6367, 6432, 6492, 6558)
granular_GAS_4L.y <- c(3959, 4037, 4068, 4076, 4076, 4111, 4133, 4155, 4198, 4237, 4237, 4216,
        4176, 4120, 4024, 3929, 3811, 3694, 3555, 3459, 3385, 3350, 3320, 3268, 3268, 3190, 3094,
        3094, 3072, 3085, 3094, 3085, 3020, 2933, 2837, 2772, 2742, 2690, 2624, 2559, 2516, 2498,
```

```r
        2455, 2420, 2390, 2376, 2376, 2433, 2455, 2476, 2529, 2594, 2655, 2711, 2785, 2894, 2976,
        3107, 3203, 3320, 3424, 3533, 3629, 3724, 3803, 3855, 3920)
granular_GAS_4R.x <- c(6256, 6280, 6325, 6391, 6447, 6522, 6624, 6698, 6773, 6854, 6941, 7015,
        7087, 7168, 7243, 7323, 7398, 7479, 7574, 7631, 7685, 7727, 7736, 7721, 7685, 7616, 7559,
        7494, 7425, 7359, 7272, 6830, 6516)
granular_GAS_4R.y <- c(5050, 4946, 4837, 4785, 4729, 4711, 4698, 4711, 4720, 4742, 4794, 4807,
        4816, 4837, 4850, 4868, 4881, 4903, 4946, 4985, 5063, 5159, 5285, 5403, 5490, 5598, 5659,
        5650, 5637, 5637, 5637, 5459, 5276)
EPL_GAS_4L.x <-c(6734, 6824, 6911, 7000, 7111, 7213, 7317, 7398, 7506, 7601, 7697, 7808, 7897,
        7978, 8014, 8029, 8023, 8014, 7984, 7927, 7978, 8050, 8104, 8089, 8065, 8023, 7984, 7927,
        7861, 7778, 7697, 7616, 7521, 7434, 7353, 7278, 7168, 7087, 6971, 6845, 6713, 6597, 6501,
        6397, 6295, 6235, 6169, 6089, 6023, 5957, 5963, 6038, 6068, 6095, 6139, 6190, 6265, 6346,
        6447, 6528, 6603, 6663)
EPL_GAS_4L.y <-c(4237, 4250, 4272, 4294, 4324, 4316, 4355, 4411, 4420, 4433, 4411, 4368, 4303,
        4185, 4016, 3876, 3746, 3598, 3424, 3290, 3211, 3129, 3063, 2933, 2829, 2755, 2668, 2594,
        2550, 2455, 2376, 2324, 2346, 2324, 2250, 2250, 2242, 2133, 2207, 2207, 2207, 2242, 2281,
        2337, 2420, 2537, 2646, 2711, 2829, 2990, 3159, 3224, 3342, 3481, 3620, 3737, 3811, 3885,
        3942, 4046, 4111, 4176)
EPL_GAS_4R.x <-c(6089, 6154, 6229, 6295, 6376, 6477, 6588, 6684, 6809, 6950, 7051, 7168, 7278,
        7374, 7470, 7574, 7676, 7772, 7837, 7903, 7903, 7912, 7897, 7867, 7808, 7691, 7631, 7506,
        7419, 7338, 7066, 6779, 6522, 6199, 6110)
EPL_GAS_4R.y <-c(4976, 4816, 4698, 4603, 4507, 4485, 4472, 4472, 4485, 4529, 4537, 4572, 4581,
        4611, 4655, 4698, 4772, 4829, 4859, 4955, 5094, 5224, 5363, 5468, 5563, 5703, 5759, 5842,
        5876, 5898, 5876, 5716, 5520, 5329, 5146)
glomerular_GAS_4L.x <-c(7808, 7697, 7559, 7449, 7344, 7249, 7141, 7045, 6935, 6860, 6764, 6684,
        6597, 6522, 6462, 6367, 6301, 6229, 6169, 6110, 6059, 6038, 6008, 5984, 5993, 5993, 6008,
        6038, 6068, 6104, 6148, 6184, 6229, 6280, 6346, 6441, 6537, 6648, 6749, 6845, 6950, 7030,
        7147, 7257, 7368, 7449, 7515, 7616, 7691, 7751, 7787, 7888, 7993, 8095, 8065, 8124, 8160,
        8184, 8199, 8184, 8145, 8095, 8118, 8124, 8118, 8110, 8095, 8065, 8014, 7993, 7948, 7897)
glomerular_GAS_4L.y <-c(4611, 4581, 4550, 4529, 4485, 4463, 4455, 4433, 4420, 4398, 4376, 4346,
        4316, 4259, 4216, 4176, 4098, 4037, 3959, 3842, 3724, 3585, 3468, 3329, 3190, 3050, 2968,
        2881, 2785, 2646, 2559, 2485, 2376, 2346, 2272, 2198, 2142, 2120, 2103, 2059, 2046, 2037,
        2024, 2037, 2059, 2059, 2059, 2081, 2133, 2207, 2294, 2390, 2455, 2572, 2690, 2742, 2837,
        2955, 3072, 3320, 3511, 3598, 3737, 3898, 4037, 4155, 4272, 4346, 4420, 4516, 4559, 4603)
glomerular_GAS_4R.x <-c(6648, 6528, 6426, 6280, 6199, 6133, 6074, 6008, 5957, 5927, 5918, 5957,
        6286, 6624, 7015, 7293, 7404, 7515, 7616, 7712, 7778, 7861, 7897, 7963, 8038, 8059, 8059,
        8029, 7984, 7888, 7808, 7697, 7559, 7449, 7344, 7249, 7141, 7045, 6935, 6860, 6764)
glomerular_GAS_4R.y <-c(4346, 4324, 4316, 4346, 4433, 4507, 4624, 4742, 4850, 4955, 5329, 5855,
        6003, 6076, 6076, 6055, 5959, 5929, 5876, 5811, 5716, 5598, 5490, 5372, 5255, 5094, 4933,
        4794, 4690, 4633, 4611, 4581, 4550, 4529, 4485, 4463, 4455, 4433, 4420, 4398, 4376)

# regional polygon coordinates for PBS 4

granular_PBS_4L.x <- c(2352, 2484, 2622, 2741, 2856, 2968, 3084, 3205, 3327, 3456, 3489, 3495,
        3416, 3327, 3222, 3090, 2978, 2889, 2800, 2701, 2639, 2573, 2500, 2395, 2289, 2200, 2102,
        2013, 1917, 1828, 1723, 1634, 1558, 1545, 1551, 1624, 1723, 1818, 1934, 2029, 2128, 2224)
granular_PBS_4L.y <- c(2878, 2892, 2938, 2985, 3026, 3073, 3119, 3128, 3119, 3063, 2925, 2800,
        2712, 2652, 2606, 2596, 2564, 2504, 2425, 2356, 2254, 2153, 2051, 2005, 1972, 1972, 2018,
        2097, 2153, 2231, 2310, 2393, 2448, 2573, 2730, 2823, 2892, 2925, 2938, 2938, 2901, 2901)
granular_PBS_4R.x <- c(1795, 1901, 2029, 2144, 2247, 2369, 2500, 2622, 2741, 2863, 2985, 3090,
        3146, 3146, 3123, 3028, 2896, 2774, 2668, 2573, 2457, 2352, 2247, 2151, 2052, 1963, 1851,
        1729, 1601, 1502, 1423, 1350, 1341, 1373, 1439, 1495, 1551, 1591, 1657, 1723)
granular_PBS_4R.y <- c(4857, 4857, 4857, 4811, 4755, 4709, 4677, 4663, 4663, 4686, 4700, 4677,
        4561, 4427, 4279, 4210, 4187, 4177, 4177, 4177, 4164, 4154, 4099, 4030, 3960, 3914, 3882,
        3835, 3858, 3868, 3914, 4006, 4154, 4288, 4390, 4515, 4631, 4732, 4811, 4848)
EPL_PBS_4L.x <- c(2695, 2807, 2945, 3090, 3228, 3341, 3456, 3594, 3657, 3706, 3706, 3617, 3528,
        3406, 3294, 3179, 3041, 2945, 2863, 2800, 2718, 2639, 2517, 2395, 2273, 2144, 1996, 1874,
        1795, 1690, 1601, 1495, 1413, 1397, 1364, 1390, 1423, 1462, 1535, 1657, 1785, 1901, 1990,
        2112, 2200, 2289, 2395, 2500, 2589)
EPL_PBS_4L.y <- c(3174, 3220, 3267, 3299, 3313, 3299, 3290, 3220, 3086, 2925, 2777, 2652, 2541,
        2472, 2393, 2324, 2300, 2231, 2083, 1949, 1880, 1787, 1764, 1732, 1709, 1709, 1741, 1810,
        1880, 1972, 2074, 2162, 2245, 2402, 2541, 2689, 2800, 2915, 2994, 3049, 3096, 3096, 3110,
        3119, 3119, 3119, 3096, 3110, 3142)
EPL_PBS_4R.x <- c(2296, 2395, 2500, 2629, 2741, 2833, 2929, 3067, 3196, 3294, 3327, 3373, 3311,
        3261, 3196, 3116, 3001, 2912, 2790, 2668, 2556, 2451, 2329, 2207, 2085, 1980, 1858, 1713,
        1545, 1406, 1268, 1196, 1146, 1123, 1140, 1202, 1252, 1285, 1373, 1462, 1574, 1696, 1812,
        1934, 2039, 2151, 2224)
EPL_PBS_4R.y <- c(4959, 4936, 4880, 4894, 4894, 4894, 4848, 4848, 4811, 4732, 4575, 4436, 4335,
        4233, 4131, 4085, 4030, 4006, 3983, 3969, 3951, 3928, 3868, 3789, 3720, 3678, 3678, 3618,
        3641, 3687, 3757, 3858, 3960, 4108, 4279, 4427, 4607, 4755, 4871, 4936, 4996, 5028, 5098,
        5098, 5065, 5028, 5005)
glomerular_PBS_4L.x <- c(1874, 2013, 2168, 2306, 2467, 2612, 2751, 2863, 3028, 3179, 3327, 3446,
        3551, 3667, 3762, 3868, 3950, 3983, 3983, 3927, 3835, 3746, 3627, 3522, 3400, 3278, 3163,
```

```
        3074, 2945, 2823, 2678, 2540, 2385, 2257, 2095, 1940, 1769, 1640, 1545, 1469, 1406, 1350,
        1285, 1219, 1202, 1186, 1156, 1219, 1285, 1357, 1456, 1518, 1640, 1713, 1795)
glomerular_PBS_4L.y <- c(3322, 3313, 3299, 3299, 3313, 3336, 3359, 3424, 3484, 3530, 3530, 3507,
        3470, 3424, 3313, 3211, 3073, 2901, 2767, 2606, 2472, 2379, 2254, 2185, 2083, 1995, 1935,
        1801, 1662, 1607, 1552, 1491, 1459, 1459, 1450, 1505, 1584, 1662, 1755, 1834, 1926, 2051,
        2162, 2300, 2448, 2573, 2744, 2823, 2901, 3073, 3142, 3220, 3267, 3299, 3322)
glomerular_PBS_4R.x <- c(1364, 1301, 1228, 1140, 1064, 1018, 985, 985, 991, 1008, 1041, 1113,
        1186, 1268, 1350, 1479, 1624, 1713, 1818, 1940, 2102, 2224, 2362, 2484, 2579, 2711, 2807,
        2952, 3084, 3179, 3285, 3373, 3472, 3535, 3601, 3667, 3683, 3667, 3611, 3505, 3383, 3285,
        3179, 3057, 2906, 2774, 2685, 2563, 2474, 2362, 2273, 2135, 2006, 1858, 1690, 1574, 1485)
glomerular_PBS_4R.y <- c(3470, 3572, 3655, 3743, 3868, 3993, 4164, 4335, 4483, 4631, 4732, 4825,
        4982, 5051, 5153, 5190, 5269, 5315, 5338, 5356, 5315, 5245, 5199, 5144, 5107, 5107, 5098,
        5107, 5084, 5074, 5065, 4996, 4936, 4880, 4792, 4709, 4575, 4460, 4312, 4201, 4053, 3969,
        3905, 3845, 3803, 3789, 3766, 3734, 3664, 3586, 3507, 3470, 3447, 3415, 3415, 3405, 3447)


# merge all microglia objects and export layer info

MERFISH_merged <- merge(MERFISH_PBS_1_microglia, y = c(MERFISH_GAS_1_microglia,
       MERFISH_PBS_2_microglia, MERFISH_GAS_2_microglia, MERFISH_PBS_3_microglia,
       MERFISH_GAS_3_microglia, MERFISH_PBS_4_microglia, MERFISH_GAS_4_microglia))
Idents(object=MERFISH_merged) <- MERFISH_merged@meta.data$layer
plot.xy.cluster(MERFISH_merged)

LayerIdentity <- MERFISH_merged[["layer"]]
MERFISH_microglia_layers <- AddMetaData(MERFISH_all, metadata = LayerIdentity,
       col.name = "microglia_layer")


# quantification of microglial gene expression by glomerular vs granular layer ----

Idents(object=MERFISH_microglia) <- MERFISH_microglia@meta.data$microglia_layer
MERFISH_microglia_glomerular <- subset(x = MERFISH_microglia, idents =
       c("glomerular"),invert = FALSE)
MERFISH_microglia_granular <- subset(x = MERFISH_microglia, idents =
       c("granular"),invert = FALSE)

microglial.genes <- c("P2ry12", "Gpr34", "Cd74", "Axl", "Ifi30")

Idents(object=MERFISH_microglia_glomerular) <-
       MERFISH_microglia_glomerular@meta.data$sample
Idents(object=MERFISH_microglia_granular) <-
       MERFISH_microglia_granular@meta.data$sample

MERFISH_microglia_Original_LogNormalized_GeneExpression_byLayer_Glomerular <-
       AverageExpression(MERFISH_microglia_glomerular, features = microglial.genes,
       slot = "counts")

MERFISH_microglia_Original_LogNormalized_GeneExpression_byLayer_Granular <-
       AverageExpression(MERFISH_microglia_granular, features = microglial.genes, slot
       = "counts")

# spatial feature plots

Idents(object=MERFISH_microglia) <- MERFISH_microglia@meta.data$sample

MERFISH_microglia_GAS_1 <- subset(x = MERFISH_microglia, idents = c("GAS_1"),invert =
       FALSE)
MERFISH_microglia_PBS_1 <- subset(x = MERFISH_microglia, idents = c("PBS_1"),invert =
       FALSE)
```

```r
library(scales)

plot.xy.gene.minimal <- function(object, gene){

object.df <- data.frame(object$center.x, object$center.y, FetchData(object = object,
       vars = gene, slot = "counts"))
x <- object$center.x
y <- object$center.y
Expression <- object.df[,3]
mid <- mean(Expression)
ggplot(object.df, aes(x, y, colour = Expression)) + geom_point(size = 2.5) +
       ggtitle(paste(gene)) + theme_linedraw() + scale_color_gradient(trans = "log",
       limits = c(.001,.2), na.value = "gray90", low = "light gray", high = "#1e0bfe",
       oob=squish)+ theme(plot.title = element_text(face = "italic", size=24, vjust =
       0, hjust = 0.5), axis.title = element_blank(), panel.grid = element_blank(),
       panel.border = element_rect(color = "white"), axis.ticks = element_blank(),
       axis.text = element_blank()) + NoLegend()
}      # update limits as needed, depending on expression range

plot.xy.gene.minimal(MERFISH_microglia_PBS_1, "P2ry12")# limits = c(.001,.2)
plot.xy.gene.minimal(MERFISH_microglia_GAS_1, "P2ry12")# limits = c(.001,.2)

plot.xy.gene.minimal(MERFISH_microglia_PBS_1, "Gpr34")# limits = c(.001,.06)
plot.xy.gene.minimal(MERFISH_microglia_GAS_1, "Gpr34")# limits = c(.001,.06)

plot.xy.gene.minimal(MERFISH_microglia_PBS_1, "Cd74")# limits = c(.01,.05)
plot.xy.gene.minimal(MERFISH_microglia_GAS_1, "Cd74")# limits = c(.01,.05)

plot.xy.gene.minimal(MERFISH_microglia_PBS_1, "Ifi30")# limits = c(.001,.05)
plot.xy.gene.minimal(MERFISH_microglia_GAS_1, "Ifi30")# limits = c(.001,.05)

plot.xy.gene.minimal(MERFISH_microglia_PBS_1, "Axl")# limits = c(.001,.05)
plot.xy.gene.minimal(MERFISH_microglia_GAS_1, "Axl")# limits = c(.001,.05)
```

#### KEY RESOURCES TABLE

| REAGENT or RESOURCE | SOURCE | IDENTIFIER |
| --- | --- | --- |
| **Antibodies** | | |
| anti-mouse Caveolin 1 | Abcam | Cat: ab18199 |
| anti-mouse CD4 | BD Pharmingen | Cat: 553727 |
| anti-mouse CD68 | Abcam | Cat: ab53444 |
| anti-mouse GLUT1 | Millipore Sigma | Cat: 400060 |
| anti-mouse Iba1 | WAKO | Cat: 016-20001 |
| anti-mouse Mfsd2a (clone J9590) | Lab of Chengua Gu | Pfau et al. (2021)[94] |
| anti-mouse Podocalyxin | R&D Systems | Cat: AF1556 |
| Donkey IgG (H+L) Highly Cross-Adsorbed Secondary Antibody against rat, Alexa Fluor 594, 647 | Invitrogen | Cat: A-21209, A-78947 |
| Donkey IgG (H+L) Highly Cross-Adsorbed Secondary Antibody against rabbit, Alexa Fluor 488, 647 | Invitrogen | Cat: A-21206, A-31573 |
| Donkey IgG (H+L) Highly Cross-Adsorbed Secondary Antibody against goat, Alexa Fluor 647 | Invitrogen | Cat: A-21447 |
| anti-mouse CD31 | BioLegend | Cat: 102502 |
| anti-mouse CCL5 PE-Cy7 | BioLegend | Cat: 149105 |
| anti-mouse CD31 FITC | BD Biosciences | Cat: 553372 |
| anti-mouse CD4 BV605 | BD Biosciences | Cat: 563151 |
| anti-mouse/human CD11b BV421 | BioLegend | Cat: 101235 |
| anti-mouse CD11b PerCP-Cy5.5 | BioLegend | Cat: 101227 |
| anti-mouse/human CD44 BV421 | BioLegend | Cat: 103040 |
| anti-mouse CD45 AF700 | BioLegend | Cat: 103127 |
| anti-mouse CD45 BUV395 | BD Biosciences | Cat: 564279 |
| anti-mouse CD45 BV421 | BD Biosciences | Cat: 563890 |
| anti-mouse CD62L PE-Cy7 | BioLegend | Cat: 104417 |
| anti-mouse CD68 APC | BioLegend | Cat: 137007 |
| anti-mouse CD74 BV711 | BD Biosciences | Cat: 740748 |
| anti-mouse CX3CR1 BV786 | BioLegend | Cat: 149029 |
| anti-mouse GM-CSF FITC | BioLegend | Cat: 505403 |
| anti-mouse IFNγ APC | BD Biosciences | Cat: 554413 |
| anti-mouse IL-17A PE | BD Biosciences | Cat: 559502 |
| anti-mouse Ki67 BV605 | BioLegend | Cat: 652413 |
| anti-mouse MHC II I-A/I-E APC-Cy7 | BioLegend | Cat: 107627 |
| anti-mouse P2RY12 PE | BioLegend | Cat: 848003 |
| anti-mouse TMEM119 AF488 | Abcam | Cat: ab225497 |
| anti-mouse TNF BV421 | BioLegend | Cat: 506327 |
| anti-mouse CD16/CD32 (Fc block) | BD Biosciences | Cat: 553141 |

| | | |
|---|---|---|
| anti-mouse IL-17A monoclonal antibody, InVivoMAb (clone 17F3) | Bio X Cell | Cat: BE0173 |
| mouse IgG1, κ isotype control monoclonal antibody (clone MOPC-21) | Bio X Cell | Cat: BE0083 |
| anti-Digoxigenin-AP, Fab fragments, from sheep | Roche | Cat: 11093274910 |
| anti-Digoxigenin-POD, Fab fragments, from sheep | Roche | Cat: 11207733910 |
| **Chemicals, peptides, and recombinant proteins** | | |
| Isoflurane | Covetrus | Cat: 029405 |
| Ribonucleic acid from baker's yeast | Sigma-Aldrich | Cat: R6750 |
| Salmon sperm DNA solution | ThermoFisher | Cat: 15632011 |
| 4-Nitro blue tetrazolium chloride, solution (NBT) | Roche | Cat: 11383213001 |
| BCIP, 4-toluidine salt | Roche | Cat: 11383221001 |
| Ethylenediaminetetraacetic acid (EDTA) solution (0.5 M) | ThermoFisher | Cat: 1861274 |
| Paraformaldehyde, 32% aqueous solution | Electron Microscopy Sciences | Cat: 15714 |
| Triton X-100 | VWR | Cat: PAH5141 |
| Saline sodium citrate (SSC) | ThermoFisher | Cat: AM9765 |
| Acrylamide/Bis, 40% solution, 19:1 | Bio-Rad | Cat: 1610144 |
| Ammonium persulfate | Millipore Sigma | Cat: 09913 |
| SDS, 10% solution | ThermoFisher | Cat: AM9823 |
| Proteinase K | New England BioLabs | Cat: P8107S |
| N,N,N′,N′-Tetramethyl ethylenediamine (TEMED) | Millipore Sigma | Cat: T7024 |
| (Z)-4-Hydroxytamoxifen | Millipore Sigma | Cat: H7904 |
| Paraformaldehyde | Fisher Scientific | Cat: AC416785000 |
| Cell lysis buffer (10X) | Abcam | Cat: ab152163 |
| Gelatin | Sigma Aldrich | Cat: G1319 |
| Poly-D-lysine hydrobromide | Sigma Aldrich | Cat: P6407 |
| Collagen IV | Corning | Cat: CB-40233 |
| recombinant mouse IL-1β | R&D Systems | Cat: 401-ML |
| recombinant mouse TNF | R&D Systems | Cat: 410-MT |
| recombinant mouse IFNγ | R&D Systems | Cat: 485-MI-100 |
| recombinant mouse IL-17A | R&D Systems | Cat: 7956ML025 |
| recombinant mouse GM-CSF | R&D Systems | Cat: 415-ML-010 |
| Albumin AlexaFluor 647 from bovine serum | ThermoFisher | Cat: A34785 |
| **Critical commercial assays** | | |
| Neural Tissue Dissociation Kit (P) | Miltenyi | Cat: 130-092-628 |
| Myelin removal beads | Miltenyi | Cat: 130-096-733 |
| LS columns | Miltenyi | Cat: 130-042-401 |
| C tubes | Miltenyi | Cat: 130-093-237 |

| | | |
|---|---|---|
| octoMACS with heaters | Miltenyi | Cat: 130-096-427 |
| Digoxigenin RNA Labeling Kit (SP6/T7) | Roche | Cat: 11175025910 |
| Microspin G-50 Columns | GE Healthcare | Cat: 27533001 |
| TSA Plus Cyanine 3 system | Akoya Bio | Cat: NEL744001KT |
| TSA Plus Fluorescein system | Akoya Bio | Cat: NEL741001KT |
| Cell Stimulation Cocktail (plus protein transport inhibitors) | eBiosciences | Cat: 00-4975-03 |
| Intracellular Fixation & Permeabilization Buffer Set | BD Biosciences | Cat: 554722 |
| ZTheta 96 Well Array Station | Applied BioPhysics | biophysics.com/ztheta.php |
| 96 Well Electrode Array Plate | Applied BioPhysics | Cat: 96W20idf |
| Halt Protease and Phosphatase Inhibitor Cocktail | ThermoFisher | Cat: 78440 |
| Pierce BCA Protein Assay Kit | ThermoFisher | Cat: 23225 |
| Slide-A-Lyzer Dialysis Cassettes, 2K MWCO, 0.5 mL | ThermoFisher | Cat: 66205 |
| Mouse ProcartaPlex Mix & Match 12-plex | ThermoFisher | Cat: PPX-12-MXEPUF3 |
| Human 45-Plex ProcartaPlex Panel 1, Cytokine/Chemokine/Growth Factor | Invitrogen | Cat: EPX260-26088-901 |
| **Deposited data** | | |
| Olfactory bulb scRNAseq data | This paper | GEO: GSE221724 |
| Nasal lymphoid tissue scRNAseq data | This paper | GEO: GSE221724 |
| Olfactory bulb MERFISH data | This paper | GEO: GSE221106 |
| *In vitro* microglia bulk RNAseq data | This paper | GEO: GSE221715 |
| Experimental autoimmune encephalomyelitis data | Shahriar et al., *bioRxiv*, 2022[36] | GEO: GSE210776 |
| **Experimental Models: Cell lines** | | |
| Bacteria: *Streptococcus pyogenes* (2W) | Lab of Pat Cleary | Dileepan et al. (2011)[20] |
| Primary mouse brain microvascular endothelial cells (BMEC) | Cell Biologics | Cat: C57-6023 |
| Primary human umbilical vein endothelial cells (HUVEC) | ATCC | Cat: PCS-100-010 |
| **Experimental Models: Organisms/strains** | | |
| Mouse: C57BL/6J | The Jackson Laboratory | Cat: 000664 |
| Mouse: RORg (B6.129P2(Cg)-*Rorctm2Litt*/J) | The Jackson Laboratory | Cat: 007572 |
| Mouse: Csf2$^{flox/flox}$ | Lab of Bogoljub Ciric | Louis et al. (2020)[54] |
| Mouse: CD4-Cre ERT2 (B6(129X1)-Tg(Cd4-cre/ERT2)11Gnri/J) | The Jackson Laboratory | Cat: 022356 |
| Mouse: CX3CR1-GFP | Lab of Wassim Elyaman | Jung et al. (2000)[40] |
| Mouse: TMEM119-tdTomato | Lab of Wassim Elyaman | Ruan et al. (2020)[41] |
| **Recombinant DNA** | | |

| | | |
|---|---|---|
| *Mus musculus Itih5* (pCMV-SPORT6) glycerol stock | Transomic Technologies | Cat: BC043314-seq |
| *Mus musculus Itm2a* (pCMV-SPORT6) glycerol stock | Transomic Technologies | Cat: BC031420-seq |
| **Software and algorithms** | | |
| R v4.0.2 | R Core Team (2020) | r-project.org |
| R Studio v1.3.959 | R Studio Team (2020) | rstudio.com |
| Seurat v4.0.2 | Hao et al. (2021)[98] | github.com/satijalab/seurat |
| Harmony package, v1 | Korsunsky et al. (2019) [99] | portals.broadinstitute.org/harmony |
| Gene set enrichment analysis (GSEA) v4.1.0 | Subramanian et al. (2005)[31] | gsea-msigdb.org/gsea |
| BBrowser 3 | BioTuring | bioturing.com |
| ImageJ (FIJI) | Schindelin et al. (2012)[102] | imagej.net/software/fiji |
| Prism 9 | GraphPad | graphpad.com |
| MERlin v0.1.12 | Vizgen | github.com/ZhuangLab/MERlin |
| ComBat v1 | Zhang et al. (2020)[101] | github.com/zhangyuqing/ComBat-seq |
| MobileFish: Record XY mouse coordinates on an uploaded image | MobileFish | mobilefish.com |
| FlowJo v10.5 | FlowJo, LLC | flowjo.com |
| Illustrator v25.4.1 | Adobe | adobe.com |
| **Other** | | |
| Trypicase Soy Agar with 5% Sheep Blood | Fisher Scientific | Cat: B21239 |
| Todd-Hewitt Broth | Bacto | Cat: 90003-430 |
| Neopeptone | Bacto | Cat: 90000-268 |
| DietGel 76A | Clear$H_2O$ | Cat: 72-07-5022 |
| Hanks Balanced Salt Solution (HBSS) | ThermoFisher | Cat: 14175095 |
| DRAQ5 | BioLegend | Cat: 424101 |
| anti-rat Ig κ/Negative compensation beads | BD Biosciences | Cat: 552844 |
| UltraComp eBeads Plus Compensation Beads | ThermoFisher | Cat: 01-3333-41 |
| Tissue-Plus O.C.T. | Fisher | Cat. 4585 |
| Bovine serum albumin | Sigma Aldrich | Cat: A9647 |
| Vectashield with DAPI | Vector Labs | Cat: H-1200 |
| Fetal bovine serum | Cytiva | Cat: SH30071.03 |
| Normal goat serum | Abcam | Cat: ab7481 |
| HybriSlip Hybridization Covers | Grace Bio-Labs | Cat: 726024 |
| Glycergel mounting medium | Dako | Cat: C0563 |
| Blocking reagent | Roche | Cat: 11096176001 |

| | | |
|---|---|---|
| Cell lysis buffer (2X) | Abcam | Cat: ab152163 |
| Slide-A-Lyzer Dialysis Cassettes, 2K MWCO | ThermoFisher | Cat: 66205 |
| Dulbecco's Modified Eagle Medium (DMEM) | Genesee | Cat: 25-500 |
| RPMI | Corning | Cat: 15-040-CV |
| Percoll | Cytiva | Cat: GE17-0891-01 |
| Propidium iodide | ThermoFisher | Cat: P3566 |
| Live/Dead Fixable Aqua Dead Cell Stain Kit | ThermoFisher | Cat: L34965 |
| Fixable viability dye 780 | ThermoFisher | Cat: 65-0865-14 |
| Intracellular Fixation & Permeabilization kit | eBiosciences | Cat: 88-8824-00 |
| Complete endothelial cell medium kit | Cell Biologics | Cat: M1168 |
| Transparent 3.0 µm PET membranes for 24-well plates | Corning | Cat: 353096 |
| Antibody dilution buffers | Bio X Cell | Cat: IP0070, cat. IP0065 |
| Corn oil | Millipore Sigma | Cat: C8267 |
